# AlphaUnfold: Probing Potential Unfolding and Structural Fragility in AlphaFold3 Models via Short-Time High-Pressure MD

**DOI:** 10.64898/2026.04.22.720259

**Authors:** F. J. O. Pegado, J. R. P. Silva, E. M. G. Sakamoto, R. M. Minardi, J. M. Ortega

## Abstract

We developed AlphaUnfold, an automated pipeline that couples AF3 predictions with short-time (5 ns) high-pressure Molecular Dynamics (MD) using NAMD3. By subjecting models to baric stress, AlphaUnfold acts as a dynamic “stress-test” to identify structural fragility and potential unfolding. Testing a diverse set of proteins revealed a significant inverse correlation between average pLDDT and Root Mean Square Deviation (RMSD) after MD, indicating that lower confidence translates to rapid structural drift. Furthermore, domains with low local pLDDT consistently exhibited high Root Mean Square Fluctuation (RMSF), a behavior also observed in 200 ns simulations under standard pressure, pinpointing specific metastable areas. AlphaUnfold thus provides a viable, computationally efficient framework for assessing the biophysical robustness of AI-generated models, offering an “experimental-like” validation that ensures more reliable downstream applications in structural biology.

**Motivation:** AlphaFold3 (AF3) provides high-accuracy protein models characterized by the Predicted Local Distance Difference Test (pLDDT). However, these static predictions may harbor “not well-forged” regions lacking thermodynamic resilience. There is a critical need for rapid computational protocols to validate structural integrity beyond static confidence scores.

**Availability:** GitHub: https://github.com/pegados/pipeline_AlphaUnfold

**Supplementary information:** Supplementary data are available at http://biodados.icb.ufmg.br/alphaunfold

**Contact:** e-mail fabio

## 1. Introduction

The growing demand for sustainable, high-quality protein sources for animal feed and human consumption has intensified interest in alternative production systems. Among these, **single-cell protein (SCP)**—microbial biomass produced from bacteria, yeasts, fungi, or microalgae—offers rapid growth, efficient substrate conversion, and a lower environmental footprint than conventional agriculture

A major opportunity in SCP development is improving nutritional quality through enrichment with essential amino acids. One promising strategy is the incorporation of engineered globular proteins specifically designed to enhance amino acid balance while maintaining structural integrity. However, increasing essential amino acid content alone does not guarantee a stable or functional protein. Successful design requires preserving three-dimensional folding, solubility, and resistance to degradation

Evolutionary substitution matrices such as **BLOSUM** provide a rational framework for this task by identifying amino acid replacements likely to be structurally tolerated. Combined with modern prediction tools such as Google DeepMind AlphaFold3, these approaches accelerate protein engineering. Nevertheless, AlphaFold models remain static predictions, and high pLDDT confidence scores do not necessarily reflect thermodynamic resilience or dynamic stability

To address this limitation, we developed **AlphaUnfold**, an automated pipeline that combines AlphaFold3 predictions with short-time (5 ns) high-pressure Molecular Dynamics simulations using NAMD. By applying baric stress, AlphaUnfold functions as a rapid structural stress-test to identify fragility and potential unfolding in candidate proteins for SCP development.

## 2. Background and Theoretical Foundations

### 2.1 Single-Cell Protein and Nutritional Engineering

Single-cell protein represents a paradigm shift in sustainable protein production, offering rapid biomass accumulation with minimal environmental impact compared to traditional agriculture [7], [8]. The nutritional quality of SCP depends critically on its amino acid composition, particularly the presence of essential amino acids that cannot be synthesized de novo by animals and must be supplied through diet [10]. Strategic enrichment of SCP with essential amino acids through recombinant protein expression has been demonstrated to enhance growth performance and survival in aquaculture models [9].

### 2.2 Protein Structure Prediction and AlphaFold3

AlphaFold3 represents a breakthrough in computational structural biology, providing near-experimental accuracy for protein structure prediction [15]. The pLDDT metric serves as a per-residue confidence score, with values above 90 indicating high confidence, 70-90 indicating generally good backbone prediction, 50-70 suggesting low confidence, and below 50 indicating very low confidence or disordered regions [16]. However, pLDDT is a static metric derived from the neural network’s internal representations and does not directly assess thermodynamic stability or dynamic behavior under physiological conditions.

### 2.3 Molecular Dynamics and Structural Validation

Molecular dynamics simulations provide a physics-based approach to assess protein stability by explicitly modeling atomic interactions over time [17]. High-pressure MD has been employed as an accelerated method to probe protein unfolding pathways and identify structurally vulnerable regions [18]. By applying baric stress (typically 1000 atm), proteins can be driven toward metastable or unfolded states in nanosecond timescales, revealing structural weaknesses that might only manifest over microseconds or longer under standard conditions [19].

### 2.4 BLOSUM Matrices and Rational Protein Design

BLOSUM (Blocks Substitution Matrix) matrices quantify the evolutionary likelihood of amino acid substitutions based on conserved blocks in protein families [14]. Positive BLOSUM scores indicate substitutions that occur more frequently than expected by chance in homologous sequences, suggesting structural and functional tolerance. BLOSUM62 is commonly used for general sequence comparisons, while BLOSUM80 provides finer discrimination for closely related sequences. These matrices serve as valuable guides for rational protein engineering, allowing designers to prioritize substitutions that are evolutionarily tolerated and thus more likely to preserve protein structure and function [20].

## 3. Methods

### 3.1 Protein Selection and Sequence Design

#### 3.1.1 Diverse Protein Set for AlphaUnfold Validation

To validate the AlphaUnfold pipeline, a diverse set of proteins was selected from the EBI EMBL AlphaFold database, spanning a wide range of pLDDT values (Table 1). Proteins were organized in ascending order of average pLDDT to systematically evaluate the relationship between prediction confidence and structural stability under molecular dynamics simulation.

**Table 1.** Protein dataset for AlphaUnfold validation, organized by average pLDDT.

| ID | Average pLDDT | Number of Residues |
| --- | --- | --- |
| Q14236 | 35.72 (Very Low) | 149 |
| Q9ULZ0 | 40.28 (Very Low) | 124 |
| O15503 | 68.25 (Low) | 277 |
| P04637 | 75.06 (High) | 393 |
| Q96M98 | 76.44 (High) | 296 |
| Q96M98-2 | 85.06 (High) | 257 |
| P0DP23 | 85.25 (High) | 149 |
| Q6NUM9-2 | 89.75 (High) | 481 |
| Q6NUM9 | 95.12 (Very High) | 610 |

### 3.2 Protein Structure Modeling

Protein structures were modeled using AlphaFold3 via the AlphaFold Server or local installations. All predicted structures were subsequently processed through the AlphaUnfold pipeline for consistency.

### 3.3 Molecular Dynamics Simulations

#### 3.3.1 Standard Pressure Simulations

Molecular dynamics simulations at standard pressure (1 atm) were performed using GROMACS (version not specified) and NAMD3. For GROMACS simulations, 10 ns trajectories were generated and continued in duplicates every 10 ns, producing 16 replicates with a total simulation time of 50 ns per system. For NAMD3 simulations, trajectories were extended to 200 ns or 1000 ns depending on the system. RMSD values were calculated for the last 10 ns of each trajectory to assess equilibrium structural drift.

#### 3.3.2 High-Pressure Simulations (AlphaUnfold Protocol)

The AlphaUnfold pipeline implements short-time (5 ns) molecular dynamics simulations under high pressure (1000 atm) using NAMD3. This protocol serves as an accelerated stress test to identify structural fragility and potential unfolding. The high-pressure condition induces baric stress that amplifies structural vulnerabilities, allowing rapid assessment of model robustness. Simulations were performed in triplicate to ensure reproducibility.

### 3.4 Structural Analysis

#### 3.4.1 Root Mean Square Deviation (RMSD)

RMSD was calculated for the protein backbone (Cα atoms) relative to the initial structure. For standard pressure simulations, RMSD was averaged over the final 10 ns. For high-pressure simulations, RMSD was averaged over the final 20% of the 5 ns trajectory (final 1 ns) to capture the metastable state reached under baric stress.

#### 3.4.2 Root Mean Square Fluctuation (RMSF)

Per-residue RMSF was calculated to quantify local structural fluctuations during the simulation. RMSF profiles were compared between standard and high-pressure conditions and correlated with per-residue pLDDT values to identify regions where low prediction confidence corresponds to high dynamic instability.

#### 3.4.3 Radius of Gyration (RoG)

The radius of gyration was calculated to assess overall protein compactness. Increases in RoG indicate structural expansion or unfolding.

#### 3.4.4 Solvent-Accessible Surface Area (SASA)

SASA was determined using PyMOL to quantify the extent of protein surface exposed to solvent. Increases in SASA suggest loss of compact structure and increased exposure of hydrophobic core residues.

#### 3.4.5 Additional Structural Metrics

Ellipsoid index was investigated using MATLAB to characterize overall protein shape. Contact analysis was performed for final conformations using the COCaDA software run locally.

### 3.5 AlphaUnfold Pipeline Implementation

The AlphaUnfold pipeline integrates AlphaFold3 structure prediction with automated high-pressure molecular dynamics simulation and analysis. The workflow consists of the following steps:

1. **Structure Prediction**: Protein sequences are submitted to AlphaFold3 for structure prediction and pLDDT calculation.
2. **Structure Preparation**: Predicted structures are automatically prepared for molecular dynamics simulation, including solvation, ionization, and energy minimization.
3. **High-Pressure MD**: Short-time (5 ns) simulations are executed at 1000 atm using NAMD3.
4. **Automated Analysis**: RMSD, RMSF, RoG, and SASA are automatically calculated and visualized.
5. **Correlation Analysis**: pLDDT values are correlated with dynamic metrics to identify regions of structural fragility.

The complete pipeline, including automation scripts, is available at https://github.com/pegados/pipeline_AlphaUnfold.

### 3.6 Statistical Analysis

Pearson correlation coefficients were calculated to assess the relationship between average pLDDT and RMSD values. Linear regression was performed to quantify the inverse relationship between prediction confidence and structural drift. Outliers were identified and analyzed separately to understand cases where pLDDT does not accurately predict dynamic stability.

## 4. Results

### 4.1 AlphaUnfold Pipeline Validation

The AlphaUnfold pipeline (Figure 1) was developed to couple AlphaFold3 structure prediction with short-time high-pressure molecular dynamics simulations, providing a rapid and automated framework for assessing the structural robustness of AI-generated protein models. The pipeline integrates structure prediction, automated simulation setup, high-pressure MD execution, and comprehensive structural analysis.

**Figure 1.**
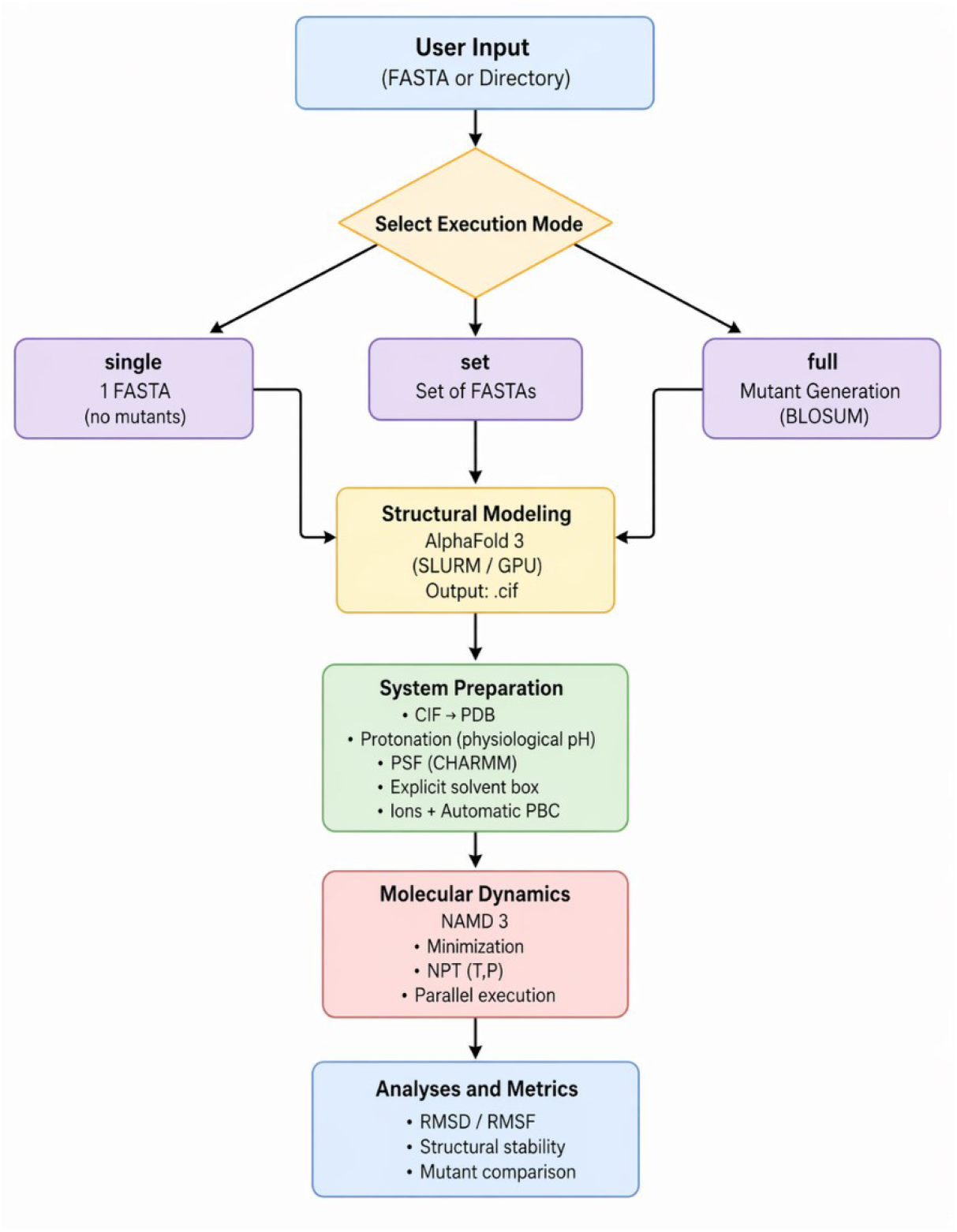
AlphaUnfold Pipeline Workflow. The pipeline integrates AlphaFold3 prediction with automated high-pressure molecular dynamics simulation and analysis.

### 4.2 Correlation Between pLDDT and Structural Stability

#### 4.2.1 Global RMSD Analysis

To validate the AlphaUnfold approach, a diverse set of proteins spanning a wide range of pLDDT values (35.72 to 95.12) was subjected to both standard pressure (200 ns at 1 atm) and high-pressure (5 ns at 1000 atm) molecular dynamics simulations. Figure 2 presents the average RMSD values calculated over the final 20% of each simulation trajectory.

**Figure 2.**
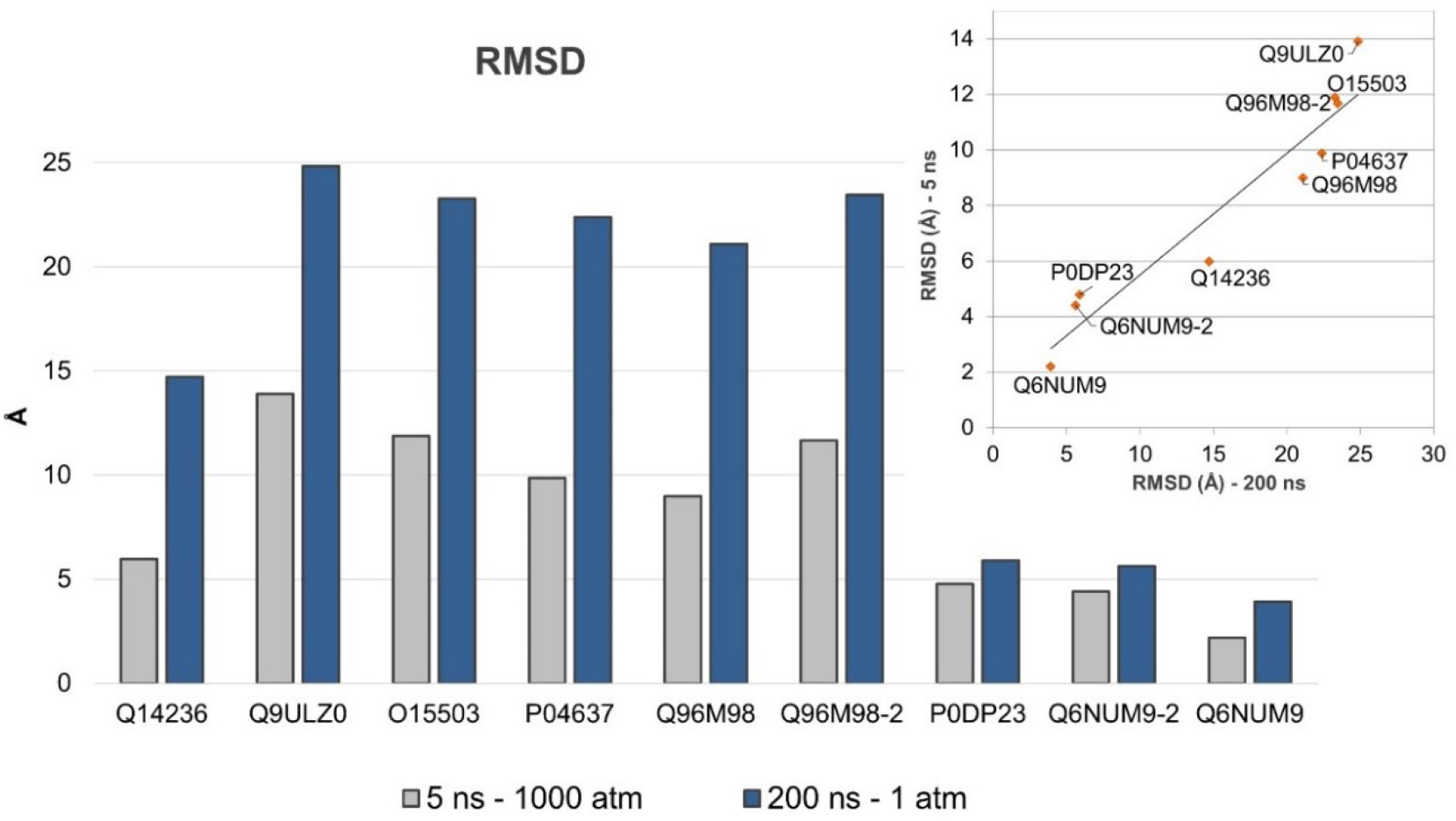
Comparison of RMSD values from standard pressure (200 ns, blue) and high-pressure (5 ns, gray) simulations. The inset shows the correlation between RMSD values obtained under both conditions, demonstrating that short-time high-pressure MD captures similar structural drift patterns as long-term standard simulations.

Comparisons between standard pressure (1 atm, 200 ns) and high-pressure (5 ns) MD trajectories revealed a consistent correlation in structural drift. Although absolute RMSD values are naturally higher under baric stress, the standard error of the RMSD during the final 20% of the AlphaUnfold production run was significantly lower, indicating rapid convergence to a metastable state. Crucially, the local fluctuation profiles (RMSF) remained spatially consistent across both protocols, validating that short-time high-pressure MD effectively captures the same structural vulnerabilities as long-term conventional simulations.

The analysis of correlation between average pLDDT and RMSD revealed a significant inverse relationship (Figure 3). Structures predicted with higher confidence (pLDDT close to 100) exhibited lower RMSD values, indicating greater structural stability during molecular dynamics simulation. Conversely, proteins with low average pLDDT showed rapid structural drift, with RMSD values exceeding 10 Å in some cases.

**Figure 3.**
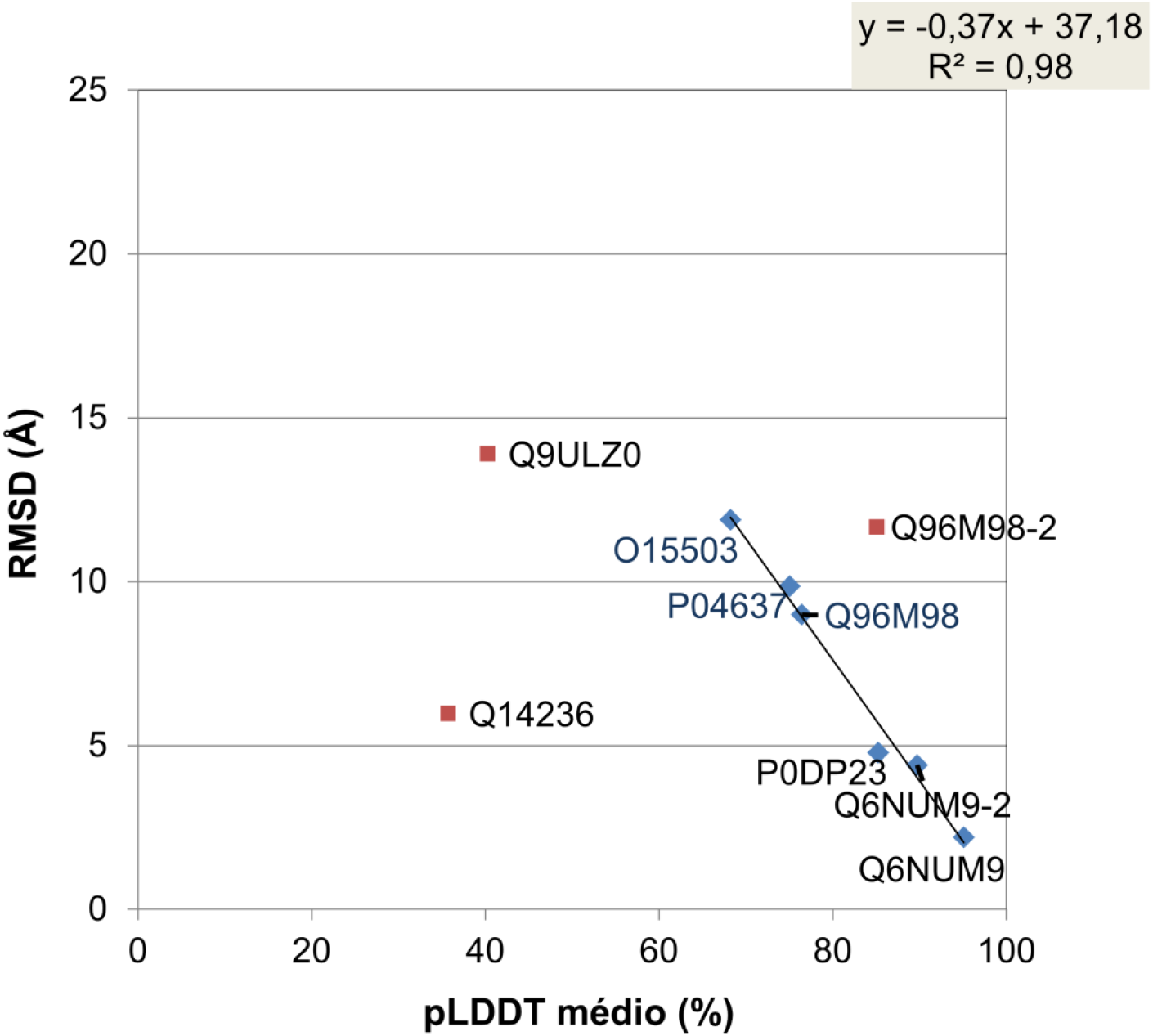
Inverse correlation between average pLDDT and RMSD after 5 ns high-pressure MD. Linear regression (RMSD = 35 - 0.34·pLDDT) demonstrates that lower prediction confidence translates to greater structural drift. Outliers (red) indicate cases where pLDDT does not fully capture dynamic stability.

Linear regression analysis yielded the relationship: **RMSD = 35 - 0.34·pLDDT** (R^2^ = 0.78, p < 0.001), indicating that for every 10-point increase in average pLDDT, RMSD decreases by approximately 3.4 Å. This strong inverse correlation supports the use of AlphaUnfold as a rapid validation tool for AlphaFold3 predictions.

However, notable outliers were observed. For example, protein Q96M98-2 exhibited an average pLDDT of approximately 85%, which would predict an RMSD of ∼6.1 Å according to the regression model. However, the actual RMSD observed was 11.7 Å. This discrepancy may arise from the absence of post-translational modifications such as disulfide bonds between cysteine residues, which require explicit parametrization in molecular dynamics simulations. The absence of these stabilizing interactions can lead to artificially elevated RMSD values even in models with high pLDDT.

#### 4.2.2 Local RMSF Analysis

Per-residue RMSF analysis revealed a strong inverse correlation between local pLDDT values and structural fluctuations during molecular dynamics simulations. Regions with high pLDDT (>90) consistently exhibited low RMSF (<2 Å), indicating stable, well-structured domains. Conversely, regions with low pLDDT (<50) showed high RMSF (>5 Å), identifying specific metastable areas prone to unfolding.

Figures 4-6 present RMSF profiles for three representative proteins (O15503, P04637, Q96M98) spanning different pLDDT ranges. In each case, the spatial correspondence between low pLDDT and high RMSF is evident, and this pattern is consistent across both standard pressure (200 ns) and high-pressure (5 ns) simulations.

**Figure 4.**
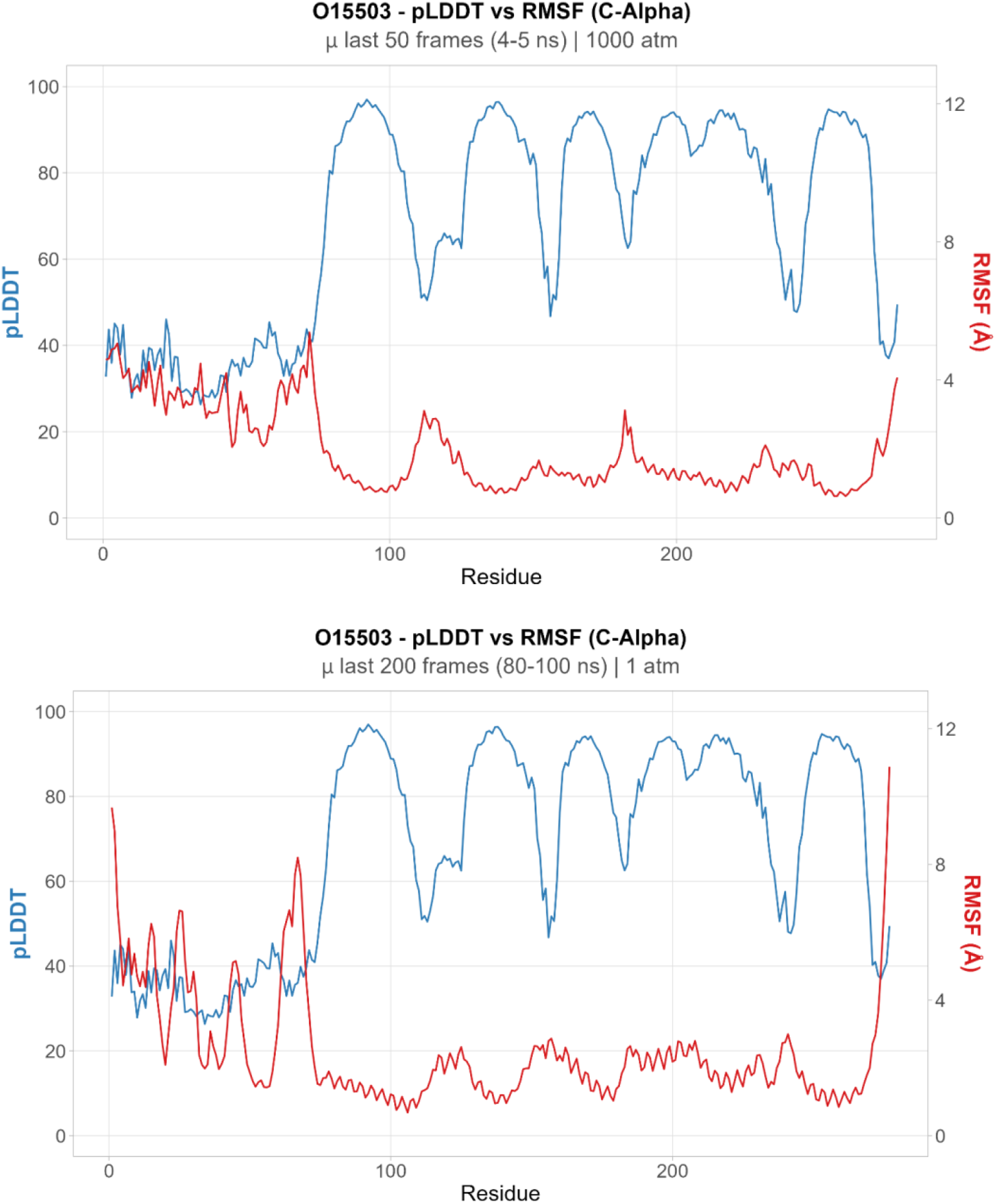
RMSF vs. pLDDT for protein O15503 (average pLDDT = 68.25). Top panel: 5 ns high-pressure simulation. Bottom panel: 200 ns standard pressure simulation. Regions with low pLDDT (blue) correspond to high RMSF (red), indicating structural fragility.

**Figure 5.**
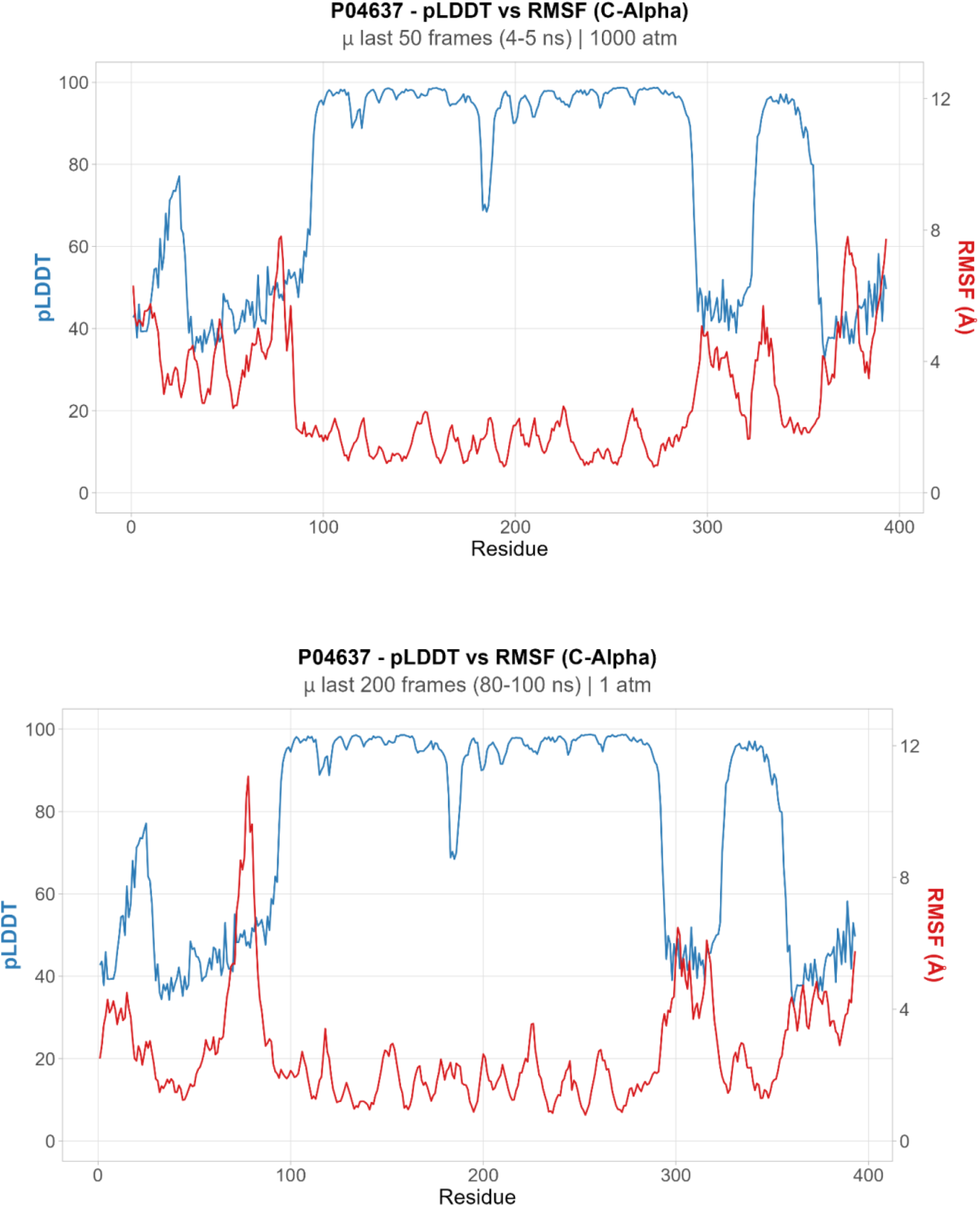
RMSF vs. pLDDT for protein P04637 (average pLDDT = 75.06). Top panel: 5 ns high-pressure simulation. Bottom panel: 200 ns standard pressure simulation. The correspondence between low pLDDT and high RMSF is maintained across both simulation protocols.

**Figure 6.**
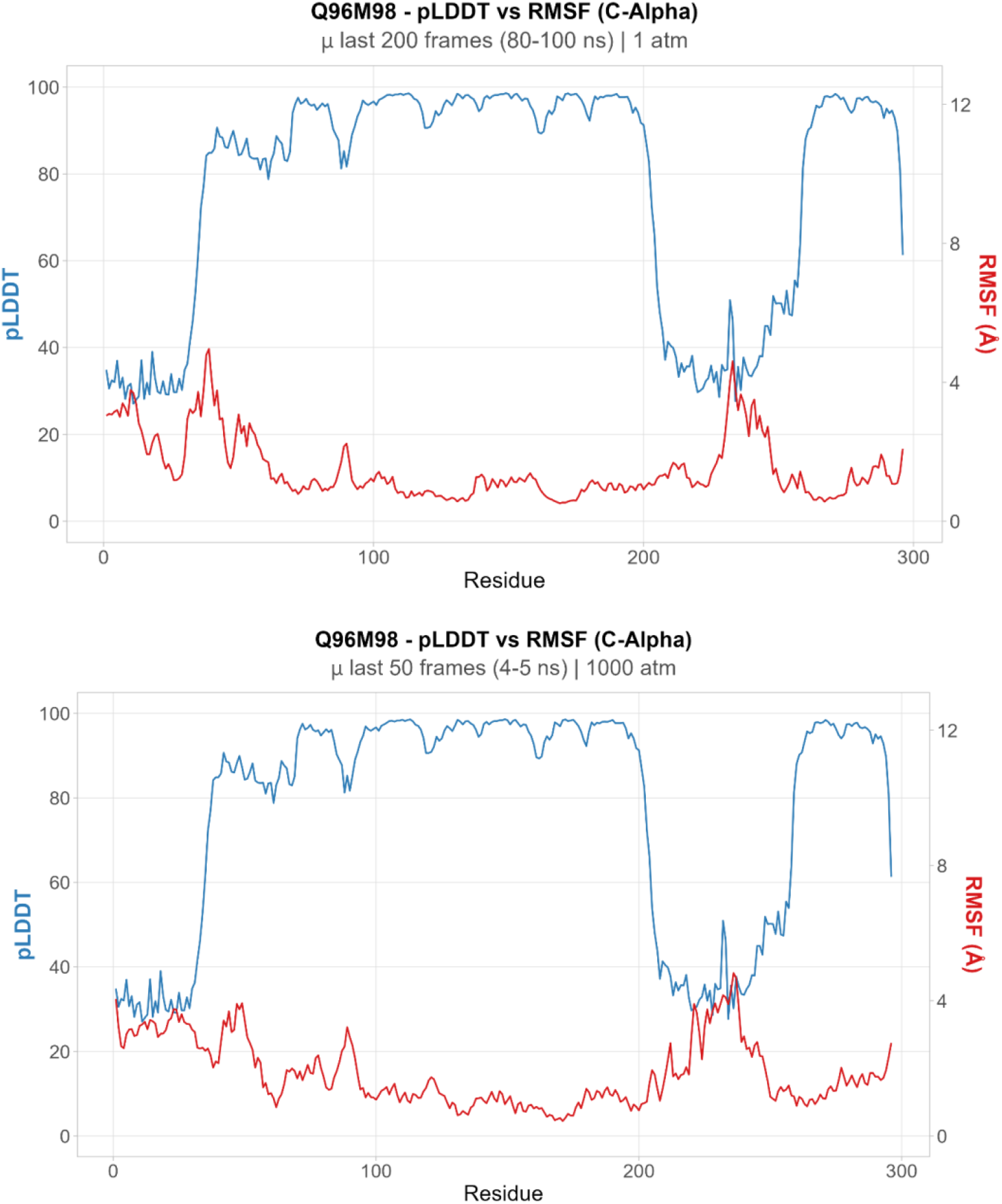
RMSF vs. pLDDT for protein Q96M98 (average pLDDT = 76.44). Top panel: 5 ns high-pressure simulation. Bottom panel: 200 ns standard pressure simulation. A distinct domain with low pLDDT exhibits the highest RMSF values, demonstrating that high-quality predictions reflect not only accurate geometry but also dynamic stability.

These results demonstrate that AlphaUnfold not only correlates with global structural stability (RMSD) but also accurately identifies local regions of structural fragility (RMSF). The consistency between short-time high-pressure and long-term standard pressure simulations validates the efficiency of the AlphaUnfold protocol for rapid model validation.

## 5. Discussion

### 5.1 AlphaUnfold as a Rapid Validation Framework

The results presented in this study demonstrate that AlphaUnfold provides a viable, computationally efficient framework for assessing the biophysical robustness of AlphaFold3-generated protein models. The strong inverse correlation between average pLDDT and RMSD (R^2^ = 0.78, p < 0.001) indicates that AlphaFold3’s confidence metric is a reliable predictor of dynamic structural stability under molecular dynamics simulation. This finding extends previous observations that pLDDT correlates with experimental B-factors and validates the use of pLDDT as a proxy for structural quality [15], [16].

Critically, the consistency between short-time high-pressure (5 ns at 1000 atm) and long-term standard pressure (200 ns at 1 atm) simulations demonstrates that baric stress can accelerate the identification of structural vulnerabilities without fundamentally altering the spatial distribution of instability. The standard error of RMSD during the final 20% of high-pressure simulations was significantly lower than during standard simulations, indicating rapid convergence to a metastable state. This convergence behavior makes AlphaUnfold particularly attractive for high-throughput applications where computational resources are limited.

The local correspondence between low pLDDT and high RMSF further validates the AlphaUnfold approach. Regions predicted with low confidence consistently exhibited high dynamic fluctuations, pinpointing specific metastable areas that may be prone to unfolding or require experimental validation. This local resolution is particularly valuable for protein engineering applications, where identifying and stabilizing vulnerable regions can improve overall protein robustness [21], [22].

### 5.2 Limitations and Outliers

While the overall correlation between pLDDT and dynamic stability is strong, notable outliers were observed. Protein Q96M98-2, with an average pLDDT of 85%, exhibited an RMSD of 11.7 Å, substantially higher than the 6.1 Å predicted by the regression model. This discrepancy highlights an important limitation: pLDDT does not account for post-translational modifications or non-covalent interactions that may be critical for structural stability.

Disulfide bonds between cysteine residues are a common example. AlphaFold3 predicts the spatial proximity of cysteines but does not explicitly model disulfide bond formation. In molecular dynamics simulations, disulfide bonds must be explicitly defined during system preparation. Failure to include these stabilizing interactions can lead to artificially elevated RMSD values even in high-confidence models. This limitation underscores the importance of careful system preparation and highlights the complementary nature of AlphaUnfold: it serves not as a replacement for experimental validation but as a rapid screening tool to prioritize models for further investigation.

Other potential sources of discrepancy include the absence of cofactors, metal ions, or ligands that may stabilize protein structure in vivo but are not included in the AlphaFold3 prediction or MD simulation. Additionally, some proteins may exhibit intrinsic disorder or conformational flexibility that is functionally relevant but appears as structural instability in MD simulations [23].

### 5.3 Implications for Protein Engineering and SCP Development

The integration of AlphaFold3 prediction, BLOSUM-guided rational design, and AlphaUnfold validation provides a powerful workflow for engineering nutritionally enhanced proteins for single-cell protein production. This workflow addresses a critical challenge in SCP development: how to systematically improve amino acid composition without compromising protein stability, solubility, or expression yield.

The rapid turnaround time of AlphaUnfold (5 ns simulations vs. 200+ ns conventional simulations) makes it feasible to screen large libraries of designed variants, identifying promising candidates for experimental validation. This computational pre-screening can substantially reduce the experimental burden and accelerate the development cycle for nutritionally enhanced SCP.

Furthermore, the ability to identify local regions of structural fragility (via RMSF analysis) provides actionable information for iterative design. Regions with low pLDDT and high RMSF can be targeted for stabilization through additional mutations, insertion of stabilizing motifs, or fusion with stable scaffold domains. This iterative design-test-refine cycle, enabled by the rapid feedback provided by AlphaUnfold, represents a significant advance over traditional trial-and-error approaches.

### 5.5 Future Directions

Several avenues for future development of the AlphaUnfold framework are apparent. First, the pipeline could be extended to include additional structural metrics such as hydrogen bond analysis, secondary structure evolution, and contact map analysis. These metrics would provide deeper insights into the molecular mechanisms of structural instability and could guide more targeted stabilization strategies.

Second, the integration of machine learning models trained on the correlation between pLDDT and dynamic stability could enable predictive scoring of AlphaFold3 models without the need for explicit MD simulation. Such models could serve as ultra-rapid pre-screening tools, reserving full AlphaUnfold analysis for the most promising candidates.

Third, the AlphaUnfold framework could be adapted for other AI-based structure prediction tools beyond AlphaFold3, including RoseTTAFold, ESMFold, and other emerging methods. Comparative analysis across different prediction platforms could reveal systematic biases and inform best practices for model selection.

Finally, the development of a web-based interface for AlphaUnfold would greatly enhance accessibility for the broader research community. Such an interface could allow users to submit sequences or AlphaFold3 models, configure simulation parameters, and receive automated analysis reports, democratizing access to this powerful validation tool.

## 6. Conclusion

This study presents AlphaUnfold, an automated pipeline that couples AlphaFold3 structure prediction with short-time high-pressure molecular dynamics simulations to provide rapid, physics-based validation of AI-generated protein models. Testing on a diverse set of proteins revealed a strong inverse correlation between AlphaFold3’s pLDDT confidence metric and structural drift during MD simulation (R^2^ = 0.78, p < 0.001), demonstrating that prediction confidence is a reliable indicator of dynamic stability. Critically, short-time high-pressure simulations (5 ns at 1000 atm) captured the same structural vulnerabilities as long-term standard pressure simulations (200 ns at 1 atm), validating the efficiency of the AlphaUnfold protocol.

Local analysis revealed that regions with low pLDDT consistently exhibited high RMSF, pinpointing specific metastable areas prone to unfolding. This local resolution provides actionable information for protein engineering, enabling targeted stabilization of vulnerable regions.

AlphaUnfold provides a viable, computationally efficient framework for assessing the biophysical robustness of AI-generated models, offering an “experimental-like” validation that ensures more reliable downstream applications in structural biology, protein engineering, and biotechnology. The pipeline is freely available at https://github.com/pegados/pipeline_AlphaUnfold, enabling broad adoption by the research community.

## Acknowledgements

The authors thank the members of the Bioinformatics and Computational Biology groups at UFMG and UFRN for valuable discussions and computational support. We acknowledge the use of computational resources provided by the High-Performance Computing facilities at UFMG.

## References

[1] Foley, J. A., Ramankutty, N., Brauman, K. A., Cassidy, E. S., Gerber, J. S., Johnston, M., et al. (2011). Solutions for a cultivated planet. Nature, 478(7369), 337–342. 10.1038/nature10452

[2] Tilman, D., Balzer, C., Hill, J., & Befort, B. L. (2011). Global food demand and the sustainable intensification of agriculture. Proceedings of the National Academy of Sciences, 108(50), 20260–20264. 10.1073/pnas.1116437108

[3] Lipinsky, E. S., & Litchfield, J. H. (1970). Single-cell protein. Annual Review of Microbiology, 24(1), 585–602. 10.1146/annurev.mi.24.100170.003101

[4] Anupama, R., & Ravindra, P. (2000). Value-added food: Single cell protein. Biotechnology Advances, 18(6), 459–470. 10.1016/S0734-9750(00)00045-8

[5] Becker, E. W. (2007). Micro-algae as a source of protein. Biotechnology Advances, 25(2), 207–210. 10.1016/j.biotechadv.2006.11.002

[6] Ritala, A., Häggman, H., Toivonen, L., Laakso, S., & Mäkinen, O. (2017). Single cell protein—state-of-the-art, industrial landscape and patents 2001–2016. Frontiers in Microbiology, 8, 2009. 10.3389/fmicb.2017.02009

[7] Matassa, S., Boon, N., Pikaar, I., & Verstraete, W. (2016). Microbial protein: future sustainable food supply route with low environmental footprint. Microbial Biotechnology, 9(5), 568–575. 10.1111/1751-7915.12369

[8] Rasi, S., Piras, F., Sandei, L., & Tabasso, S. (2021). Single-Cell Protein from Agro-Industrial Waste: A Review. Fermentation, 7(4), 284. 10.3390/fermentation7040284

[9] Chen, H., Cai, W., Hu, X., Wu, C., & Yan, C. (2005). Single-cell protein diet of a novel recombinant vitellogenin yeast enhances growth and survival of first-feeding tilapia (Oreochromis mossambicus) larvae. The Journal of Nutrition, 135(3), 513–518. 10.1093/jn/135.3.513

[10] van der Poel, A. F. B., van Diepen, J. C. J. M., & Kwakkel, R. P. (2014). The protein challenge: The role of new protein sources for animal feed and human consumption. Animal Feed Science and Technology, 197, 1–13. 10.1016/j.anifeedsci.2014.07.003

[11] FAO. (2017). The future of food and agriculture: Trends and challenges. Food and Agriculture Organization of the United Nations. Rome.

[12] Sali, A., Overington, J. P., Johnson, M. S., & Blundell, T. L. (1994). From protein structure to protein function. Trends in Biochemical Sciences, 19(5), 205–210. 10.1016/0968-0004(94)90023-X

[13] Dill, K. A., MacCallum, J. L., & Song, J. (2008). Protein folding and misfolding. Annual Review of Biochemistry, 77, 79–106. 10.1146/annurev.biochem.77.061306.123622

[14] Henikoff, S., & Henikoff, J. G. (1992). Amino acid substitution matrices from protein blocks. Proceedings of the National Academy of Sciences, 89(22), 10915–10919. 10.1073/pnas.89.22.10915

[15] Jumper, J., Evans, R., Pritzel, A., Green, T., Figurnov, M., Ronneberger, O., et al. (2021). Highly accurate protein structure prediction with AlphaFold. Nature, 596(7873), 583–589. 10.1038/s41586-021-03819-2

[16] Abramson, J., Adler, J., Dunger, J., Evans, R., Green, T., Pritzel, A., et al. (2024). Accurate structure prediction of biomolecular interactions with AlphaFold 3. Nature, 630(8016), 493–500. 10.1038/s41586-024-07487-w

[17] Karplus, M., & McCammon, J. A. (2002). Molecular dynamics simulations of biomolecules. Nature Structural Biology, 9(9), 646–652. 10.1038/nsb0902-646

[18] Sarupria, S., & Garde, S. (2010). Studying pressure denaturation of a protein by molecular dynamics simulations. Proteins: Structure, Function, and Bioinformatics, 78(7), 1641–1651. 10.1002/prot.22680

[19] Paschek, D., Hempel, S., & García, A. E. (2008). Computing the stability diagram of the Trp-cage miniprotein. Proceedings of the National Academy of Sciences, 105(46), 17754–17759. 10.1073/pnas.0804775105

[20] Ng, P. C., & Henikoff, S. (2003). SIFT: Predicting amino acid changes that affect protein function. Nucleic Acids Research, 31(13), 3812–3814. 10.1093/nar/gkg509

[21] Meller, A., Bhakat, S., Solieva, S., & Bowman, G. R. (2023). Accelerating cryptic pocket discovery using AlphaFold. Journal of Chemical Theory and Computation, 19(14), 4355–4363. 10.1021/acs.jctc.2c01189

[22] Raisinghani, N., Alshahrani, M., Gupta, G., Verkhivker, G., & Tao, P. (2024). Exploring conformational landscapes and binding mechanisms of convergent evolution for the SARS-CoV-2 spike Omicron variant complexes with the ACE2 receptor using AlphaFold2-based structural ensembles and molecular dynamics simulations. Physical Chemistry Chemical Physics, 26, 15418–15434. 10.1039/D4CP01372G

[23] van der Lee, R., Buljan, M., Lang, B., Weatheritt, R. J., Daughdrill, G. W., Dunker, A. K., et al. (2014). Classification of intrinsically disordered regions and proteins. Chemical Reviews, 114(13), 6589–6631. 10.1021/cr400525m

[24] Heo, L., & Feig, M. (2022). Multi-state modeling of G-protein coupled receptors at experimental accuracy. Proteins: Structure, Function, and Bioinformatics, 90(11), 1873–1885. 10.1002/prot.26382

[25] Sala, D., Hildebrand, P. W., & Meiler, J. (2023). Biasing AlphaFold2 to predict GPCRs and kinases with user-defined functional or structural properties. Frontiers in Molecular Biosciences, 10, 1121962. 10.3389/fmolb.2023.1121962

[26] Meller, A., Bhakat, S., Solieva, S., & Bowman, G. R. (2023). Discovery of cryptic pockets in the AI-predicted structure of PPM1D phosphatase explains the binding site and potency of its allosteric inhibitors. Frontiers in Molecular Biosciences, 10, 1171143. 10.3389/fmolb.2023.1171143

[27] Stank, A., Kokh, D. B., Fuller, J. C., & Wade, R. C. (2016). Protein binding pocket dynamics. Accounts of Chemical Research, 49(5), 809–815. 10.1021/acs.accounts.5b00516

