## Supplemental Data 1 for "AlphaUnfold: Probing Potential Unfolding and Structural Fragility in AlphaFold3 Models via Short-Time High-Pressure MD"

UNIVERSIDADE FEDERAL DO RIO GRANDE DO NORTE - UFRN
PROGRAMA DE PÓS-GRADUAÇÃO EM BIOINFORMÁTICA - PPgBioINFO

### ALPHAUNFOLD: PIPELINE DE ALTO DESEMPENHO APLICADO À BIOINFORMÁTICA ESTRUTURAL

NATAL/2026

### ALPHAUNFOLD: PIPELINE DE ALTO DESEMPENHO APLICADO À BIOINFORMÁTICA ESTRUTURAL

Dissertação de Mestrado apresentada ao Programa de Pós-Graduação em Bioinformática da Universidade Federal do Rio Grande do Norte como requisito para a obtenção do grau de Mestre em Bioinformática.

Linha de pesquisa:
Desenvolvimento de Produtos e Processos

NATAL/2026

#### AGRADECIMENTOS

**EPÍGRAFE**

"Porque se uma máquina, um Exterminador, pode aprender o valor da vida humana, talvez um dia a gente também possa."

(Sarah Connor)

**RESUMO**

Inicialmente, este trabalho concentrou-se no desenvolvimento do sistema A-DOBRA, um webservice de alto desempenho voltado à submissão e execução automatizada de predições estruturais com AlphaFold em ambiente de computação de alto desempenho (HPC). Com a evolução do projeto, o A-DOBRA passou a integrar o AlphaUnFold, um pipeline mais abrangente que combina inteligência artificial, automação computacional e dinâmica molecular para ampliar o acesso a ferramentas avançadas de modelagem e análise proteica. A pesquisa foi iniciada com o AlphaFold2 e posteriormente expandida para incorporar o AlphaFold3, aumentando a capacidade de modelagem estrutural e de interação entre biomoléculas. Além disso, foi integrado um módulo de dinâmica molecular, permitindo avaliar não apenas as estruturas preditas, mas também sua estabilidade e comportamento conformacional ao longo do tempo. Estudos de caso foram conduzidos em parceria com um projeto de doutorado, aplicando a plataforma em problemas reais e explorando seu potencial para geração de descobertas científicas. Os resultados demonstram que os artefatos desenvolvidos atuam não apenas como ferramentas computacionais, mas como um ambiente integrado para pesquisa em bioinformática estrutural.

**Palavras-chave:** Computação de Alto Desempenho (HPC); Bioinformática Estrutural; AlphaFold; Inteligência Artificial.

#### ABSTRACT

Initially, this work focused on the development of A-DOBRA, a high-performance web service designed for the automated submission and execution of structural predictions using AlphaFold in a high-performance computing (HPC) environment. As the project evolved, A-DOBRA became part of AlphaUnFold, a broader pipeline that combines artificial intelligence, computational automation, and molecular dynamics to expand access to advanced tools for protein modeling and analysis. The research began with AlphaFold2 and was later expanded to incorporate AlphaFold3, increasing its structural modeling capabilities and support for biomolecular interactions. In addition, a molecular dynamics module was integrated, enabling the evaluation not only of predicted structures, but also of their stability and conformational behavior over time. Case studies were conducted in collaboration with a doctoral research project, applying the platform to real-world problems and exploring its potential for scientific discoveries. The results demonstrate that the developed artifacts function not only as computational tools, but also as an integrated environment for research in structural bioinformatics.

**Keywords:** High-Performance Computing (HPC; Structural Bioinformatics; AlphaFold; Artificial Intelligence

**SUMÁRIO**

**LISTA DE FIGURAS**

#### LISTA DE TABELAS

### ALPHAUNFOLD: PIPELINE DE ALTO DESEMPENHO APLICADO À BIOINFORMÁTICA ESTRUTURAL

#### 1. INTRODUÇÃO

Não existiria vida se não fossem as proteínas. É uma proposição que reflete a importância dessas moléculas para os mais diversos processos biológicos existentes nas diferentes formas de seres vivos. Proteínas são macromoléculas que apresentam estrutura tridimensional, um vasto leque de funções, formatos e ligações químicas, que são determinantes para compreender e representar suas estruturas. Devido a toda essa complexidade, prever com acurácia em um sistema vivo tem sido uma tarefa desafiadora. (BRÄNDÉN; TOOZE, 1999).

A compreensão detalhada da estrutura tridimensional das proteínas tornou-se fundamental para o avanço de diversas áreas científicas e tecnológicas, especialmente no desenvolvimento farmacêutico, na biotecnologia e na medicina de precisão. O conhecimento preciso de como uma proteína se dobra e interage com outras moléculas, pode contribuir com a criação de enzimas industriais mais eficientes e o desenvolvimento de terapias personalizadas para doenças complexas. Tradicionalmente, métodos experimentais como cristalografia de raios-X, ressonância magnética nuclear (RMN) e criomicroscopia eletrônica têm sido empregados para determinar essas estruturas, porém tais técnicas demandam recursos consideráveis, tempo extenso e nem sempre são aplicáveis a todas as proteínas. Esta limitação impulsionou o desenvolvimento de abordagens computacionais baseadas em inteligência artificial, que emergiram como alternativas promissoras para acelerar e democratizar o acesso à informação estrutural de proteínas, revolucionando assim o panorama da biologia estrutural e suas aplicações práticas (JUMPER *et al.*, 2021).

##### 1.1 BIOLOGIA ESTRUTURAL NA BIOINFORMÁTICA

A Biologia Estrutural na bioinformática constitui um campo interdisciplinar que integra métodos computacionais, estatísticos e biológicos para analisar e predizer a organização tridimensional de macromoléculas. Nesse contexto, técnicas de bioinformática desempenham papel central ao permitir a manipulação de grandes volumes de dados biológicos, como sequências de aminoácidos, alinhamentos múltiplos (MSA) e bancos de estruturas experimentais. Esses dados são explorados por algoritmos capazes de identificar padrões evolutivos, relações de homologia e sinais de coevolução entre resíduos, fornecendo subsídios fundamentais para inferir estruturas e funções moleculares. Dessa forma, a bioinformática não apenas complementa, mas também potencializa as abordagens experimentais tradicionais, ampliando significativamente a capacidade de investigação estrutural em larga escala (LEHNINGER; NELSON; COX, 2018).

Com o avanço recente da inteligência artificial, essas abordagens foram ainda mais fortalecidas por modelos capazes de integrar informações evolutivas, físicas e geométricas em sistemas preditivos altamente precisos. Ferramentas como o AlphaFold exemplificam essa evolução ao utilizar redes neurais profundas para inferir estruturas tridimensionais diretamente a partir de sequências, incorporando dados derivados de alinhamentos múltiplos e bases estruturais conhecidas. Essa integração permite não apenas prever a estrutura, mas também estimar a confiabilidade das regiões modeladas, fornecendo uma base robusta para análises posteriores, como estudos de dinâmica molecular e interações intermoleculares. Assim, a bioinformática estrutural consolida-se como um elo entre dados biológicos brutos e conhecimento funcional, impulsionando descobertas em diferentes áreas da biologia e da biotecnologia (JUMPER et al., 2021).

#### 1.2 IMPACTOS DA IA NA BIOLOGIA ESTRUTURAL

Nos últimos anos, a biologia estrutural passou por uma transformação significativa com a incorporação de métodos computacionais baseados em inteligência artificial. Ferramentas de predição estrutural, como AlphaFold, demonstraram que é possível prever estruturas proteicas com alta precisão a partir da sequência de aminoácidos, ampliando consideravelmente o acesso à informação estrutural e complementando abordagens experimentais tradicionais (JUMPER *et al.*, 2021).

Paralelamente, avanços em técnicas experimentais têm ampliado a capacidade de determinar e analisar estruturas biomoleculares com maior resolução e em condições mais próximas do ambiente fisiológico. O desenvolvimento da criomicroscopia eletrônica de alta resolução tem permitido investigar complexos biomoleculares grandes e dinâmicos que antes eram difíceis de estudar por cristalografia ou Ressonância Magnética Nuclear (RMN). Essas técnicas possibilitam observar diferentes estados conformacionais de macromoléculas e compreender de forma mais detalhada processos celulares associados à função biológica (CHENG, 2018).

Nesse contexto, a integração entre dados experimentais e modelos computacionais representa uma das principais direções futuras da biologia estrutural. O uso combinado dessas abordagens tem potencial para acelerar a descoberta de novos alvos terapêuticos, aprimorar estratégias de engenharia de proteínas e aprofundar o entendimento de mecanismos moleculares complexos. Entre essas ferramentas emergentes, destacam-se métodos de predição estrutural baseados em aprendizado profundo, que serão discutidos na seção seguinte (VAN DEN BEDEM; FRASER, 2015).

##### 1.3 DO ALPHAFOLD

O alicerce deste trabalho é o sistema de predição estrutural AlphaFold, desenvolvido pela DeepMind, empresa de pesquisa em inteligência artificial pertencente à Google. Em sua segunda versão, o AlphaFold2 (e posteriormente na versão 3) introduziu uma abordagem baseada em aprendizado profundo capaz de prever a estrutura tridimensional de proteínas a partir de suas sequências de aminoácidos com um nível de precisão comparável, em muitos casos, ao obtido por métodos experimentais tradicionais.

O funcionamento do algoritmo baseia-se na integração de informações evolutivas e estruturais por meio de redes neurais profundas, permitindo inferir a estrutura tridimensional de proteínas diretamente a partir de suas sequências de aminoácidos. Esse processo explora padrões conservados observados em alinhamentos múltiplos de sequências e em estruturas previamente determinadas experimentalmente, contribuindo para a alta precisão das predições (JUMPER *et al*., 2021).

> Nossa abordagem alcança níveis de precisão comparáveis a experimentos de resolução experimental de estruturas proteicas, representando um marco na compreensão estrutural de proteínas em escala genômica (Jumper *et al*., 2021).

A competição CASP (Critical Assessment of protein Structure Prediction), realizada bienalmente desde 1994, representa o benchmark mais importante para avaliar métodos de predição de estruturas proteicas, como se fosse as “Olimpíadas de modelagem proteica”. Na CASP14, realizada em 2020, o AlphaFold 2 da DeepMind alcançou um marco histórico ao atingir precisão comparável aos métodos experimentais, com GDT (Global Distance Test) médio superior a 90 para proteínas de dificuldade moderada. Este feito é amplamente reconhecido como o primeiro grande problema científico fundamental resolvido pela inteligência artificial, representando a solução para o desafio de 50 anos do dobramento proteico proposto por Christian Anfinsen. O sistema demonstrou capacidade de prever estruturas com precisão atômica mesmo para proteínas sem homólogos conhecidos, algo anteriormente considerado impossível apenas com métodos computacionais (KRYSHTAFOVYCH *et al*., 2021).

O AlphaFold, portanto, representa uma mudança de paradigma na predição estrutural de proteínas, oferecendo alta precisão na predição de estruturas, capacidade de processamento em larga escala e redução significativa do tempo e custos associados a métodos experimentais tradicionais. O método revoluciona a compreensão estrutural de proteínas, tendo implicações profundas.

#### 1.4 DA DINÂMICA MOLECULAR

A compreensão estrutural apresentada na subseção anterior, especialmente no contexto das predições realizadas por modelos baseados em inteligência artificial, conduz naturalmente à necessidade de discutir a dinâmica conformacional das proteínas. Ainda que uma estrutura tridimensional represente um estado energeticamente favorável, as proteínas não são entidades estáticas, ao contrário, exibem movimentos intrínsecos que variam desde flutuações locais de cadeias laterais até grandes rearranjos dominiais associados à função biológica. Como destacam Nelson e Cox (2021), “as proteínas são moléculas dinâmicas cujas funções dependem de mudanças conformacionais que ocorrem em múltiplas escalas de tempo” (NELSON; COX, 2021). Assim, a análise estrutural deve ser complementada por abordagens que permitam investigar estabilidade, flexibilidade e transições conformacionais, estabelecendo uma ponte entre estrutura prevista e comportamento físico-químico em ambiente simulado.

Nesse contexto, a dinâmica molecular emerge como uma ferramenta central para avaliar a estabilidade estrutural e explorar o espaço conformacional acessível a uma proteína ao longo do tempo. Segundo Daura, van Gunsteren e Mark (2019), simulações de dinâmica molecular permitem “acompanhar a evolução temporal de sistemas biomoleculares, fornecendo informações detalhadas sobre estabilidade estrutural e flutuações atômicas”. Essa abordagem torna-se particularmente relevante na validação de modelos estruturais preditivos, pois métricas como o Root Mean Square Deviation (RMSD) e o Root Mean Square Fluctuation (RMSF), permitem avaliar a estabilidade global e mobilidade local da proteína. Esses parâmetros podem ser analisados em conjunto com o predicted Local Distance Difference Test (pLDDT), contribuindo para correlacionar confiança preditiva e estabilidade dinâmica. Ao decorrer do trabalho, serão apresentadas as bases computacionais dessas simulações e os critérios quantitativos utilizados para integrar modelagem estrutural e validação físico-molecular.

##### 1.5 DA IMPORTÂNCIA DA IA PARA INFERÊNCIA DE PROTEÍNAS

A inteligência artificial tem desempenhado um papel central na transformação da biologia estrutural, especialmente ao enfrentar o tradicional desafio da predição do dobramento de proteínas. Diferentemente dos métodos computacionais clássicos, os modelos baseados em aprendizado profundo conseguem identificar padrões complexos que relacionam a sequência de aminoácidos à estrutura tridimensional das proteínas. Isso ocorre porque essas arquiteturas são capazes de analisar grandes volumes de dados evolutivos e capturar relações entre resíduos distantes na sequência, ampliando significativamente a precisão das predições estruturais (Jumper *et al.*, 2021; Senior *et al.*, 2020).

Os modelos de IA também contribuem para a compreensão das possíveis variações conformacionais das proteínas e de suas interações moleculares. A integração de alinhamentos múltiplos de sequências (MSA) permite explorar sinais evolutivos conservados, tornando a análise estrutural mais eficiente e acessível quando comparada a abordagens tradicionais baseadas apenas em simulação. Esse avanço representa um marco importante para a bioinformática estrutural, ampliando a capacidade de investigação de sistemas biomoleculares complexos (Jumper *et al.*, 2021; Baek *et al.*, 2021).

##### 1.6 DA JUSTIFICATIVA DA PESQUISA

Com base nos argumentos supracitados, a respeito da potencialidade da modelagem e dinâmica de proteínas na biologia estrutural, é de fundamental importância o desenvolvimento de uma ferramenta que facilite no método científico de pesquisadores de diversas áreas das ciências naturais para experimentos com estruturas de proteínas

Para além dos desafios técnicos na área de tecnologia da informação, ferramentas desse tipo supririam a necessidade do alto poder computacional requisitado, que comumente não é evidenciado para pesquisas científicas no país.

Foi lançado em meados de 2024 uma plataforma online pela própria Google, que oferece uma cota individual para envios de trabalhos a serem processados pelo AlphaFold 3, mas nos esbarramos em diversas licenças e limitações, como a proibição de usar a estrutura adquirida em processos de “docking” ou até mesmo de dinâmica molecular, além de não ser usado todo o potencial generativo do AlphaFold em sua mais moderna versão. Visando ainda em ir além dos objetivos iniciais e para implementarmos um potencial mais inovador do trabalho, nos propomos a realizar experimentos e testes com dinâmicas moleculares e análise de métricas que podem ser geradas, não se restringindo apenas à modelagem.

#### 1.7 DA COMPOSIÇÃO DO TRABALHO

Ao longo deste trabalho, o leitor encontrará, inicialmente, uma contextualização sobre a importância da biologia estrutural, suas principais aplicações e o impacto da inteligência artificial na predição de estruturas proteicas, com ênfase no AlphaFold. Em seguida, são apresentados os fundamentos teóricos que sustentam a pesquisa, incluindo conceitos de modelagem de proteínas, computação de alto desempenho, engenharia de software e as ferramentas utilizadas no pipeline. Na sequência, a metodologia descreve de forma detalhada o desenvolvimento do sistema A-DOBRA e do pipeline AlphaUnFold, bem como as decisões técnicas adotadas. Posteriormente, são apresentados e discutidos os resultados, contemplando tanto a implementação do sistema quanto os experimentos de dinâmica molecular e análises estruturais. Por fim, o trabalho se encerra com as considerações finais, nas quais são discutidas as contribuições alcançadas, limitações e possíveis direções futuras da proposta

### 2. OBJETIVOS

#### 2.1 OBJETIVO GERAL

Desenvolver pipeline automatizado para facilitar o processo de modelagem e dinâmica de proteínas, com uso de ferramentas da bioinformática estrutural, utilizando Computação de Alto Desempenho (HPC).

#### 2.2 OBJETIVOS ESPECÍFICOS

- Contribuir com o trabalho de bioinformatas na execução de atividades e experimentos *in silico* que demandam altos custos financeiros e computacionais;
- Disponibilizar plataforma intuitiva de alto nível para profissionais das ciências naturais que não possuem expertise computacional;
- Prover otimizações nos fluxos de execução de modelagem e dinâmica de proteínas *in silico*;
- Divulgar à comunidade científica os artefatos gerados pela pesquisa.

### 3. REFERENCIAL TEÓRICO

#### 3.1 MODELAGEM DE PROTEÍNAS

A modelagem de proteína é um processo computacional custoso, do ponto de vista técnico e científico, pois exige tecnologias avançadas e passíveis de erro para gerar bases de dados, como a cristalografia de raios-x. Obter a estrutura terciária da proteína é o objetivo desse tipo de modelagem e partimos do pressuposto de que apenas com a sequência primária, ou sequência de aminoácidos, podemos adquiri-la (ANFINSEN, 1973). Embora existam muitos passos acessórios como a busca por homologia ou métodos computacionais avançados, o indispensável realmente é o conjunto de caracteres que representa linearmente os aminoácidos, unidade básica de um peptídeo que, por sua vez, se agrupam formando uma proteína.

> A caracterização experimental de intermediários de dobramento permanece tecnicamente desafiadora porque essas espécies são frequentemente transitoriamente populadas e difíceis de capturar para análise estrutural detalhada, deixando lacunas significativas em nossa compreensão mecanística de como as proteínas alcançam seu dobramento nativo. (ENGLANDER & MAYNE, 2014, p. 1567).

O processo de modelagem busca reproduzir *in silico* o complexo mecanismo de dobramento que ocorre in vivo, partindo da sequência linear de aminoácidos para prever a estrutura tridimensional final, como supracitado. Porém, apesar dos avanços significativos, ainda não compreendemos completamente a cascata de eventos moleculares, ou seja, as interações transitórias e os estados intermediários que a cadeia polipeptídica atravessa durante seu dobramento no ambiente celular. O paradoxo de Levinthal ilustra esta complexidade: se uma proteína explorasse aleatoriamente todas as conformações possíveis, levaria mais tempo que a idade do universo para encontrar sua estrutura nativa, mas na realidade este processo ocorre apenas em milissegundos a segundos. Os métodos atuais de modelagem mitigam esta limitação através de aprendizado de padrões estatísticos de estruturas conhecidas, mas não revelam necessariamente a dinâmica e os mecanismos reais do dobramento proteico, permanecendo como uma "caixa-preta" que produz resultados precisos sem elucidar completamente os princípios físico-químicos subjacentes (DILL & MACCALLUM, 2012).

#### 3.2 CONCEITOS DE DINÂMICA DE PROTEÍNAS

A dinâmica de proteínas trata dos movimentos estruturais que essas macromoléculas apresentam ao longo do tempo, complementando a visão estática obtida por métodos experimentais. Em sistemas biológicos, proteínas exploram diferentes conformações, e essa flexibilidade está diretamente relacionada à sua função, influenciando processos como ligação a ligantes e atividade enzimática (KARPLUS; MCCAMMON, 2002).

Nesse contexto, a dinâmica molecular se destaca como uma abordagem computacional capaz de descrever o comportamento temporal de sistemas biomoleculares em nível atômico. Baseada na mecânica clássica, essa técnica permite observar como interações intermoleculares e condições do ambiente influenciam a evolução estrutural das proteínas, revelando transições conformacionais e estados intermediários (ALLEN; TILDESLEY, 2017).

A integração entre modelagem estrutural e dinâmica molecular tem se consolidado como uma estratégia importante na bioinformática estrutural. Estruturas preditas podem ser analisadas quanto à sua estabilidade e comportamento conformacional, permitindo uma interpretação mais completa do sistema e contribuindo para a validação de modelos computacionais (HOLLINGSWORTH; DROR, 2018).

#### 3.3 HPC - HIGH PERFORMANCE COMPUTING

A computação de alto desempenho surgiu como resposta natural aos desafios computacionais modernos que exigem processamento massivo de dados em tempo viável. No contexto da bioinformática estrutural, especialmente na modelagem de proteínas, a necessidade de processar milhões de interações atômicas e realizar cálculos complexos de energia tornou o HPC não apenas desejável, mas absolutamente essencial para o avanço científico. Essa infraestrutura computacional robusta permite que pesquisadores executem simulações que seriam impossíveis em computadores convencionais, democratizando o acesso a ferramentas antes restritas aos grandes centros de pesquisa. (ABRAHAM; MURRAY, 2020)

##### 3.3.1 O ADVENTO DAS GPU’S

As unidades de processamento gráfico deram uma nova direção para o panorama da computação científica ao demonstrarem capacidades extraordinárias para cálculos paralelos. Conforme destacado por Nickolls e Dally (2010), as GPUs evoluíram de simples aceleradores gráficos para processadores altamente paralelos capazes de executar milhares de operações simultaneamente, oferecendo desempenho superior às CPUs tradicionais em tarefas que envolvem processamento massivo de dados. Essa característica tornou-se particularmente valiosa para simulações moleculares, onde cada átomo pode ser processado independentemente, permitindo acelerações de até cem vezes em comparação com processadores convencionais.

A adoção das GPUs na bioinformática estrutural transformou radicalmente os tempos de execução das análises. Proteínas que antes levavam semanas para serem modeladas agora podem ter suas estruturas preditas em questão de horas, permitindo que pesquisadores avancem mais rapidamente em seus experimentos e hipóteses. Essa aceleração não apenas economiza tempo valioso de pesquisa, mas também reduz significativamente os custos operacionais dos centros de computação, tornando a modelagem de proteínas mais acessível para instituições com orçamentos limitados (ABRAHAM; MURRAY, 2020).

##### 3.3.2 WORKLOAD MANAGER SLURM

O gerenciamento eficiente de recursos computacionais em ambientes de alto desempenho exige sistemas sofisticados de orquestração, e o Slurm emergiu como uma das soluções mais robustas para essa necessidade. Este sistema permite que múltiplos usuários compartilhem recursos computacionais de forma justa e eficiente, garantindo que trabalhos sejam executados na ordem apropriada e que os recursos sejam alocados de maneira otimizada. A capacidade do Slurm de gerenciar filas de trabalho, monitorar o uso de recursos e distribuir tarefas entre diferentes nós computacionais tornou-se fundamental para a operação de clusters modernos (YOO; JOYCE; GHERMAN, 2003).

A implementação do Slurm em ambientes de pesquisa bioinformática oferece vantagens significativas na execução de pipelines complexos como o AlphaFold. Como observado por Yoo e colaboradores (2003), o sistema fornece mecanismos avançados de agendamento que consideram não apenas a disponibilidade de recursos, mas também prioridades de usuários, requisitos específicos de memória e GPU, e até mesmo políticas de uso justo entre diferentes grupos de pesquisa. Essa flexibilidade permite que instituições acadêmicas maximizem o retorno sobre seus investimentos em infraestrutura computacional, garantindo que recursos valiosos sejam utilizados de forma eficiente e equitativa.

##### 3.4 CONCEITOS DE IA E FUNCIONAMENTO DO ALPHAFOLD

Para entendermos o funcionamento do AlphaFold, precisamos explicitar alguns conceitos de Inteligência Artificial. Um dos principais é o de Deep Learning, subcampo do aprendizado de máquina que utiliza redes neurais artificiais para identificar padrões complexos em grandes volumes de dados. É um tipo especial de rede que é utilizada no processo, uma chamada “rede neural de atenção” ou rede *transformer*, que infere tanto por mudanças locais (aminoácidos vizinhos), quanto nas globais (aminoácidos mais distantes na cadeia). Os pesquisadores, do AlphaFold, batizaram essa rede pelo nome de “Evoformer”.

A rede Evoformer é o núcleo do modelo do AlphaFold em sua versão 2. Ela processa simultaneamente duas informações principais: os alinhamentos múltiplos de sequências (MSA), que mostram a relação evolutiva entre proteínas semelhantes, e os pares de resíduos de aminoácidos da sequência-alvo. Durante o treinamento, a Evoformer aprende a identificar padrões de coevolução, isto é, mudanças coordenadas entre posições da sequência que indicam contato físico na estrutura final. Essa rede consegue trabalhar com camadas de atenção bidirecional, permitindo que cada posição “observe” todas as outras e ajuste suas representações internas até que surja uma predição coerente das distâncias e orientações entre os átomos. (JUMPER *et al*., 2021). O resultado é uma representação informativa da proteína, que posteriormente é refinada por módulos geométricos que constroem a estrutura tridimensional.

Durante o desenvolvimento deste trabalho, foi lançado o AlphaFold 3, que diferentemente da versão anterior expandiu as predições para complexos biomoleculares inteiros, capazes de incluir interações entre proteínas, DNA, RNA e pequenas moléculas. Esta nova versão, lançada oficialmente em novembro de 2024, possui em sua uma arquitetura a Evoformer-like, uma release aprimorada da rede original que combina de maneira mais direta informações químicas e estruturais, resultando em predições mais acuradas das interações entre moléculas. Outra inovação é o uso de modelos generativos baseados em difusão, que permitem explorar diferentes conformações e aumentar a precisão dos resultados. Com essas melhorias, o AlphaFold 3 se consolida como uma ferramenta mais completa e versátil para compreender os mecanismos moleculares com nível de detalhe atômico (ABRAMSON *et al*., 2024).

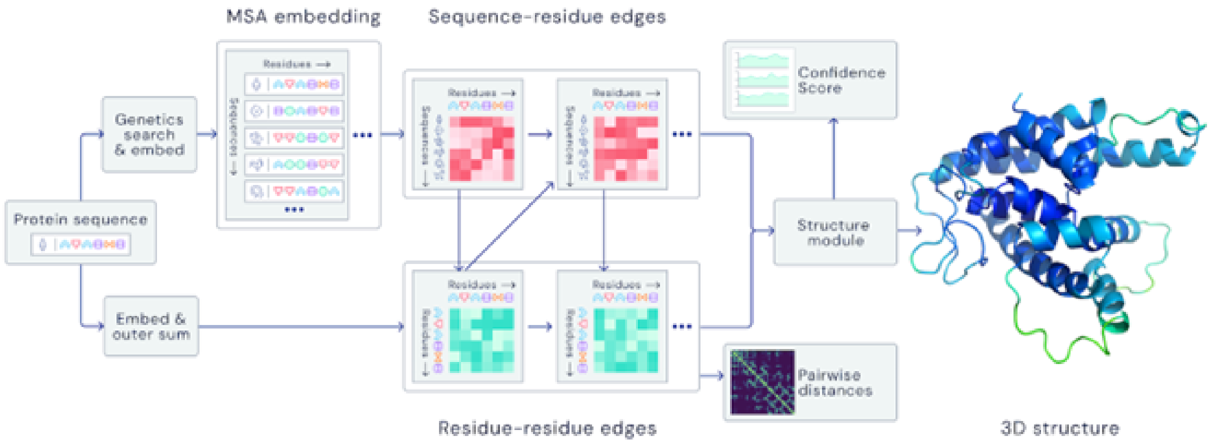

**Figura 01. Esquema simplificado da arquitetura do AlphaFold 2, destacando o fluxo entre embeddings de sequência, Evoformer e módulo estrutural.**
Fonte**:** Adaptado de Jumper *et al*. (2021).

O funcionamento interno do AlphaFold segue um pipeline integrado que transforma a sequência primária de aminoácidos em uma representação tridimensional detalhada da proteína. O processo começa com a busca em bases de dados para gerar os alinhamentos múltiplos de sequências (MSA) e identificar possíveis homólogos estruturais. Esses dados alimentam a rede de atenção — Evoformer na versão 2 e Evoformer-like na versão 3 —, responsável por combinar informações evolutivas e espaciais para compreender as interações entre resíduos. A partir dessas representações, o Structure Module realiza a reconstrução geométrica, ajustando as coordenadas atômicas e refinando o modelo final com base em restrições físico-químicas e índices de confiança, como o pLDDT escore. O resultado é uma estrutura tridimensional altamente precisa, muitas vezes comparável a resultados experimentais (DESAI *et al.*, 2024).

O aprendizado supervisionado e o auto-supervisionado são duas abordagens que apesar das sutilezas individuais, podem caminhar juntas e são bastante utilizadas em sistemas de inteligência artificial. No aprendizado supervisionado, o modelo é treinado a partir de exemplos rotulados, um conjunto de dados cuja relação entre entrada e saída é conhecida, permitindo que ele aprenda a prever resultados ajustando seus parâmetros para minimizar o erro entre a predição e o valor real (Goodfellow, Bengio e Courville, 2016). Já o aprendizado auto-supervisionado não depende de rótulos externos. Ou seja, o próprio modelo formula tarefas internas que o levam a aprender padrões e relações a partir dos dados brutos, explorando estruturas e correlações de forma autônoma (LeCun, Misra e Mnih, 2022).

No caso do AlphaFold, essas duas abordagens são utilizadas de forma integrada. O aprendizado supervisionado é aplicado quando o modelo é exposto a proteínas cujas estruturas tridimensionais já foram determinadas experimentalmente, ajustando suas previsões com base nesses exemplos reais. Paralelamente, o aprendizado auto-supervisionado permite que o sistema aprenda diretamente a partir de grandes conjuntos de sequências sem estrutura conhecida, extraindo informações evolutivas e padrões de coevolução entre resíduos. A combinação dessas técnicas possibilita ao AlphaFold aprender tanto com exemplos diretos quanto com o contexto implícito dos dados biológicos, conferindo uma notável capacidade de generalização e acurácia (JUMPER *et al.*, 2021).

##### 3.4.1 FERRAMENTAS E UTILITÁRIOS DO ALPHAFOLD

O funcionamento do AlphaFold 3 depende de um conjunto integrado de ferramentas da bioinformática responsáveis pela preparação e enriquecimento dos dados de entrada antes do processamento pelo modelo propriamente dito de inteligência artificial. Uma etapa central desse processo é a geração de alinhamentos múltiplos de sequências (MSA), realizada principalmente por ferramentas da família HMMER, como o jackhmmer. Arquiteturalmente, é paralelizado em 4 processos rodando Jackhmmer, a fim de otimizar em tempo essa busca, uma vez que este utilitário executa buscas interativas em grandes bases de dados de sequências proteicas, identificando homólogos evolutivos e construindo perfis estatísticos que capturam relações de conservação e coevolução entre resíduos (EDDY, 2011).

Além do jackhmmer, o AlphaFold emprega outras ferramentas especializadas para ampliar a diversidade e a qualidade dos alinhamentos, bem como para a identificação de estruturas relacionadas. Ferramentas como HHblits e HMMsearch são utilizadas para explorar bases de dados complementares, permitindo a detecção de homologias mais distantes. No caso do AlphaFold 3, esse processo foi estendido para acomodar diferentes tipos de biomoléculas, integrando informações químicas e estruturais adicionais, o que amplia a capacidade do sistema de modelar complexos envolvendo proteínas, ácidos nucleicos e pequenas moléculas (STEINEGGER; SÖDING, 2017).

Ademais, o pipeline do AlphaFold incorpora utilitários voltados ao pós-processamento e à validação dos modelos gerados. Esses componentes incluem rotinas internas para avaliação de confiança estrutural, como o cálculo do pLDDT e métricas de interação intermolecular, além de mecanismos de refinamento geométrico baseados em princípios físico-químicos

#### 3.5 AMBIENTES VIRTUAIS COM MICROMAMBA

Em testes realizados com o AlphaFold 2, pode-se utilizar uma arquitetura alternativa à convencional proposta pela DEEPMIND, que utiliza o Docker. Com isso, é possível e bastante simplificado utilizar ambientes virtuais através da aplicação Micromamba, que como diz na própria documentação oficial, é uma solução autônoma e simplificada para criar e gerenciar ambientes isolados e instalar pacotes em fluxos de trabalho científicos e de desenvolvimento.

> O Micromamba é um binário independente, estaticamente ligado, que pode ser utilizado sem a

> necessidade de uma instalação prévia do Python ou do Conda, tornando-o particularmente adequado para ambientes conteinerizados e de computação de alto desempenho. (Traduzido de MAMBA DEVELOPMENT TEAM, 2023).

Nesse sentido, utilizar Micromamba representa praticidade e isolamento, tendo em vista que a possibilidade de executar a partir de um binário traz muitas vantagens como a portabilidade entre usuários e sistemas. E quanto ao isolamento, traz naturalmente por gerar ambiente virtual independente do sistema operacional hospedeiro.

#### 3.6 RECURSOS DA ENGENHARIA DE SOFTWARE

Em pesquisa e desenvolvimento de software e sistemas são utilizadas diversas ferramentas e princípios da Engenharia de Software, como manutenibilidade, reprodutibilidade, escalabilidade e confiabilidade, que regem o ciclo de vida de todo software.

Para sistemas web uma arquitetura amplamente difundida é a Model–View–Controller (MVC) que como Martin Fowler, referência em arquitetura de software, nos diz

> O Model–View–Controller (MVC) separa a aplicação em três componentes principais: o modelo, que representa o domínio e os dados; a visão, responsável pela apresentação; e o controlador, que coordena a entrada do usuário e a comunicação entre modelo e visão. (FOWLER, 2006).

Esse recorte evidencia que a arquitetura MVC tem como objetivo central a separação de responsabilidades, princípio fundamental da Engenharia de Software. Ao separar a lógica de negócio (Model), a interface de apresentação (View) e o fluxo de controle (Controller), o padrão contribui diretamente para a manutenção e a escalabilidade do sistema. No contexto deste trabalho, a adoção do MVC na interface web permite a evolução independente das camadas do sistema, facilitando assim possíveis ajustes na lógica de aplicação ou na apresentação sem impactos diretos no pipeline científico ou na infraestrutura de computação de alto desempenho integrada (MARTIN, 2017).

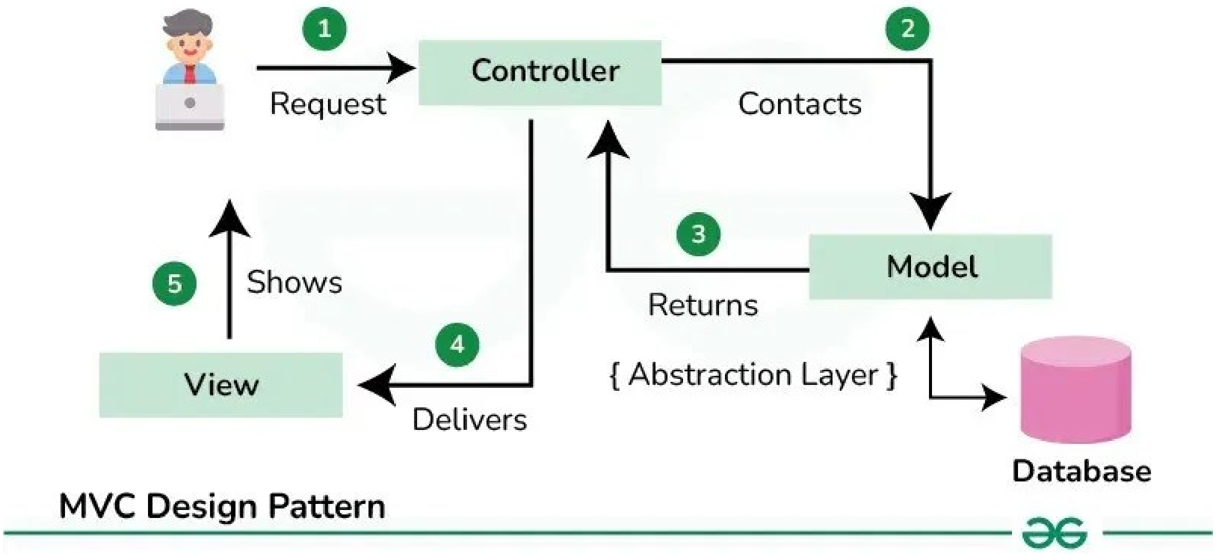

**Figura 02. Funcionamento da arquitetura Model–View–Controller (MVC).**
Fonte: Adaptado de PUTRA et al. (2025).

Como podemos ver no diagrama (Figura 03), no MVC existe a separação de responsabilidades, em que a *View* é responsável pela interface e interação com o usuário, o *Controller* gerencia o fluxo de controle e interpreta as ações recebidas, e o *Model* concentra a lógica de negócio e os dados da aplicação. Essa organização promove desacoplamento entre as camadas, facilitando manutenção, escalabilidade e evolução independente do sistema.

No que tange à reprodutividade, implementações de software que apresentam um pipeline modularizado e reproduzível trazem muitas vantagens. Sendo comumente utilizado em experimentos científicos de entradas, saídas e execuções padronizadas e possibilitando rastreabilidade na busca por erros. Isso torna o produto passível de auditoria, que é um dos pilares da engenharia de software.

Outro ponto importante é a automação de funcionalidades e procedimentos, que é um mecanismo de controle, redução de erros humanos e, por conseguinte, uma característica da confiabilidade. A partir de experimentos sucessivos e observações na lógica de fluxos, é possível agregar tarefas repetitivas e vislumbrar melhorias tanto de recursos computacionais quanto de tempo de execução. (HUMBLE; FARLEY, 2011).

##### 3.6.1 DIVISÃO ENTRE CIÊNCIA E INFRAESTRUTURA

A aplicação de princípios de Engenharia de Software em sistemas científicos é fundamental para garantir flexibilidade, manutenção e evolução ao longo do tempo. Um dos aspectos centrais nesse contexto é a separação entre a lógica científica e a infraestrutura computacional, o que contribui para a redução do acoplamento entre componentes e facilita adaptações futuras. Conforme destacado por Fowler (2006), a separação de responsabilidades permite que diferentes partes do sistema evoluam de forma independente, tornando essa prática especialmente relevante em ambientes de computação de alto desempenho (HPC), onde a forte dependência entre software e hardware pode comprometer a portabilidade e a eficiência.

A adoção desse princípio favorece a execução de aplicações científicas em diferentes arquiteturas computacionais, permitindo maior flexibilidade na utilização de recursos e adaptação a ambientes heterogêneos. Ao desacoplar a lógica do pipeline científico da infraestrutura subjacente, torna-se possível direcionar execuções para diferentes nós computacionais sem alterações na lógica principal do sistema, contribuindo para a consistência e escalabilidade das execuções (EASTMAN; PANDE, 2010).

Além disso, a evolução de sistemas científicos está diretamente associada à capacidade de adaptação e reutilização do software ao longo de seu ciclo de vida. De acordo com Sommerville (2011), sistemas bem projetados devem permitir manutenção e evolução com esforço controlado, considerando desde sua concepção aspectos como modularidade e extensibilidade.

Nesse sentido, a aplicação desses princípios resulta em soluções mais reutilizáveis e flexíveis, capazes de serem executadas em diferentes ambientes com mínimas alterações. Essa característica favorece não apenas a evolução contínua do sistema, mas também a reprodutibilidade dos experimentos e a disseminação das soluções desenvolvidas, aspectos essenciais no contexto da pesquisa científica (DEAN; GHEMAWAT, 2008).

#### 3.7 WEBSERVICES

Webservices, ou sistemas webs, são componentes de software que possibilitam a comunicação entre sistemas diferentes, independentemente da linguagem utilizada e utilizam o protocolo HTTP para que possa haver a transmissão de dados e informações. São utilizados formatos padronizados como JSON ou XML, para federalizar, ou seja, focar em baixa dependência de tipagem de dados, entre as aplicações envolvidas.

A principal característica de um webservice é a exposição de funcionalidades por meio de interfaces bem definidas, permitindo que diferentes sistemas consumam serviços de forma transparente, segura e reutilizável, sem a necessidade de conhecimento prévio sobre a implementação interna do sistema provedor. O que fica nítido segundo a W3C: “Os web services fornecem um meio padronizado de interoperabilidade entre diferentes aplicações de software, executadas em uma variedade de plataformas e frameworks” (W3C, 2004).

A importância dos webservices é evidenciada pelo fato de que eles viabilizam a integração eficiente de sistemas heterogêneos, promovendo flexibilidade em corrigir e reaproveitar componentes de software. Em ambientes científicos e computacionais complexos, como os que envolvem bioinformática estrutural e computação de alto desempenho, os webservices permitem a separação clara entre a camada de apresentação, a lógica de negócio e a infraestrutura computacional subjacente. Essa abordagem facilita a automatização de fluxos de trabalho, o acesso remoto a recursos especializados e a democratização de ferramentas avançadas, tornando possível que usuários sem profundo conhecimento técnico interajam com sistemas de alta complexidade por meio de interfaces simples e padronizadas. Dessa forma, os webservices assumem papel central no desenvolvimento de plataformas escaláveis, distribuídas e orientadas a serviços (NEWMAN, 2021).

#### 3.8 CONTAINERS COM APPTAINER

A utilização de containers tem se consolidado como uma estratégia fundamental para garantir isolamento, portabilidade e reprodutibilidade de ambientes computacionais complexos. Diferentemente de máquinas virtuais tradicionais, os containers compartilham o núcleo do sistema operacional hospedeiro, ao mesmo tempo em que encapsulam dependências, bibliotecas e configurações específicas da aplicação, reduzindo sobrecarga e facilitando a padronização dos ambientes de execução (MERKEL, 2014).

O Apptainer, anteriormente conhecido como Singularity, destaca-se no contexto da computação de alto desempenho por oferecer um modelo de execução compatível com clusters HPC, sem a necessidade de privilégios administrativos durante a execução dos containers. Essa característica é particularmente relevante em ambientes multiusuário, nos quais segurança, controle de permissões e integração com gerenciadores de carga são requisitos essenciais para o uso compartilhado da infraestrutura computacional.

> O Singularity foi projetado para permitir que aplicações científicas sejam executadas de forma segura e reprodutível em ambientes de computação de alto desempenho. Seu modelo elimina a necessidade de privilégios elevados durante a execução, garantindo que o usuário dentro do container possua os mesmos privilégios do usuário no sistema hospedeiro. Essa abordagem facilita a adoção de containers em clusters HPC tradicionais, preservando políticas de segurança e promovendo a portabilidade de aplicações científicas complexas. (KURTZER; SOCHAT; BAUER, 2017).

A utilização de containers, especialmente por meio de tecnologias como o Apptainer, tem sido amplamente adotada em ambientes científicos como estratégia para garantir consistência, portabilidade e reprodutibilidade de aplicações complexas. Essa abordagem permite o empacotamento completo do ambiente de execução, incluindo dependências, bibliotecas e configurações específicas, reduzindo inconsistências entre diferentes nós computacionais e evitando conflitos de software. Além disso, o uso de containers contribui para a padronização dos fluxos de trabalho, facilitando a manutenção, a escalabilidade e a reprodutibilidade de experimentos, aspectos essenciais na bioinformática estrutural e em aplicações baseadas em computação de alto desempenho (KURTZER; SOCHAT; BAUER, 2017).

### 4. METODOLOGIA

#### 4.1 ACESSO ÀS BASES DE DADOS E APRIMORAÇÕES

As bases de dados são disponibilizadas por diferentes instituições que agregam informações sobre proteínas. Os desenvolvedores do Alphafold criaram scripts para realizar os downloads destas bases, que estão disponíveis em diferentes formatos e são utilizados diferentes algoritmos ou métodos de extração e processamento dos dados, a depender do formato.

Segue a lista de bases que devem ser baixadas para a correta execução da aplicação:

```
$DOWNLOAD_DIR/                          # Total: ~ 2.62 TB (download: 556 GB)
    bfd/                                # ~ 1.8 TB (download: 271.6 GB)
        # 6 files.
    mgnify/                             # ~ 120 GB (download: 67 GB)
        mgy_clusters_2022_05.fa
    params/                             # ~ 5.3 GB (download: 5.3 GB)
        # 5 CASP14 models,
        # 5 pTM models,
        # 5 AlphaFold-Multimer models,
        # LICENSE,
        # = 16 files.
    pdb70/                              # ~ 56 GB (download: 19.5 GB)
        # 9 files.
    pdb_mmcif/                          # ~ 238 GB (download: 43 GB)
        mmcif_files/
            # About 199,000 .cif files.
        obsolete.dat
    pdb_seqres/                         # ~ 0.2 GB (download: 0.2 GB)
        pdb_seqres.txt
    small_bfd/                          # ~ 17 GB (download: 9.6 GB)
        bfd-first_non_consensus_sequences.fasta
    uniref30/                           # ~ 206 GB (download: 52.5 GB)
        # 7 files.
    uniprot/                            # ~ 105 GB (download: 53 GB)
        uniprot.fasta
    uniref90/                           # ~ 67 GB (download: 34 GB)
        uniref90.fasta
```

**Figura 03. Nomes e descrições das bases de dados.**
Fonte: https://github.com/google-deepmind/alphafold

Os scripts de download usam um software gerenciador chamado aria2, que suporta múltiplos protocolos (como o HTTP e o FTP), e é capaz de baixar segmentos paralelos de várias fontes ao mesmo tempo. Caso o download seja interrompido, sem sua conclusão, esse gerenciador cria um ponto de recuperação que possibilita retomar o processo, sem recomeçar.

Todavia, encontramos pontos que merecem atenção no que tange os possíveis erros de download e a transferência de dados pré-existentes. Os scripts atuais não possuem caminhos alternativos caso ocorra algum problema com o download aria2 e, além disso, toda vez que são executados transferem a massa de dados das bases sem verificar previamente se houve modificações e se precisam realmente ser feito um novo download.

Ajustamos o script de download para utilizar outra aplicação de download (wget) caso ocorra algum problema na execução do aria2 e, para evitar redundâncias de dowloads, faz comparações para verificar se o arquivo local precisa ser atualizado em relação ao arquivo remoto (hospedado em alguma instituição de pesquisa).

Seguem os fluxogramas comparando a estratégia inicial e a proposta:

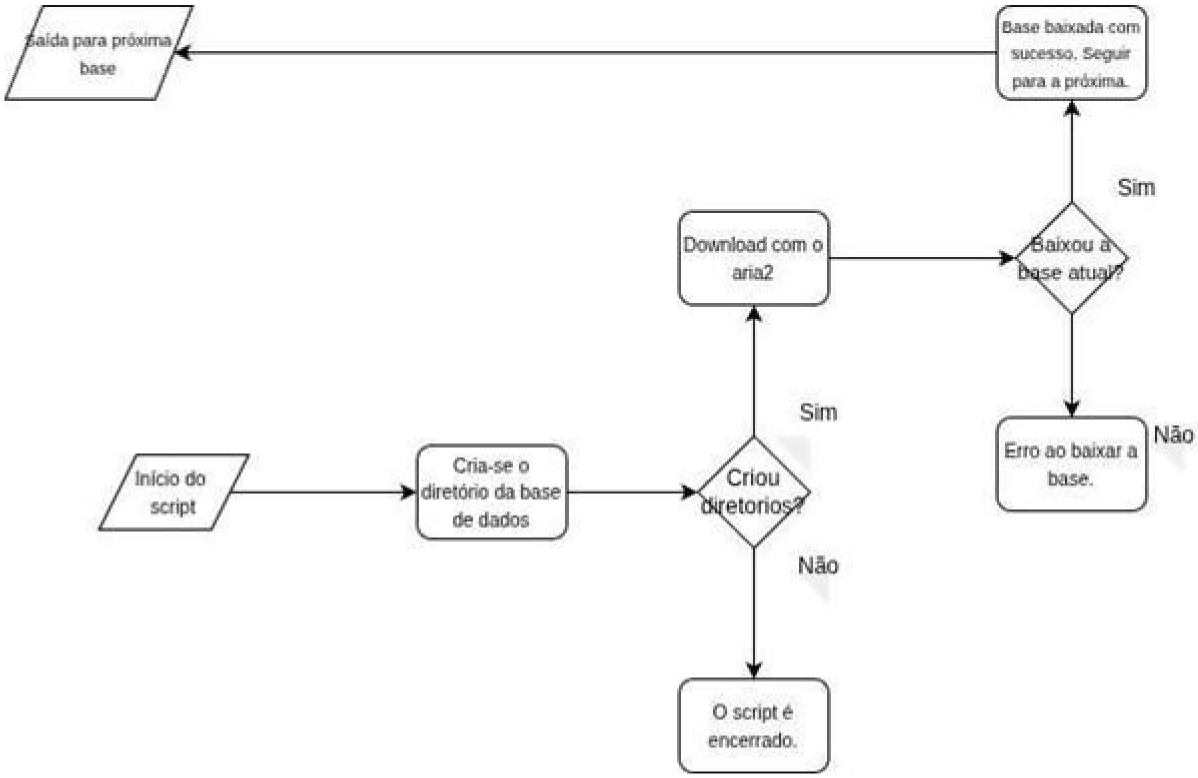

**Figura 04. Fluxograma inicial do download dos databases.** A figura apresenta o processo de obtenção e organização das bases de dados necessárias para execução do AlphaFold, incluindo as etapas de download, armazenamento e preparação dos arquivos utilizados.
Fonte: Os autores, 2024.

Pode-se visualizar (Figura 04) que se houver algum erro na criação do diretório ou no download com o aria2, o script irá ser encerrado sem que seja executado corretamente. Além disso, em todas as execuções serão baixados uma massa de dados sem garantia alguma de duplicação de informações, pois não é realizada verificação alguma dos dados locais com os dados remotos.

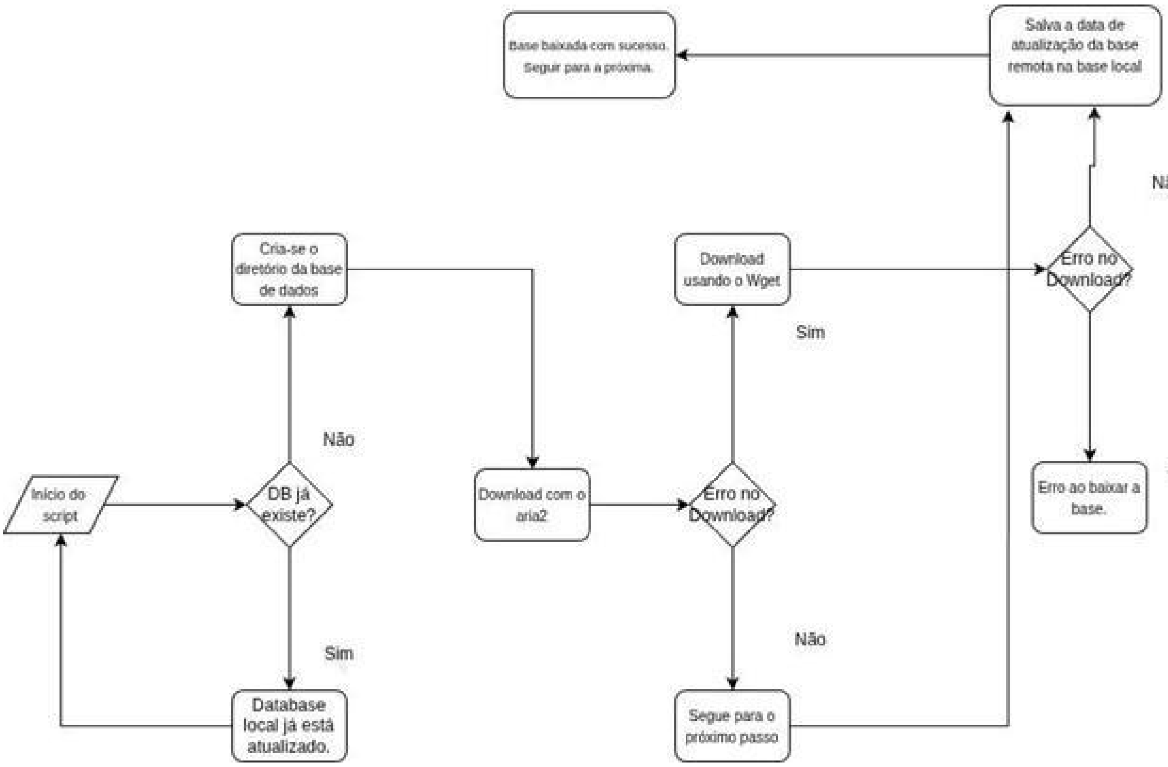

**Figura 05: Fluxograma proposto para o download dos databases.** O diagrama ilustra a estratégia adotada para obtenção, organização e validação das bases de dados necessárias ao funcionamento do AlphaFold 2.
Fonte: Os autores, 2024.

No novo fluxograma (Figura 05), temos três condições determinantes, a inicial, que verifica se o script deve ser iniciado de fato, ou se já existe uma dada base, a que verifica se houve algum problema durante o processo com a aplicação aria2 e a que monitora se ocorreu tudo certo na finalização do script.

Posteriormente, com o lançamento de uma nova versão e arquitetura do AlphaFold, para a versão 3, a forma de realizar o download das bases de dados foi modificado, sendo necessário apenas a execução do script (Algoritmo 01) na localização desejada.

**Algoritmo 01. Script de automação para download e descompactação dos bancos de dados do AlphaFold 3.**

```
#!/bin/bash
      # Copyright 2024 DeepMind Technologies Limited
      #
      # AlphaFold 3 source code is licensed under CC BY-NC-SA 4.0. To
view a copy of
      # this license, visit https://creativecommons.org/licenses/by-
nc-sa/4.0/
      #
```

```
# To request access to the AlphaFold 3 model parameters, follow the process set
# out at https://github.com/google-deepmind/alphafold3. You may only use these
# if received directly from Google. Use is subject to terms of use available at
# https://github.com/google-deepmind/alphafold3/blob/main/WEIGHTS_TERMS_OF_USE.md

set -euo pipefail

readonly db_dir=${1:-$HOME/public_databases}

for cmd in wget tar zstd ; do
  if ! command -v "${cmd}" > /dev/null 2>&1; then
    echo "${cmd} is not installed. Please install it."
  fi
done

echo "Fetching databases to ${db_dir}"
mkdir -p "${db_dir}"

readonly SOURCE=https://storage.googleapis.com/alphafold-databases/v3.0

echo "Start Fetching and Untarring 'pdb_2022_09_28_mmcif_files.tar'"
wget --quiet --output-document=- \
    "${SOURCE}/pdb_2022_09_28_mmcif_files.tar.zst" | \
    tar --no-same-owner --no-same-permissions \
    --use-compress-program=zstd -xf - --directory="${db_dir}" &

for NAME in mgy_clusters_2022_05.fa \
            bfd-first_non_consensus_sequences.fasta \
            uniref90_2022_05.fa uniprot_all_2021_04.fa \
            pdb_seqres_2022_09_28.fasta \
            rnacentral_active_seq_id_90_cov_80_linclust.fasta \
            nt_rna_2023_02_23_clust_seq_id_90_cov_80_rep_seq.fasta \
            rfam_14_9_clust_seq_id_90_cov_80_rep_seq.fasta ; do
  echo "Start Fetching '${NAME}'"
  wget --quiet --output-document=- "${SOURCE}/${NAME}.zst" | \
      zstd --decompress > "${db_dir}/${NAME}" &
done

wait
echo "Complete"
```

Fonte: https://github.com/google-deepmind/alphafold3/blob/main/fetch_databases.sh

Para a utilização do AlphaFold em sua versão 2 fica o artefato para a otimização de ganhos de tempo e menos I/O na aplicação.

#### 4.2 EXECUÇÃO OTIMIZADA DO ALPHAFOLD 2

##### 4.2.1 INSTALAÇÃO DO MICROMAMBA

Primeiro, foi realizado o download e a instalação do Micromamba:

```
bash
curl -Ls https://micro.mamba.pm/api/micromamba/linux-64/latest | tar
-xvj bin/micromamba

sudo ln -s /st02data/bin/micromamba/micromamba
/usr/local/bin/micromamba
```

##### 4.2.2 CONFIGURAÇÃO DO AMBIENTE SHELL

Em seguida, foi configurado o ambiente shell para o Micromamba:

```
bash

/st02data/bin/micromamba/micromamba shell init -s bash -p
/st02data/bin/micromamba
```

Foram criados aliases para facilitar a inicialização:

```
bash
alias micromamba init = /st02data/bin/micromamba/micromamba shell
init -s bash -p /st02data/bin/micromamba && source ~/.bashrc

alias micromamba shell = eval "$(micromamba shell hook --shell
bash)"
```

Inicialização do ambiente:

```
bash

micromamba init
```

##### 4.2.3 CRIAÇÃO DO AMBIENTE PARA ALPHAFOLD

Foi criado um ambiente específico para o AlphaFold com Python 3.8:

```
bash
micromamba create -n alphafold xtensor -c conda-forge python==3.8

micromamba activate alphafold
```

##### 4.2.4 CONFIGURAÇÃO DOS CANAIS E ATUALIZAÇÕES

Foram realizadas configurações adicionais nos canais do Micromamba:

```
bash
micromamba config append channels conda-forge
micromamba config set channel_priority strict
micromamba self-update
micromamba update -n base conda
```

##### 4.2.5 INSTALAÇÃO DAS DEPENDÊNCIAS

Foram instaladas as dependências necessárias para o AlphaFold 2:

```
bash
# Instalação do OpenMM, CUDA e PDBFixer
micromamba install -y -c conda-forge openmm==7.5.1
cudatoolkit==11.2.2 pdbfixer

# Instalação de ferramentas bioinformáticas
micromamba install -y -c bioconda hmmer hhsuite==3.3.0 kalign2
```

##### 4.2.6 INSTALAÇÃO DAS DEPENDÊNCIAS PYTHON

Por fim, foram instaladas as dependências Python específicas para o AlphaFold 2:

```
bash
micromamba pip install absl-py==1.0.0 biopython==1.79 chex==0.0.7
dm-haiku==0.0.9 dm-tree==0.1.6 immutabledict==2.0.0 jax==0.3.25 ml-
collections==0.1.0 numpy==1.21.6 pandas==1.3.4 protobuf==3.20.1
scipy==1.7.0 tensorflow-cpu==2.9.0
```

##### 4.2.7 EXECUÇÃO DO ALPHAFOLD 2

Após a configuração do ambiente e instalação de todas as dependências, o AlphaFold 2 pode ser executado utilizando o script `run_alphafold.sh`:

```
bash
bash run_alphafold.sh -d /st01data/DBalphafold_Mar_24 -o
/home/miguel/output -f /home/miguel/myoglobin.fasta
```

Este comando executa o AlphaFold 2 com os seguintes parâmetros:

- `-d /st01data/DBalphafold_Mar_24`: Especifica o diretório contendo as bases de dados genéticas e estruturais necessárias para o AlphaFold
- `-o /home/miguel/output`: Define o diretório de saída onde serão salvos os resultados da predição
- `-f /home/miguel/myoglobin.fasta`: Indica o arquivo FASTA contendo a sequência de aminoácidos da proteína a ser modelada (neste caso, mioglobina)

O ambiente configurado com Micromamba fornece um sistema isolado e otimizado para a execução do AlphaFold 2. As versões específicas dos pacotes foram selecionadas para garantir compatibilidade com o AlphaFold 2, embora o ambiente possa precisar de ajustes adicionais dependendo das necessidades específicas de modelagem de proteínas.

#### 4.2 ATUALIZAÇÃO PARA O ALPHAFOLD 3

Começamos o nosso trabalho utilizando o AlphaFold em sua versão 2. Como citado, utilizamos uma arquitetura diferenciada para download das bases de dados, uma vez que para termos melhores predições precisávamos usar os bancos de dados estendidos (os desenvolvedores utilizavam bancos de dados menores para disponibilizar à comunidade) que juntos somavam mais de 2.6 Terabytes, tornando inviável downloads redundantes. Além disso, usamos ambientes virtuais com o micromamba, instalando todas as bibliotecas e dependências e formando um ambiente personalizado. Todavia, com o lançamento do AlphaFold 3, em meados de 2024, tivemos que mudar completamente nossa abordagem e pensamos, já se atentando ao futuro, em uma arquitetura que seja mais genérica e escalonável.

O AlphaFold 3 trouxe diversas melhorias, como a diminuição dos volumes de dados que precisavam ser transferidos para o ambiente local, com compressão e melhorias nos tipos de dados. Além de proporcionar um acesso mais rápido a estes bancos de dados. No que tange o provisionamento do ecossistema de execução, os desenvolvedores colocaram o docker como opção oficial, para baixar todas as bibliotecas necessárias e executar o programa. Todavia, utilizamos o Apptainer, por nos proporcionar requisitos que se adequavam mais ao nosso parque computacional. Quanto ao mecanismo de submissão de sequências, a versão 3 trouxe consigo apenas o formato .JSON, um problema que tivemos que contornar, pois o formato popularizado pela comunidade de bioinformática é o .FASTA. Em nosso script final, disponibilizado como artefato no github, detalhamos o processo de conversão, que possibilita ao usuário enviar arquivos tanto no formato .FASTA como no formato .JSON.

##### 4.2.1 EXECUÇÃO DA APLICAÇÃO

Para as primeiras modelagens com o AlphaFold 3, seguimos seguintes passos:

1. Instalação de drivers da NVIDIA;
2. Instalação do Apptainer (para a conversão final);
3. Clone do repositório oficial do AlphaFold 3;
4. Ajustes no Dockerfile para incluir alguma dependência que estava faltando ou atualizar o CUDA;
5. Build: Rodou o docker build;
6. Conversão: Rodou o `apptainer build` para gerar o `.sif`.

Por fim, para execução foi necessário rodar o seguinte comando, com cada argumento detalhado:

```
apptainer exec \
    ${APPTAINER_GPU_FLAG} \
    --bind ${INPUT_DIR}:/data/input \
    --bind ${JOB_OUTPUT_DIR}:/data/output \
    --bind /home/alphafold:/home/alphafold \
    --env OMP_NUM_THREADS=${SLURM_CPUS_PER_TASK} \
    ${CONTAINER} \
    python ${ALPHAFOLD_SCRIPT} \
        --json_path=/data/input/${INPUT_JSON} \
        --output_dir=/data/output \
        --model_dir=${MODEL_DIR} \
        --db_dir=${DB_DIR}
```

`apptainer exec`

Executa um comando dentro de um container Apptainer já existente.

`${APPTAINER_GPU_FLAG}`

Flag condicional para habilitar acesso à GPU (por exemplo, `--nv`), permitindo aceleração por CUDA.

`--bind ${INPUT_DIR}:/data/input`

Monta o diretório de entrada do host no container, tornando os arquivos de entrada acessíveis internamente.

`--bind ${JOB_OUTPUT_DIR}:/data/output`

Mapeia o diretório de saída do container para o host, garantindo persistência dos resultados.

`--bind /home/alphafold:/home/alphafold`

Disponibiliza o diretório do usuário dentro do container, necessário para scripts, permissões ou dependências locais.

`--env OMP_NUM_THREADS=${SLURM_CPUS_PER_TASK}`

Define o número de *threads* OpenMP de acordo com os recursos alocados pelo Slurm.

`${CONTAINER}`

Caminho para a imagem do container Apptainer contendo o AlphaFold configurado.

`python ${ALPHAFOLD_SCRIPT}`

Invoca o script principal de execução do AlphaFold dentro do container.

`--json_path=/data/input/${INPUT_JSON}`

Especifica o arquivo JSON de entrada com a descrição da proteína e parâmetros da execução.

`--output_dir=/data/output`

Diretório onde os resultados da predição estrutural serão gravados.

`--model_dir=${MODEL_DIR}`

Caminho para os pesos e modelos neurais do AlphaFold.

`--db_dir=${DB_DIR}`

Diretório contendo as bases de dados biológicas necessárias para a inferência estrutural.

##### 4.2.2 MÉTRICAS DE CONFIANÇA DO ALPHAFOLD 3

AlphaFold 3 fornece métricas de confiança que ajudam a interpretar a qualidade das estruturas preditas. Essas métricas indicam o nível de confiabilidade dos resultados, tanto em regiões específicas da proteína quanto na estrutura como um todo. Neste trabalho, foram utilizadas as métricas pLDDT, pPDE e ipTM, pois juntas permitem uma análise mais completa da qualidade estrutural das predições.

O pLDDT (predicted Local Distance Difference Test) é uma métrica de confiança local que indica o quão confiável é a posição de cada resíduo na estrutura. Seus valores variam de 0 a 100, sendo que valores mais altos indicam maior confiança. No arquivo .pdb gerado pelo AlphaFold, o pLDDT é armazenado na coluna de *B-factor* (penúltima coluna de cada linha). Embora o valor apareça repetido para todos os átomos de um mesmo resíduo, o mais comum é considerar apenas o carbono alfa (Cα), pois ele representa o esqueleto da proteína e evita redundância na análise. A Figura 06 destaca essa coluna no arquivo, onde estão os valores de pLDDT associados a cada resíduo.

```
MODEL        1
ATOM      1  N   MET A   1       3.693 -45.106  23.930  1.00 60.62           N
ATOM      2  CA  MET A   1       3.200 -44.263  25.047  1.00 60.62           C
ATOM      3  C   MET A   1       3.103 -42.771  24.712  1.00 60.62           C
ATOM      4  CB  MET A   1       4.008 -44.497  26.326  1.00 60.62           C
ATOM      5  O   MET A   1       2.273 -42.105  25.306  1.00 60.62           O
ATOM      6  CG  MET A   1       3.724 -45.901  26.872  1.00 60.62           C
ATOM      7  SD  MET A   1       4.646 -46.319  28.365  1.00 60.62           S
ATOM      8  CE  MET A   1       3.834 -45.250  29.588  1.00 60.62           C
ATOM      9  N   TRP A   2       3.869 -42.239  23.751  1.00 76.69           N
ATOM     10  CA  TRP A   2       3.842 -40.804  23.416  1.00 76.69           C
ATOM     11  C   TRP A   2       2.694 -40.370  22.492  1.00 76.69           C
ATOM     12  CB  TRP A   2       5.205 -40.445  22.819  1.00 76.69           C
ATOM     13  O   TRP A   2       2.204 -39.254  22.619  1.00 76.69           O
ATOM     14  CG  TRP A   2       6.329 -40.618  23.792  1.00 76.69           C
ATOM     15  CD1 TRP A   2       7.116 -41.711  23.941  1.00 76.69           C
ATOM     16  CD2 TRP A   2       6.816 -39.641  24.761  1.00 76.69           C
ATOM     17  CE2 TRP A   2       7.888 -40.229  25.495  1.00 76.69           C
ATOM     18  CE3 TRP A   2       6.474 -38.307  25.073  1.00 76.69           C
ATOM     19  NE1 TRP A   2       8.017 -41.495  24.966  1.00 76.69           N
ATOM     20  CH2 TRP A   2       8.227 -38.207  26.774  1.00 76.69           C
ATOM     21  CZ2 TRP A   2       8.587 -39.535  26.490  1.00 76.69           C
ATOM     22  CZ3 TRP A   2       7.175 -37.597  26.066  1.00 76.69           C
ATOM     23  N   LEU A   3       2.220 -41.249  21.602  1.00 82.38           N
ATOM     24  CA  LEU A   3       1.170 -40.916  20.627  1.00 82.38           C
ATOM     25  C   LEU A   3      -0.132 -40.374  21.262  1.00 82.38           C
ATOM     26  CB  LEU A   3       0.875 -42.138  19.729  1.00 82.38           C
ATOM     27  O   LEU A   3      -0.613 -39.339  20.803  1.00 82.38           O
ATOM     28  CG  LEU A   3       0.573 -41.731  18.275  1.00 82.38           C
ATOM     29  CD1 LEU A   3       1.869 -41.704  17.462  1.00 82.38           C
ATOM     30  CD2 LEU A   3      -0.391 -42.721  17.627  1.00 82.38           C
ATOM     31  N   PRO A   4      -0.683 -40.993  22.331  1.00 86.69           N
ATOM     32  CA  PRO A   4      -1.888 -40.486  22.988  1.00 86.69           C
ATOM     33  C   PRO A   4      -1.670 -39.109  23.621  1.00 86.69           C
ATOM     34  CB  PRO A   4      -2.261 -41.521  24.059  1.00 86.69           C
ATOM     35  O   PRO A   4      -2.550 -38.259  23.561  1.00 86.69           O
ATOM     36  CG  PRO A   4      -1.523 -42.787  23.633  1.00 86.69           C
ATOM     37  CD  PRO A   4      -0.274 -42.238  22.960  1.00 86.69           C
ATOM     38  N   LEU A   5      -0.482 -38.871  24.189  1.00 86.50           N
ATOM     39  CA  LEU A   5      -0.130 -37.597  24.812  1.00 86.50           C
ATOM     40  C   LEU A   5      -0.022 -36.476  23.768  1.00 86.50           C
ATOM     41  CB  LEU A   5       1.194 -37.774  25.580  1.00 86.50           C
ATOM     42  O   LEU A   5      -0.544 -35.384  23.977  1.00 86.50           O
```

**Figura 06: Visualização textual do arquivo .pdb da Proteína Q6NUM9.**
Fonte: Os autores, 2025.

O pPDE (predicted Pairwise Distance Error) estima o erro esperado entre pares de resíduos, permitindo avaliar a consistência estrutural entre diferentes regiões da proteína. Diferente do pLDDT, que é local, o pPDE fornece uma visão mais global da qualidade da estrutura, sendo útil para identificar possíveis inconsistências na organização tridimensional.

O ipTM (interface predicted Template Modeling score) é utilizado para avaliar a qualidade de interações entre cadeias ou domínios, sendo especialmente relevante em complexos proteicos. Seus valores variam entre 0 e 1, onde valores mais próximos de 1 indicam maior confiança nas interfaces previstas. Essa métrica é importante para analisar a confiabilidade de interações intermoleculares e a organização estrutural em sistemas mais complexos.

##### 4.3 TECNOLOGIAS DE DESENVOLVIMENTO WEB UTILIZADAS

Para o desenvolvimento do sistema web, utilizamos o servidor Web Apache, a linguagem de programação PHP com o framework Laravel e o banco de dados que utilizamos foi o PostgreSQL.

Essa pilha de tecnologias adotadas apresenta vantagens significativas em termos de desempenho, segurança e escalabilidade. O servidor web Apache é amplamente reconhecido por sua estabilidade e robustez em ambientes de produção, oferecendo suporte consolidado a aplicações de grande porte e alto número de acessos simultâneos. A linguagem PHP, por sua vez, aliada ao framework Laravel, proporciona uma estrutura moderna para o desenvolvimento de aplicações web, incorporando boas práticas de Engenharia de Software, como organização baseada no padrão Model–View–Controller (MVC), proteção contra vulnerabilidades comuns e facilidade de integração com serviços externos. O uso do banco de dados PostgreSQL complementa esse ecossistema ao oferecer um sistema gerenciador de banco de dados relacional altamente confiável, com suporte avançado a transações, integridade referencial e consultas complexas, características essenciais para o armazenamento consistente e seguro das informações geradas pelo sistema.

A escolha dessas tecnologias foi motivada tanto por critérios técnicos quanto por adequação ao contexto acadêmico e científico no qual o sistema está inserido. Apache, PHP/Laravel e PostgreSQL são bastante utilizados em ambientes institucionais e de pesquisa, contando com extensa documentação, comunidades ativas e suporte contínuo. Além disso, trata-se de um conjunto de ferramentas de código aberto, alinhado aos princípios de reprodutibilidade, transparência e baixo custo, fundamentais em projetos científicos. A familiaridade prévia com essas tecnologias também contribuiu para uma implementação mais eficiente, permitindo maior foco na integração do sistema web com a infraestrutura de computação de alto desempenho (HPC) e no atendimento aos requisitos específicos de bioinformática estrutural proposto neste trabalho.

#### 4.4 REQUISITOS FUNCIONAIS E NÃO-FUNCIONAIS

Os requisitos funcionais do sistema foram definidos com o objetivo de garantir que o usuário consiga executar todo o fluxo de modelagem de proteínas (ou outros experimentos de bioinformática estrutural) de forma autônoma e controlada. O sistema permite o cadastro de usuários, autenticação segura por meio de login, e a submissão de arquivos de sequência no formato JSON ou FASTA, viabilizando a execução do pipeline do AlphaFold 3. Além disso, o usuário pode acompanhar o histórico completo de suas submissões, realizar o download de todos os arquivos gerados durante o processamento e receber notificações claras sobre o sucesso ou falha de cada *job* submetido, assegurando transparência e rastreabilidade dos experimentos realizados.

Os requisitos não-funcionais concentram-se em aspectos de usabilidade, desempenho, organização dos dados e segurança da aplicação. A interface do sistema foi projetada para ser intuitiva, facilitando o uso por pesquisadores sem conhecimento técnico aprofundado em computação. O banco de dados foi estruturado de forma consistente para garantir integridade, eficiência nas consultas e persistência confiável das informações. Adicionalmente, o sistema prioriza a rapidez no download dos artefatos gerados e adota mecanismos de navegação segura, utilizando protocolos de segurança para proteção dos dados e das credenciais dos usuários.

A relação entre os requisitos funcionais e não-funcionais é fundamental para assegurar que as funcionalidades oferecidas sejam executadas de maneira eficiente, segura e acessível. Enquanto os requisitos funcionais definem o que o sistema deve fazer, os requisitos não-funcionais estabelecem como essas operações devem ocorrer, garantindo qualidade, desempenho e confiabilidade. Essa integração assegura que o sistema não apenas atenda às necessidades científicas dos usuários, mas também ofereça uma experiência estável, segura e adequada ao contexto de pesquisa em bioinformática estrutural.

#### 4.5 DESENVOLVIMENTO E FUNCIONAMENTO DO SISTEMA A-DOBRA

O pipeline é composto de duas grandes partes: Um HPC e um servidor Web. O HPC é gerenciado pelo Workloader Slurm, que é uma espécie de orquestrador de máquinas virtuais ou nós físicos em um cluster. Com ele é possível ter uma visão sobre todos os nós, balanceamento de carga e gerenciar os recursos computacionais.

No cluster HPC foi montada uma estrutura de containers, com o uso do Apptainer, que oferece uma segregação do ambiente principal do sistema operacional, que é o gatilho de execução do AlphaFold.

O servidor web, por sua vez, roda em PHP com o framework laravel, provendo de um SGBD PostgreSQL. O servidor web faz todo o gerenciamento de usuários e trabalhos enviados, com persistência de dados.

O servidor web, com o envio do arquivo a ser modelado por parte do usuário, faz uma requisição através de tunelamento SSH e acessa o cluster disparando um script que aciona o Apptainer configurado com o AlphaFold e é enviado um trabalho ao Slurm, que direciona, com diretrizes de eficiência, para o melhor nó no momento. O servidor web fica verificando a fila de trabalhos até o alvo atual ser concluído. Após a conclusão, os outputs são resgatados pelo servidor web e serão disponibilizados para o usuário. Com a evolução e novas exigências da pesquisa, o sistema A-dobra passou a integrar o pipeline AlphaUnFold, como o módulo de modelagem de proteínas.

#### 4.6 PREPARAÇÃO DO PIPELINE ALPHAUNFOLD

A preparação do pipeline AlphaUnFold foi pensada para automatizar um fluxo mais abrangente de modelagem e análise estrutural de proteínas. Implementado por meio de scripts em Shell, Python e TCL (Linguagem nativa do software VMD - Visual Molecular Dynamics), o pipeline executa de maneira sequencial as etapas de geração dos modelos, preparação dos sistemas e submissão das simulações, podendo ser executado posteriormente scripts de geração de métricas e gráficos. Essa organização permite maior flexibilidade em ambientes de HPC, além de facilitar a adaptação do fluxo e a inclusão de novas análises conforme a necessidade.

As simulações de dinâmica molecular foram realizadas com o NAMD3, utilizando o VMD no preparo dos sistemas e na extração das métricas. Para reduzir o custo computacional, foram adotadas simulações curtas, de aproximadamente 5 ns, sob condição de alta pressão (1000 atm). Essa estratégia favorece uma exploração conformacional mais rápida, permitindo observar mudanças estruturais relevantes em menor tempo.

Como em meados de novembro de 2023 a Deepmind lançou o Alphafold 3 para a comunidade científica e consolidou seu servidor web para envio de sequências e entregas de modelagem, analisamos a possibilidade de incrementar a dinâmica molecular em nosso trabalho, a fim de expandir nossas features e podermos entregar mais artefatos oriundos da pesquisa e resultados para a comunidade científica. A este pipeline demos o nome de AlphaUnFold.

Em parceria com outra pesquisa, no âmbito de um trabalho de doutoramento, foram conduzidos experimentos envolvendo proteínas em sua forma selvagem, bem como variantes mutantes obtidas por substituições específicas de aminoácidos por resíduos considerados essenciais. Essas modificações tiveram como objetivo avaliar a viabilidade estrutural das proteínas modeladas, por meio da análise de métricas de globularidade e de qualidade estrutural, permitindo comparar de forma sistemática os efeitos das mutações introduzidas.

Adicionalmente, no âmbito dessa pesquisa parceira, foi demonstrado que a adoção de protocolos de simulação ajustados, envolvendo a modulação controlada de parâmetros termodinâmicos, como temperatura e pressão, possibilita a obtenção de dinâmicas moleculares de curta duração (da ordem de 5 nanossegundos) com qualidade comparável àquelas tradicionalmente obtidas em simulações significativamente mais longas (até 200 nanossegundos). Esse resultado é particularmente relevante, pois estabelece uma estratégia eficiente para redução substancial do custo computacional e do tempo de execução, viabilizando a análise de múltiplos sistemas proteicos em cenários de computação de alto desempenho. Vale salientar que simulações de 200 ns podem demorar vários dias, dependendo do tamanho da proteína, enquanto as nossas demoraram apenas algumas horas, entre 3 horas e 8 horas, variando conforme a proteína. O trabalho de doutorado da colaboradora SILVA, Juliana (2022-atual) discorre sobre essa descoberta.

##### 4.6.1 PROTEÍNAS SELECIONADAS PARA ANÁLISE

###### 4.6.1.1 ANÁLISE DO PLDDT

Para estudos de correlações de métricas do Alphafold com métricas de estabilidade estrutural, foram selecionadas proteínas humanas com diferentes perfis de pLDDT, abrangendo desde estruturas com baixa confiança preditiva até modelos altamente confiáveis. As proteínas, identificadas pelo ID Uniprot foram organizadas de modo a representar esse espectro, permitindo analisar como a qualidade estrutural prevista se correlaciona com métricas dinâmicas como RMSD e RMSF.

A proteína Q14236, proveniente de Homo sapiens, apresenta baixo pLDDT médio, indicando que grande parte de sua estrutura é prevista com baixa confiança, o que sugere elevada flexibilidade estrutural ou possível presença de regiões intrinsicamente desordenadas. De forma semelhante, O15503 exibe pLDDT intermediário, refletindo a coexistência de regiões parcialmente estruturadas com segmentos menos definidos conformacionalmente. Já a proteína Q9ULZ0 apresenta pLDDT elevado, indicando uma organização estrutural mais consistente, com domínios bem definidos segundo a predição computacional.

Entre as proteínas com maior confiabilidade estrutural, destaca-se P04637, amplamente estudada experimentalmente e conhecida por seu papel central na regulação do ciclo celular, apresentando pLDDT muito elevado. As proteínas Q96M98 e sua isoforma Q96M98-2 também exibem pLDDT alto, sugerindo estruturas compactas e bem organizadas, embora pequenas variações de sequência possam introduzir diferenças estruturais sutis entre as isoformas. De forma semelhante, P0DP23 apresenta alto grau de conservação estrutural e pLDDT muito elevado, reforçando a robustez de sua predição.

Por fim, a isoforma Q6NUM9-2 mantém pLDDT elevado, indicando que alterações de isoforma não comprometem significativamente a confiança estrutural global. A inclusão dessas proteínas permite uma análise comparativa robusta, abrangendo diferentes níveis de confiança preditiva e possibilitando investigar, de maneira integrada, como essas variações influenciam a estabilidade e a dinâmica estrutural observadas em simulações computacionais. Para ilustrar a relação entre confiança preditiva e comportamento dinâmico em nível local, foi investigada a correlação entre pLDDT e RMSF por resíduo, permitindo avaliar como a mobilidade estrutural varia ao longo da sequência da proteína.

###### 4.6.1.2 MIOGLOBINA 5XLO

Em um dos estudos de casos foi selecionada a mioglobina, representada pela estrutura 5XLO do Protein Data Bank (PDB). Trata-se de uma proteína globular clássica, responsável pelo armazenamento de oxigênio em tecidos musculares, com estrutura predominantemente composta por alpha hélices organizadas de forma compacta. Sua estabilidade estrutural e ampla caracterização experimental a tornam um modelo adequado para avaliar a robustez de predições obtidas com AlphaFold e sua validação por dinâmica molecular.

A mioglobina apresenta tamanho moderado (≈150 resíduos) e organização estrutural bem definida, permitindo analisar de forma controlada o impacto de perturbações introduzidas artificialmente. Neste contexto, foram realizadas substituições progressivas de resíduos por arginina (R) ao longo da sequência.

###### 4.6.1.2 FASEOLINA 2PHL

A faseolina, representada pela estrutura 2PHL do Protein Data Bank (PDB), é a principal proteína de reserva do feijão comum (*Phaseolus vulgaris*), pertencente à família das globulinas 7S. É uma proteína bastante estudada, com estrutura tridimensional bem definida e predominantemente formada por folhas beta organizadas de forma compacta. Sua função está relacionada ao armazenamento de aminoácidos durante o desenvolvimento da semente, sendo utilizada no processo de germinação. Por apresentar alta estabilidade estrutural e boa caracterização experimental, a faseolina se mostra um modelo adequado para estudos de engenharia de proteínas, permitindo avaliar com maior segurança os efeitos de modificações na sequência.

Neste trabalho, as substituições de aminoácidos foram realizadas com base em matrizes do tipo BLOSUM, que indicam quais trocas são mais prováveis ao longo da evolução. Esse tipo de abordagem permite escolher substituições mais realistas do ponto de vista biológico, levando em conta propriedades como tamanho, carga e características químicas dos resíduos. Foram consideradas tanto substituições mais conservadoras, que tendem a manter a estrutura original, quanto substituições menos conservadoras, que podem causar alterações mais significativas. Com isso, foi possível gerar diferentes variantes da faseolina e analisar como essas mudanças influenciam sua estabilidade e comportamento dinâmico ao longo das simulações.

#### 4.7 MÉTRICAS DE AVALIAÇÃO E ESTABILIDADE ESTRUTURAL

A avaliação da estabilidade estrutural de proteínas em simulações de dinâmica molecular requer o uso de métricas que capturem diferentes aspectos do comportamento conformacional ao longo do tempo. Neste trabalho, foram utilizadas quatro métricas complementares, RMSD, RMSF, SASA e raio de giro, por permitirem uma análise integrada entre estabilidades global e local, exposição ao solvente e grau de compactação estrutural. A escolha dessas métricas se justifica por serem amplamente consolidadas na literatura e por fornecerem uma visão abrangente da integridade estrutural das proteínas modeladas.

##### 4.7.1 RMSD

O RMSD (Root Mean Square Deviation) foi empregado como principal indicador de estabilidade global, medindo o desvio médio da estrutura em relação a uma conformação de referência. Para o cálculo, foram considerados os últimos 50 frames da simulação, correspondentes ao último nanossegundo, de modo a focar na região já estabilizada da trajetória. Cada frame foi alinhado à estrutura inicial e o desvio médio foi então calculado, permitindo avaliar a conservação estrutural ao final da dinâmica.

$$RMSD = \sqrt{\frac{1}{n} \sum_{i=1}^{n} (x_{i1} - x_{i2})^2 + (y_{i1} - y_{i2})^2 + (z_{i1} - z_{i2})^2}$$

**Figura 07. Fórmula do RMSD**
Fonte: OnlineBioinfo, 2022.

Essa expressão (Figura 07) descreve como o RMSD é calculado a partir das diferenças de posição dos átomos entre duas estruturas. Para cada átomo, são comparadas as coordenadas *x, y* e *z* em dois momentos, geralmente entre a estrutura inicial e um frame da simulação. Essas diferenças são elevadas ao quadrado, somadas e depois divididas pelo número total de átomos considerados, obtendo-se uma média. Ao final, aplica-se a raiz quadrada para trazer o valor de volta à unidade de distância, resultando em uma medida global de quanto a estrutura se desviou ao longo do tempo.

##### 4.7.2 RMSF

O RMSF (Root Mean Square Fluctuation) foi utilizado para avaliar a mobilidade local de cada resíduo, sendo calculado a partir dos átomos de carbono alfa (Cα), que representam o esqueleto da proteína. Essa métrica mede a variação da posição de cada resíduo em relação à sua posição média ao longo da simulação, permitindo identificar regiões mais flexíveis ou instáveis.

$$RMSF_{\text{átomo}} = \sqrt{\frac{1}{t}\sum_{i=1}^{t}(x_i^{ref}-x_i)^2+(y_i^{ref}-y_i)^2+(z_i^{ref}-z_i)^2}$$

**Figura 08. Fórmula do RMSF**
Fonte: OnlineBioinfo, 2022.

Essa expressão (Figura 08) mostra como o RMSF é calculado para cada átomo ao longo da simulação. Em vez de comparar duas estruturas específicas, aqui a posição de cada átomo é comparada com sua posição média ao longo do tempo. Para isso, são consideradas as variações nas coordenadas *x, y* e *z* em cada frame em relação a essa posição de referência, elevando essas diferenças ao quadrado e fazendo a média sobre os frames analisados. Ao final, a raiz quadrada fornece uma medida da flutuação daquele átomo, indicando o quanto ele oscila durante a simulação.

##### 4.7.3 RAIO DE GIRO

O raio de giro (Rg) foi calculado a partir desses mesmos frames, com base na distribuição dos átomos em relação ao centro de massa, indicando o grau de compactação estrutural na fase final da simulação.

$$R_g = \sqrt{\frac{\sum_{i=1}^{N} m_i \cdot \|r_i - r_{cm}\|^2}{\sum_{i=1}^{N} m_i}}$$

**Figura 09. Fórmula do Raio de Giro**
Fonte: LEACH, 2001.

O raio de giro (Rg) mede o grau de compactação da proteína, indicando como os átomos estão distribuídos em relação ao centro de massa. Como mostrado na Figura 09, ele é calculado a partir das distâncias dos átomos ao centro de massa, considerando suas massas. Valores menores indicam estruturas mais compactas, enquanto valores maiores sugerem estruturas mais abertas.

##### 4.7.4 SASA

A SASA (Solvent Accessible Surface Area) foi obtida considerando também os últimos 50 frames, por meio do cálculo da área acessível ao solvente com o uso de uma sonda que percorre a superfície da proteína.

$$SASA = \sum_{i=1}^{N} A_i$$

**Figura 10. Representação conceitual da SASA**
Fonte: HOLLINGSWORTH, 2018.

A SASA representa a área da proteína que está acessível ao solvente. Diferentemente das métricas anteriores, a expressão apresentada (Figura 10) é conceitual, indicando que a área total resulta da soma das contribuições de cada átomo, e não de uma fórmula analítica direta. Na prática, o cálculo é feito por métodos numéricos, simulando uma esfera que percorre a superfície da proteína para identificar as regiões expostas ao solvente.

#### 4.8 FERRAMENTA OTIMIZADA PARA REPRODUZIR RESULTADOS

A extração e análise dessas métricas foram realizadas utilizando o software VMD (Visual Molecular Dynamics), por meio de scripts em linguagem TCL integrados ao pipeline desenvolvido. As trajetórias foram processadas frame a frame, aplicando seleções específicas de átomos para cada métrica, como o uso dos carbonos alfa no cálculo do RMSF. A partir desses dados, foram geradas séries temporais e valores médios, permitindo uma análise detalhada do comportamento dinâmico e da estabilidade estrutural das proteínas ao longo das simulações.

Foram utilizados dois scripts para automatizar o fluxo de análise. O script em TCL (Algoritmo 02) foi utilizado dentro do VMD para processar as trajetórias e calcular as métricas, realizando a leitura dos arquivos, aplicação das seleções atômicas e extração das métricas, sendo utilizados para geração de métricas os 20% finais das trajetórias geradas pelas simulações. Em seguida, o script em Python (Algoritmo 03) foi responsável pelo tratamento dos dados gerados, organizando os resultados, calculando valores médios e preparando os arquivos para análise e visualização.

**Algoritmo 02. Script TLC para geração de métricas**

```
set psf_file [lindex $argv 0]
set dcd_file [lindex $argv 1]

set dir_traj [file dirname [file normalize $dcd_file]]
cd $dir_traj

set arq_stats "raw_stats.csv"
set arq_rmsf  "raw_rmsf.csv"

mol new $psf_file type psf
mol addfile $dcd_file type dcd waitfor all

set nframes [molinfo top get numframes]
set min_frames 250
set probe 1.4

if {$nframes < $min_frames} {
    set f [open $arq_stats w]
    puts $f "Arquivo;Metrica;Valor"
    puts $f "SISTEMA;STATUS;INCOMPLETO"
    close $f
    exit
}

set ini [expr int($nframes * 0.8)]

set sel_ca  [atomselect top "protein and name CA"]
set ref_ca  [atomselect top "protein and name CA" frame 0]
set sel_all [atomselect top "protein and noh"]

set lista_rmsd {}
set lista_rog  {}
```

```
set lista_sasa {}

for {set i $ini} {$i < $nframes} {incr i} {
    $sel_ca frame $i
    $sel_all frame $i

    $sel_all move [measure fit $sel_ca $ref_ca]

    lappend lista_rmsd [measure rmsd $sel_ca $ref_ca]
    lappend lista_rog  [measure rgyr $sel_all]
    lappend lista_sasa [measure sasa $probe $sel_all]
}

proc calc_stats {valores} {
    set soma 0
    foreach v $valores {
        set soma [expr $soma + $v]
    }

    set media [expr $soma / double([llength $valores])]

    set soma_quad 0
    foreach v $valores {
        set soma_quad [expr $soma_quad + pow($v - $media, 2)]
    }

    set desvio [expr sqrt($soma_quad / double([llength $valores]))]

    return [list $media $desvio]
}

set f [open $arq_stats w]
puts $f "Arquivo;Metrica;Valor"

foreach chave {rmsd rog sasa} nome {RMSD RoG SASA_Total} {
    set resultado [calc_stats [set lista_$chave]]
```

```
    puts $f "SISTEMA;$nome;[lindex $resultado 0]"
    puts $f "SISTEMA;${nome}_SD;[lindex $resultado 1]"
}

close $f

set valores_rmsf [measure rmsf $sel_ca first $ini last [expr $nframes
- 1] step 1]
set residuos     [$sel_ca get resid]

set f2 [open $arq_rmsf w]
puts $f2 "Residuo;RMSF"

foreach r $residuos v $valores_rmsf {
    set v2 [string map {. ,} $v]
    puts $f2 "$r;$v2"
}

close $f2

exit
```

Fonte: Os Autores, 2025.

**Algoritmo 03. Script Python para Processamento das métricas**

```
import subprocess
import pandas as pd
from pathlib import Path

base = Path.cwd()
vmd = r"C:\Program Files (x86)\University of Illinois\VMD\vmd.exe"
tcl = "analise_unificada.tcl"

rmsf_dir = base / "RMSF_MEDIO"
rmsf_dir.mkdir(exist_ok=True)
```

```
for pasta in [p for p in base.iterdir() if p.is_dir() and p.name !=
"RMSF_MEDIO"]:
    nome = pasta.name

    psf = next(pasta.glob("*final.psf"), None)
    dcds = list(pasta.rglob("*.dcd"))

    if psf and dcds:
        for dcd in dcds:
            print(f"Analisando ID: {nome}")
            subprocess.run([vmd, "-dispdev", "text", "-e", tcl, "-
args", str(psf), str(dcd)])

    stats = []
    for f in pasta.rglob("raw_stats.csv"):
        df = pd.read_csv(f, sep=";")
        for _, r in df.iterrows():
            try:
                stats.append((r["Metrica"],
float(str(r["Valor"]).replace(",", "."))))
            except:
                pass

    if stats:
        df = pd.DataFrame(stats, columns=["Metrica", "Valor"])
        resumo =
df.groupby("Metrica")["Valor"].mean().round(4).reset_index()
        resumo["Modelo"] = nome
        resumo["Valor_Final"] =
resumo["Valor"].astype(str).str.replace(".", ",")
        resumo[["Modelo", "Metrica", "Valor_Final"]].to_csv(pasta /
"RESUMO.csv", sep=";", index=False)

    for f in pasta.rglob("raw_rmsf.csv"):
        dest = rmsf_dir / f"{nome}.csv"
        if dest.exists():
```

```
            dest.unlink()
        f.rename(dest)

    for f in pasta.rglob("raw_stats.csv"):
        try:
            f.unlink()
        except:
            pass

resumos = list(base.rglob("RESUMO.csv"))

if resumos:
    df = pd.concat([pd.read_csv(f, sep=";") for f in resumos],
ignore_index=True)
    metricas = ["RMSD", "RMSD_SD", "RoG", "RoG_SD", "SASA_Total",
"SASA_Total_SD"]

    tabela = []
    for m in metricas:
        linha = {"Metrica": m}
        for mod in df["Modelo"].unique():
            v = df[(df.Modelo == mod) & (df.Metrica ==
m)]["Valor_Final"]
            linha[mod.upper()] = v.iloc[0] if not v.empty else "N/A"
        tabela.append(linha)

    pd.DataFrame(tabela).to_csv("RESULTADOS_RMSD_RoG_SASA.csv",
sep=";", index=False)

print("Processo concluído! RMSFs em: RMSF_MEDIO")
```

Fonte: Os Autores, 2025.

### 5. RESULTADOS E DISCUSSÃO

#### 5.1 SISTEMA A-DOBRA

O sistema foi construído respeitando o paradigma de orientação a objetos, com atributos e funções bem definidas (Figura 11). Partindo do pressuposto de que um software tem um ciclo de vida e que seu fim não termina com a primeira versão, aprimorações e novas versões serão consideradas para garantir a manutenibilidade e escalabilidade do projeto, otimizando o desempenho. No entanto, estudos de caso apontam que o sistema atua em atendimento às expectativas e está em produção.

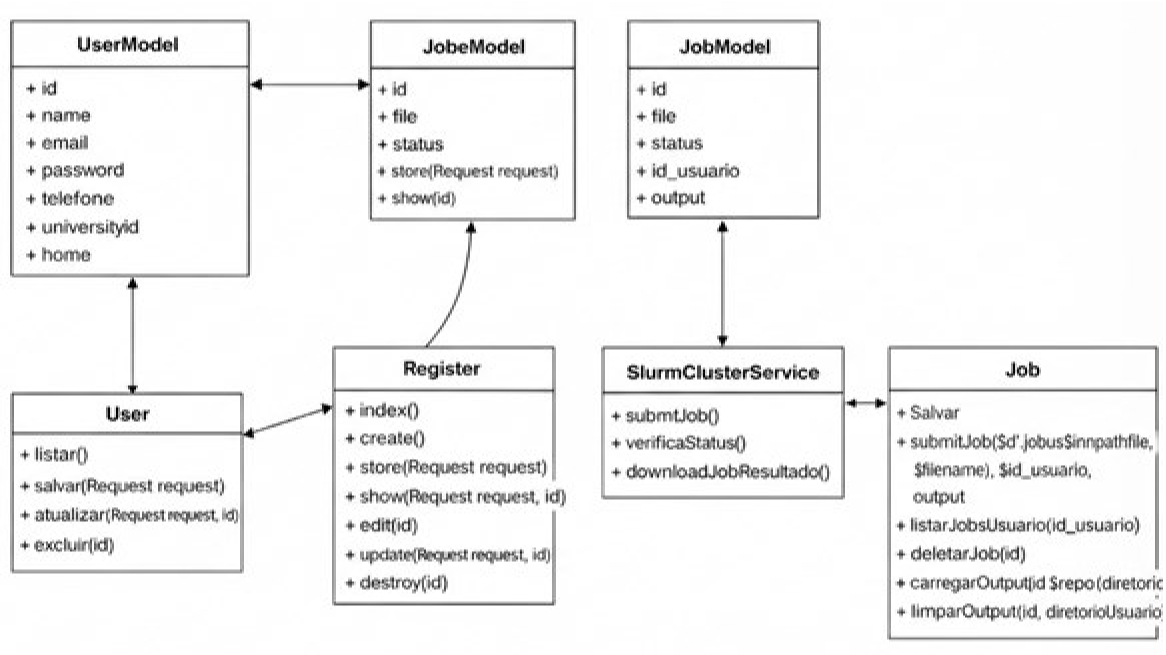

**Figura 11: Diagrama de classes do sistema A-DOBRA.** O esquema apresenta as principais classes da aplicação e suas relações, com destaque para o módulo SlurmClusterService, responsável pela integração com o ambiente veredas.lcc.ufmg.br e pela etapa mais sensível do sistema: o envio, gerenciamento e acompanhamento das execuções no cluster via Slurm**.** Fonte: Os autores, 2025.

No modelo de Entidade-Relacionamento do projeto (Figura 11), a entidade *Users* representa os usuários cadastrados no sistema, os quais submetem *jobs*. Cada *job* é submetido por um único usuário (1,1), enquanto um usuário pode submeter zero ou muitos *jobs* (0,n). Após a submissão, cada *job* possui exatamente uma execução associada à entidade *fila_jobs*, que representa a fila de processamento no Slurm e registra o estado de execução do trabalho.

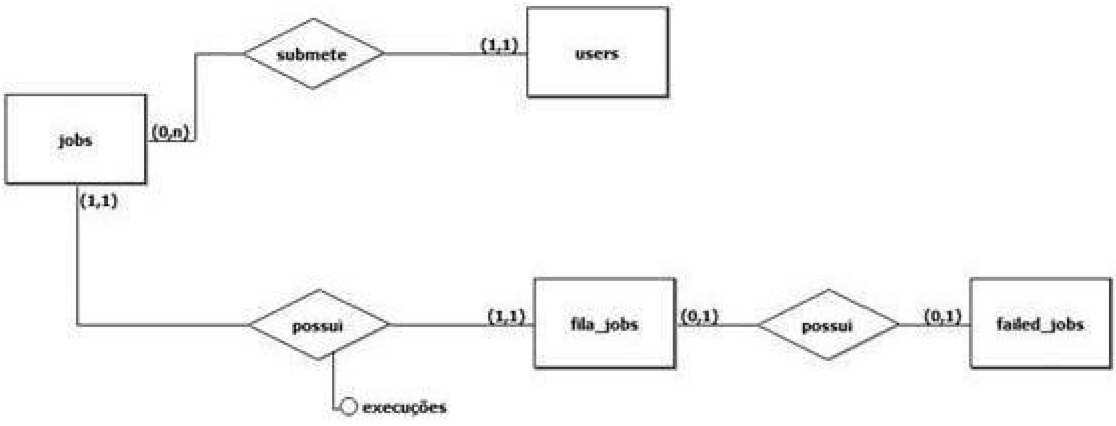

**Figura 10: Modelo Entidade-Relacionamento do sistema A-DOBRA.** O diagrama apresenta a estrutura lógica do banco de dados da aplicação, evidenciando as principais entidades, seus atributos e os relacionamentos utilizados para armazenar usuários, execuções, arquivos gerados e informações operacionais do sistema.
Fonte: Os autores, 2025.

O modelo também contempla o tratamento de falhas: uma execução na *fila_jobs* pode opcionalmente (0,1) estar associada a um registro em *failed_jobs*, que armazena informações sobre execuções que não foram concluídas com sucesso. Dessa forma, o diagrama evidencia tanto o fluxo normal de execução quanto o mecanismo de rastreabilidade de erros, garantindo controle, auditoria e consistência no gerenciamento dos *jobs* submetidos ao sistema (Figura 10).

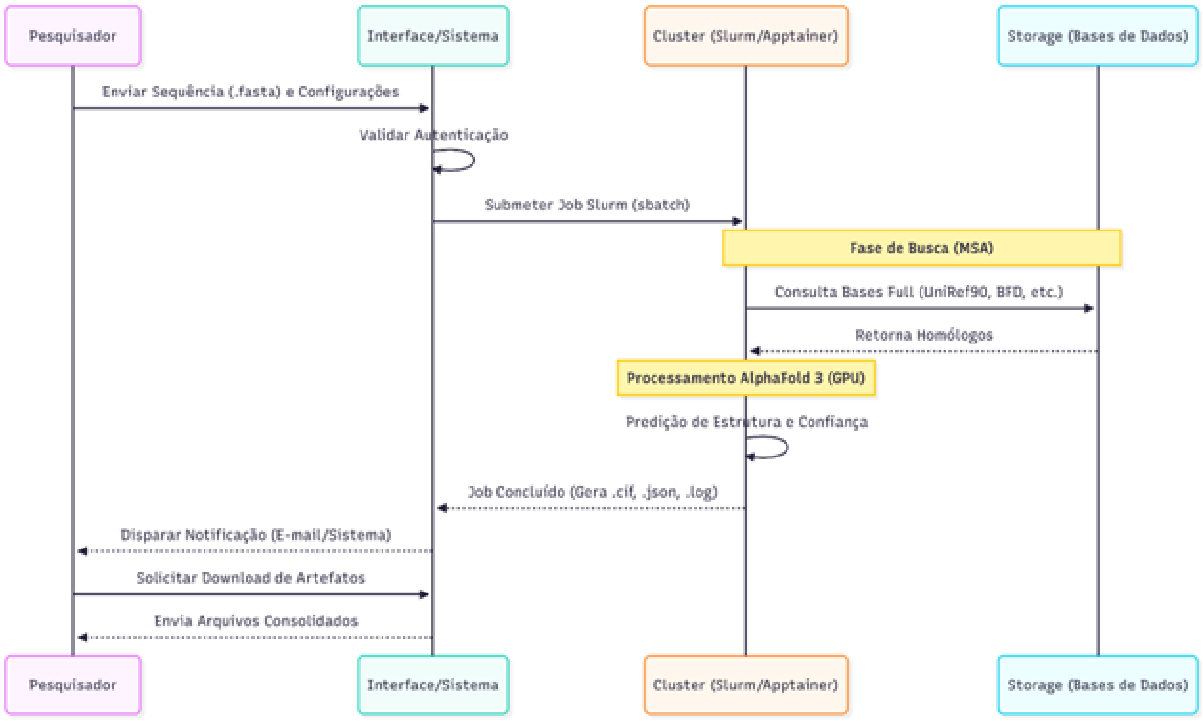

**Figura 11: Diagrama de Sequência com descrição dos passos e interações do usuário com o software.** A figura descreve o fluxo principal de uso da aplicação, apresentando as etapas da interação do usuário com o software e a sequência de processos executados para submissão, processamento e retorno das predições realizadas com o AlphaFold 3.
Fontes: Os autores, 2025.

O diagrama de sequência (Figura 11) ilustra bem os caminhos de execução do sistema A-DOBRA, evidenciando a interação entre o pesquisador, a interface web, o *cluster* HPC gerenciado pelo Slurm e o subsistema de armazenamento de dados. Com o processo iniciando com o envio da sequência biológica no formato FASTA, a interface, por sua vez, realiza a validação da autenticação e, em seguida, submete o *job* ao Slurm por meio do comando `sbatch`, acionando o pipeline de execução no cluster utilizando containers Apptainer.

A etapa inicial do processamento corresponde à geração do alinhamento múltiplo de sequências (MSA), na qual são consultadas bases de dados biológicas, como UniRef90 e BFD, para identificação de homólogos. Com os dados evolutivos obtidos, o AlphaFold 3 é executado com aceleração por GPU, realizando a predição da estrutura tridimensional e das métricas de confiança. Esta parte da execução segue o procedimento indicado pela distribuição.

Após a conclusão do *job*, os artefatos gerados (arquivos .cif, .json e logs) são persistidos e o usuário é notificado. Por fim, o pesquisador pode iniciar o download dos resultados consolidados por meio da interface do sistema. O dashboard que consiste em uma interface gráfica que organiza e exibe, de forma integrada, as principais informações e resultados das execuções (Figura 12). A Figura 13, por sua vez, traz a visualização da lista de arquivos gerados como output de um *job* enviado. O sistema está homologado e disponível preliminarmente para uso na infraestrutura do Laboratório de Computação Científica da UFMG (LCC - UFMG) pelo endereço: https://st01.lcc.ufmg.br/. A seguir o acoplamento de A-DOBRA com testes de estabilidade da predição serão descritos como evolução do sistema, acoplando dinâmica molecular de execução rápida como teste do modelo.

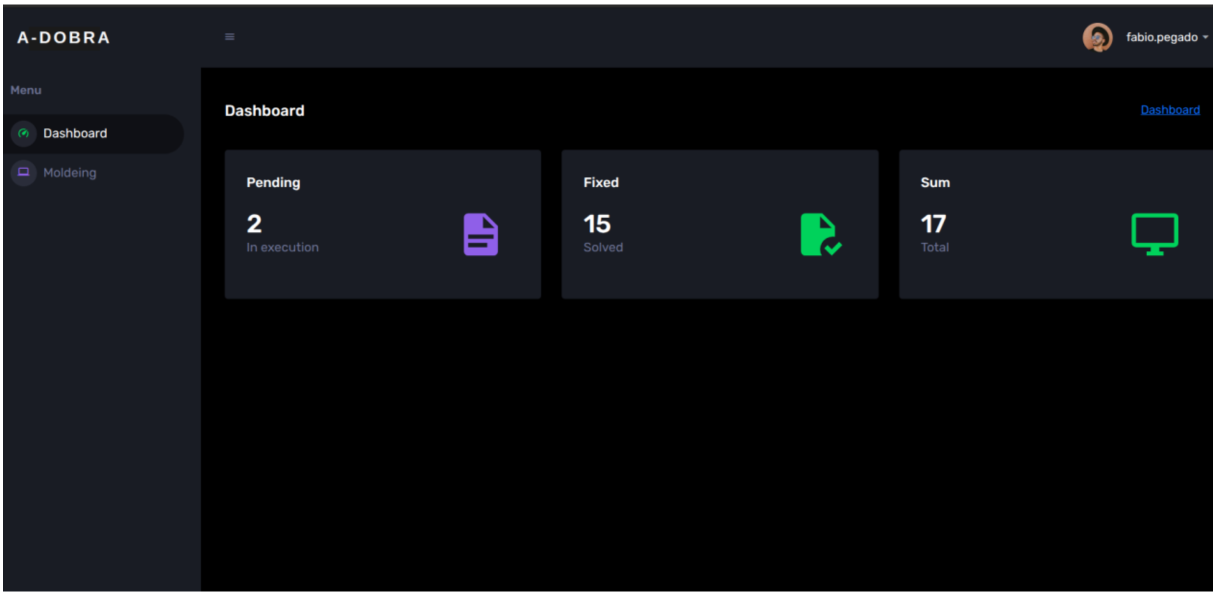

**Figura 12: Dashboard do sistema A-DOBRA.** A interface reúne, de forma centralizada, as principais informações das execuções, permitindo ao usuário acompanhar o status dos jobs, acessar resultados e gerenciar as predições realizadas no sistema.
Fonte: Os autores, 2025.

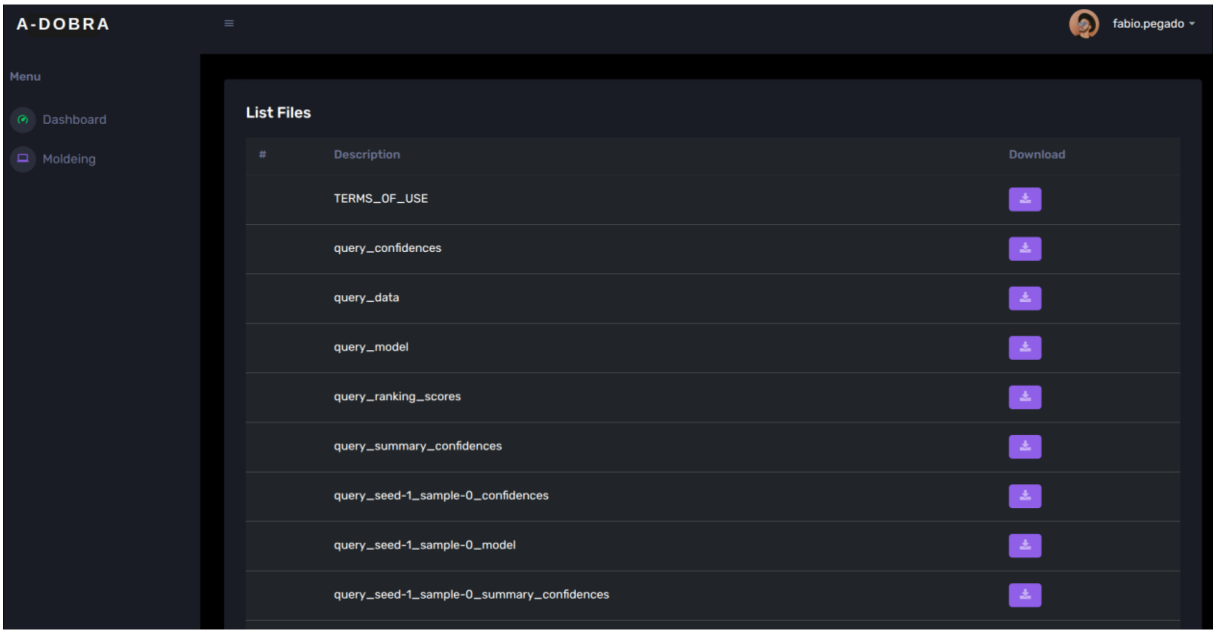

**Figura 13: Lista de Arquivos de Output gerados pelo Alphafold 3.** A interface apresenta os artefatos produzidos após a execução do pipeline, incluindo estruturas preditas, métricas de confiança, arquivos auxiliares e registros necessários para análise e download dos resultados.
Fonte: Os autores, 2025.

#### 5.2 AVALIAÇÃO DE DESEMPENHO E IMPACTO DA ARQUITETURA DE GPUS

O AlphaFold 3 apresenta um processamento mais rápido se comparado ao AlphaFold 2, utilizado inicialmente neste trabalho, com o uso menor de memória e um maior aproveitamento de CPU (Tabela 01). Essa performance deu base, para além dos testes iniciais, seguir com estudos subsequentes com o AlphaFold3.

**Tabela 01: Comparação entre as bases de dados**

| Característica | Base de Dados AlphaFold 2 | Base de Dados AlphaFold 3 |
|---|---|---|
| Principais Bancos | UniRef90, MGnify, BFD, Uniclust30, PDB70. | UniRef90, MGnify, BFD otimizado, PDB, Dados estruturais de interações biomoleculares |
| Tamanho em Disco | ~2.5 TB a 3.0 TB. | ~500 GB a 1.5 TB |
| Alinhamento (MSA) | Busca exaustiva em bilhões de sequências (BFD completo). | Uso otimizado de MSA combinado com informações estruturais e interacionais |
| Tempo de Execução | Muito maior (busca em BFD/Uniclust é lenta). | Mais rápido do que ambos do AF2. Otimizado para GPUs moderna, |
| Precisão (pLDDT) | Geralmente superior para proteínas raras ou novas. | Alta precisão, com melhorias em complexos e interações biomoleculares |
| Uso de Memória | Alta exigência de RAM para gerenciar índices. | Gerenciamento mais eficiente e modular |

Fonte: Os autores, 2025.

O script inicial executava uma modelagem por vez, em uma fila do Slurm chamada “filaGPU”, que contava com 14 máquinas com GPU NVIDIA GeForce RTX 3060. Como teste foi modelado a mesma sequência da proteína Faseolina (Figura 13) em 4 GPUs, tendo o tempo médio gasto para a modelagem da Faseolina de 1 hora e 40 minutos (Tabela 02). A disponibilidade deste modelo de GPU é bem comum em vários laboratórios, sendo uma GPU de baixo custo.

**Tabela 02: Execuções testes com o Apptainer e Slurm para modelagem de Faseolina.**

| Job ID | Node | CPUs | Execution Mode | Duration |
|---|---|---|---|---|
| 553 | gpu01 | 16 | GPU-accelerated | 0h 51m 50s |
| 562 | gpu01 | 16 | GPU-accelerated | 1h 55m 15s |
| 571 | gpu02 | 16 | GPU-accelerated | 1h 21m 10s |
| 592 | gpu03 | 16 | GPU-accelerated | 1h 25m 28s |

Fonte: Os autores, 2025.

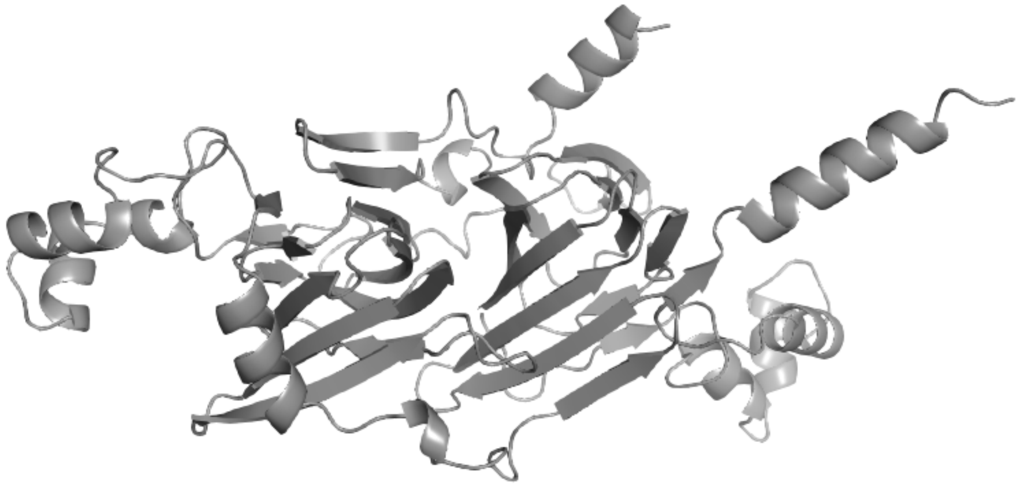

**Figura 14: Proteína Faseolina (PDB 2PV7) modela por AlphaFold 3. Estruturas secundárias apresentadas em *cartoon* pelo PyMOL.**
Fonte: Os autores, 2025.

A variação entre 51 minutos e 1h 55m é considerada um tempo alto, uma vez que usuários do contexto web procuram soluções mais rápidas, ou quase imediatas. Com isso, expandimos para uma máquina com maior poder de processamento, GPU NVIDIA A40 pois, no caso do AlphaFold 3, a execução ocorre de forma essencialmente sequencial e fortemente acoplada a uma única instância de GPU, não sendo eficiente dividir a tarefa em múltiplas GPUs com um menor poder de processamento (ABRAMSON *et al*., 2024) (Tabela 03). Dessa forma, o desempenho depende diretamente da capacidade de uma GPU individual mais robusta, resultando em um tempo total em torno de 15 minutos (Tabela 04). A fila orquestrada pelo Slurm controla a demanda por múltiplas execuções. Note-se que em seguida à execução de A-DOBRA adiante mostraremos seu acoplamento com dinâmica moléculas, logo o tempo de execução acelerado pela GPU A40 permite execução mais rápida e acoplamento com o passo seguinte.

**Tabela 03: Comparação de hardware entre as GPUS NVIDIA A40 e GeForce RTX 3060.**

| Característica | NVIDIA A40 | NVIDIA GeForce RTX 3060 |
|---|---|---|
| Arquitetura | Ampere | Ampere |
| Segmento | Datacenter / HPC | Desktop / Consumer |
| CUDA Cores | 10.752 | 3.584 |
| Tensor Cores | 336 (3ª gen) | 112 (3ª gen) |

| | | |
|---|---|---|
| Memória VRAM | 48 GB GDDR6 ECC | 12 GB GDDR6 |
| ECC | Sim | Não |
| Largura de banda | ~696 GB/s | ~360 GB/s |
| FP64 | Suporte limitado (HPC) | Muito limitado |
| TDP | 300 W | ~170 W |

Fonte: NVIDIA.

**Tabela 04. Tempo de processamento de modelagem da Faseolina (WT) e suas variantes mutantes por BLOSUM62. GPU NVIDIA A40.**

| **Modelo** | **Duração** |
|---|---|
| WT (2PV7) | 19m20s |
| F-1 | 17m13s |
| M0at | 15m55s |
| L-1 | 18m09s |
| I-1 | 19m02s |
| W-1I | 16m13s |
| Y-1 | 17m20s |
| V-1 | 16m03s |
| T0 | 15m07s |
| R0 | 16m28s |
| H0ks | 17m57s |
| K+n | 18m45s |
| C0tv | 19m01s |
| R-1 | 17m50s |
| R-2 | 18m08s |

Fonte: Os autores, 2025.

A redução do tempo de execução traz mais agilidade aos experimentos. Com execuções mais rápidas, é possível testar mais casos em menos tempo, comparar mais resultados e ajustar os experimentos de forma mais dinâmica, tornando os resultados ainda mais robustos.

Toda a infraestrutura computacional utilizada foi disponibilizada pelo Laboratório de Computação Científica da UFMG (LCC – UFMG.

#### 5.3 VALIDAÇÃO DO PIPELINE *AlphaUnFold*

Esta seção apresenta os experimentos e estudos de caso realizados com o pipeline AlphaUnFold (Figura 15). Com foco na avaliação do comportamento estrutural das proteínas preditas com A-DOBRA (alphafold3), os resultados abrangem análises de correlação entre métricas e estudos com modificações direcionadas na sequência, permitindo investigar de forma integrada a relação entre confiança preditiva e estabilidade estrutural. O sistema consolidado, bem como seus scripts de automação, encontra-se disponível no repositório: https://github.com/pegados/pipeline_AlphaUnfold. Desta forma, para execução de uma quantidade significativa de modelos, os interessados podem fazer sua própria instalação e execução, até que tenhamos possibilidade de execução via web em larga escala. O pipeline de AlphaUnFold é portanto mais detalhado e complexo, entregando o acoplamento de predição e dinâmica molecular.

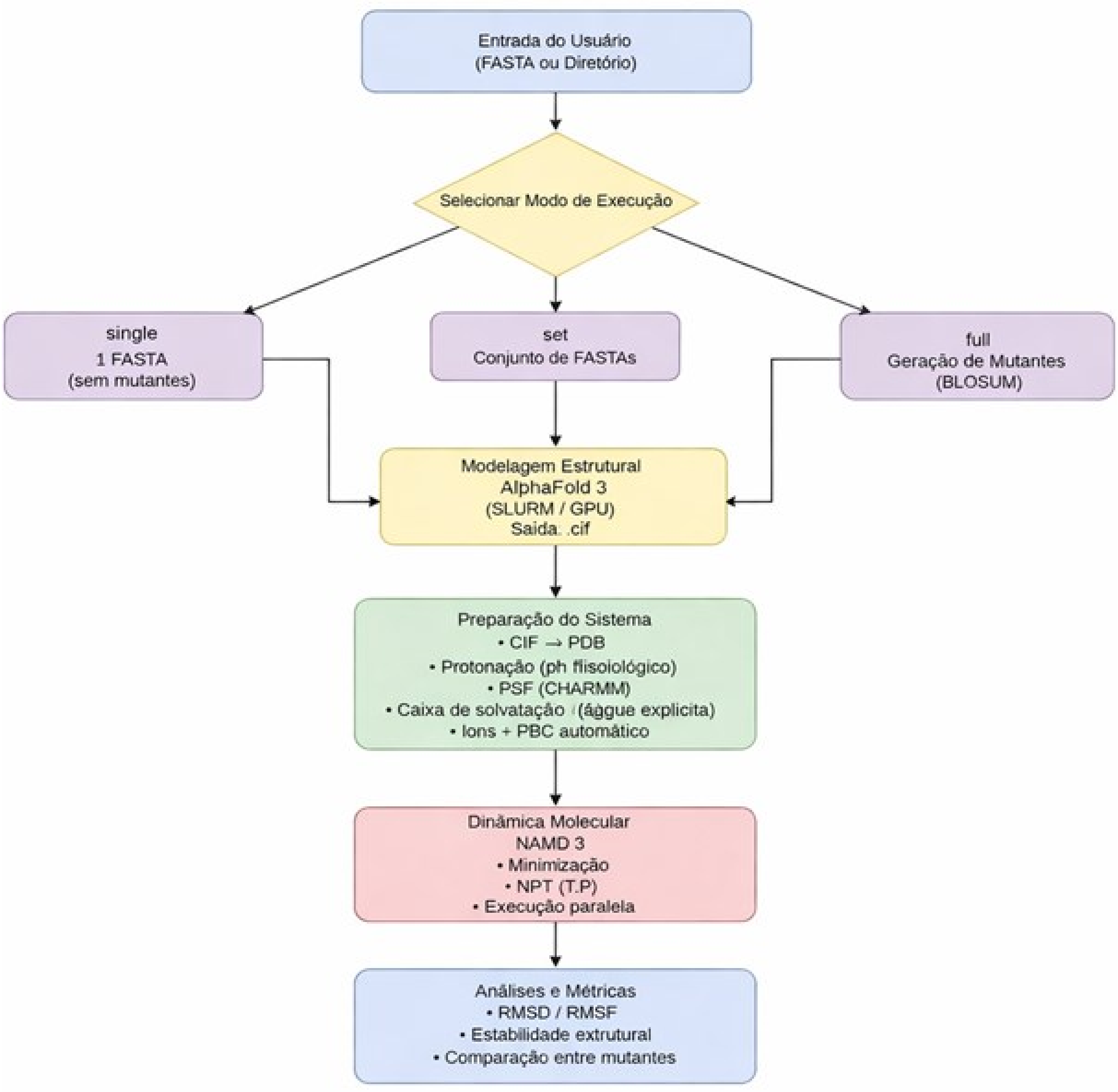

**Figura 15: Fluxograma do Pipeline AlphaUnFold.** O diagrama apresenta as etapas automatizadas do pipeline, desde a entrada da sequência proteica e geração do modelo estrutural pelo AlphaFold até a preparação, execução das simulações de dinâmica molecular e obtenção das métricas utilizadas nas análises estruturais.
Fonte: Os autores, 2025.

##### 5.3.1 ESTUDO DE CASO PARA CORRELAÇÕES DO PLDDT

Proteínas foram selecionadas aleatoriamente no site EBI EMBL Alphafold para o estudo de predição local e inferência de estabilidade sob simulação de dinâmica molecular e (Tabela 06) foram organizadas em ordem crescente de pLDDT médio, com o objetivo de abranger um espectro que vai desde modelos previstos com baixa confiança estrutural até estruturas altamente confiáveis segundo o AlphaFold. Essa organização permitiu avaliar sistematicamente como a confiança preditiva local se relaciona com métricas dinâmicas globais e locais pela execução de dinâmica molecular com o software NAMD3.

**Tabela 06: Identificação proteica, seus valores de pLDDT e número de resíduos de aminoácidos.**

| ID | pLDDT | nº de aminoácidos |
|---|---|---|
| Q14236 | 35,72 (Very Low) | 149 |
| Q9ULZ0 | 40,28 (Very Low) | 124 |
| O15503 | 68,25 (Low) | 277 |
| P04637 | 75,06 (High) | 393 |
| Q96M98 | 76,44 (High) | 296 |
| Q96M98-2 | 85,06 (High) | 257 |
| P0DP23 | 85,25 (High) | 149 |
| Q6NUM9-2 | 89,75 (High) | 481 |
| Q6NUM9 | 95,12 (Very High) | 610 |

Fonte: Os autores, 2025.

Em nosso grupo vinham sendo desenvolvidas simulações de dinâmica molecular com proteínas desestabilizadas por mutações crescentes de trocas dos resíduos por aminoácidos essenciais. De posse de moléculas que eram instabilizadas em níveis dose-efeito em função das trocas, havia sido feita a padronização de simulação de dinâmica molecular sob alta pressão ou temperatura, sendo escolhido o protocolo sob 1000 atm em temperatura normal. As mudanças na estabilidade estrutural da proteína que ocorriam em simulações nas CNTP por 200 a 500 ns eram obtidas com simulações de 5 ns a 1000 atm. Portanto, decidimos acoplar ao A-DOBRA uma automação da realização de simulação de dinâmica molecular automatizada, criando uma aplicação que faz a rpedição e testa a estabilidade da predição, denominada AlphUnFold. Os resultados a seguir mostram que em simulações curtas de 5 ns sob alta pressão, já é possível identificar padrões consistentes entre a confiança preditiva do AlphaFold e o comportamento estrutural das proteínas, surgindo como uma alternativa prática para realizar um ajuste fino inicial nos modelos gerados pelo programa, permitindo avaliar sua estabilidade em tempo computacional reduzido. Os resultados da Figura 16 foram obtidos por execução com orquestrador Slurm para as distints proteínas, mostrando-se uma relação entre a métrica RMSD após uma simulação curta de 5 ns sob alta pressão ou comumente realizada de 200 ns em pressão de 1 atm.

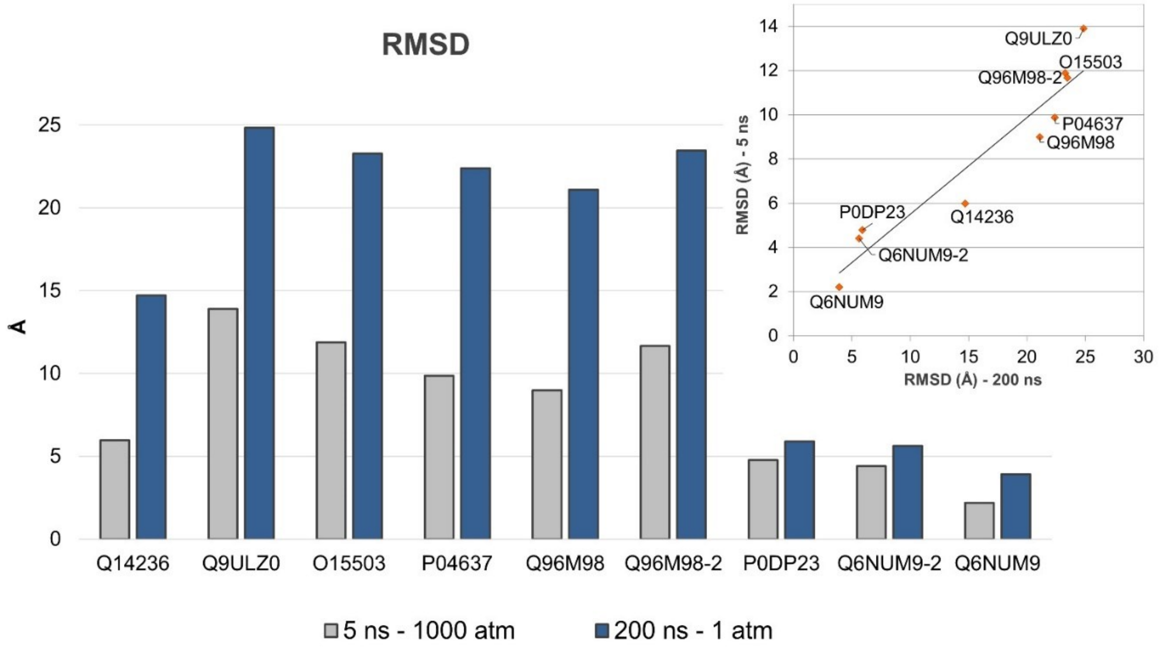

**Figura 16: Valores de RMSD médio dos últimos 20% ns finais de 5 ns (cinza) e 200 ns (azul). No inset relação entre ambos RMSD obtidos.**
Fonte: Os autores, 2025.

A análise de correlação entre o pLDDT médio e o RMSD global ao longo de simulação de 5 e 200 ns (Figuras 17 e 18) revelam que estruturas previstas com maior confiança (pLDDT alto) pelo AlphaFold tendem a apresentar maior estabilidade durante a dinâmica molecular e os de menor confiança (pLDDT baixo) teriam uma menor estabilidade, porém nem sempre essa correlação se confirma, o que está representado pelos “outliers” em vermelho. A relação inversa suporta o uso da aplicação AlphaUnFold e a presença de “outliers” indica a importância da verificação experimental, muito embora “in silico”, da estabilidade.

Todavia, métricas locais não predizem o RMSD da proteína Q96M98-2, que apresentou pLDDT médio de aproximadamente 85%, o que, pela regressão linear proposta (RMSD = 35 − 0,34·pLDDT), estimaria um RMSD de ~6,1 Å. No entanto, a simulação revelou RMSD global de 11,7 Å. Diferenças entre confiança preditiva e estabilidade dinâmica também podem decorrer da ausência de restrições pós-traducionais, como pontes dissulfeto entre resíduos de cisteína. Como essas ligações requerem tratamento explícito na parametrização da dinâmica molecular, sua ausência pode contribuir para aumentos artificiais de RMSD mesmo em modelos com pLDDT elevado.

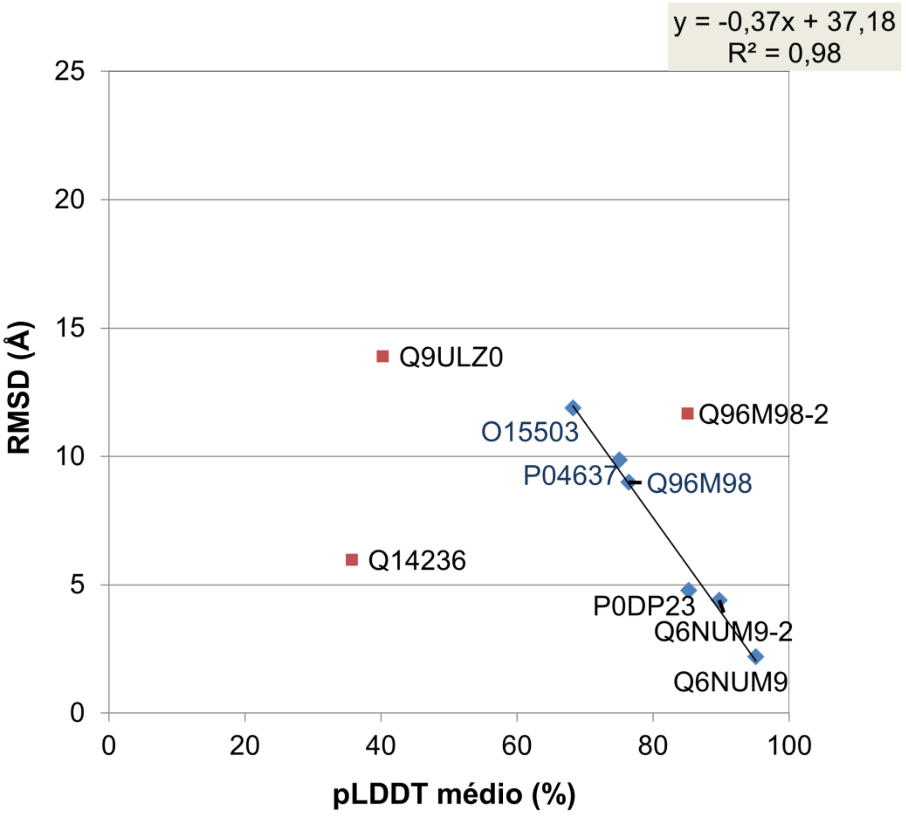

**Figura 17: Correlação entre RMSD e pLDDT médio. Simulações de 5 ns a 1000 atm.**
Fonte: Os autores, 2025.

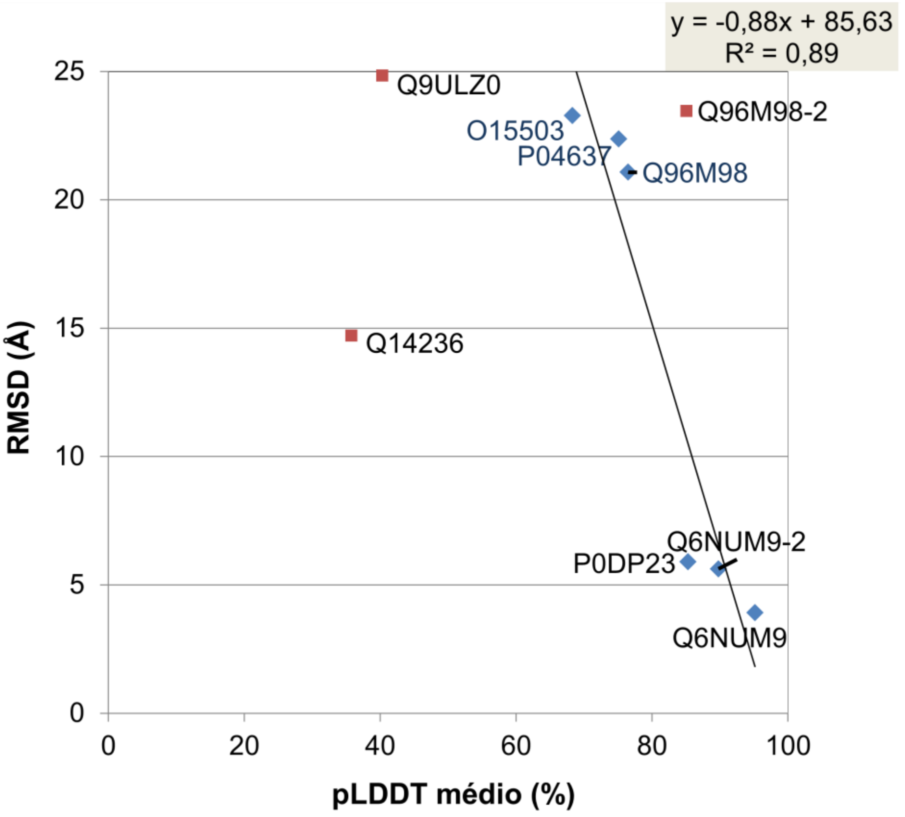

**Figura 18: Correlação entre RMSD e pLDDT médio. Simulações de 200 ns a 1 atm.**
Fonte: Os autores, 2025.

A análise de correlação inversa entre o pLDDT médio e o RMSF em simulações de 5 ns a 1000 atm e de 200 ns a 1 atm (Figuras 19 a 21 pela ordem de menor ao maior valor de pLDDT) revelam que estruturas previstas com alta confiança (pLDDT próximo a 100%), o RMSF manteve-se em níveis mínimos. Isso demonstra que os segmentos identificados pelo modelo como estruturados são, de fato, as regiões de menor mobilidade na simulação física. Isso reforça que as predições de alta confiança não são apenas geometricamente coerentes, mas representam núcleos estruturais robustos, capazes de resistir às flutuações térmicas da simulação. A ferramente desta forma não somente contribui com os modelos previstos, mas com teste rápido compatível com execução em portal de aplicação para os modelos.

Uma forma de avaliar posição a posição a correlação inversa entre valor de pLDDT e movimentação da posição durante a dinâmica molecular é o RMSF de cada resíduo. Na Figura 19 estão representados esses dois valores e observamos correlação inversa, onde domínios com excelente predição coabitam com regiões de baixo RMSF, sendo esta uma proteína bastante ilustrativa por prover segmentos consecutivos de alta e baixa qualidades de predição. Note que a implementação de baixo custo de 5 ns apenas sob alta pressão mostra os mesmos domínios estáveis ou instáveis que a simulação mais demorada em condições normais.

O mesmo é obtido para outra proteína, cujos resultados se encontram na Figura 20. Desta forma, não só há uma boa correlação entre os valores médios de RMSF e pLDDT, como o comportamento local é acompanhado. Com a programação executada neste trabalho, fazemos a execução automarizada de alphafold3, preparação da estrutura para simulação de dinâmica molecular com NAMD3 e obtenção dos resultados, na aplicação conjugada AlphaUnFold.

A proteína ensaiada na Figura 21 apresenta um domínio com baixa capacidade de predição, e é justamente este que obtém os resultados piores de RMSF, mostrando que a alta qualidade de predição se reflete em estrutura não só bem predita mas estável, ressaltando a contribuição da aplicação AlphaUnFold.

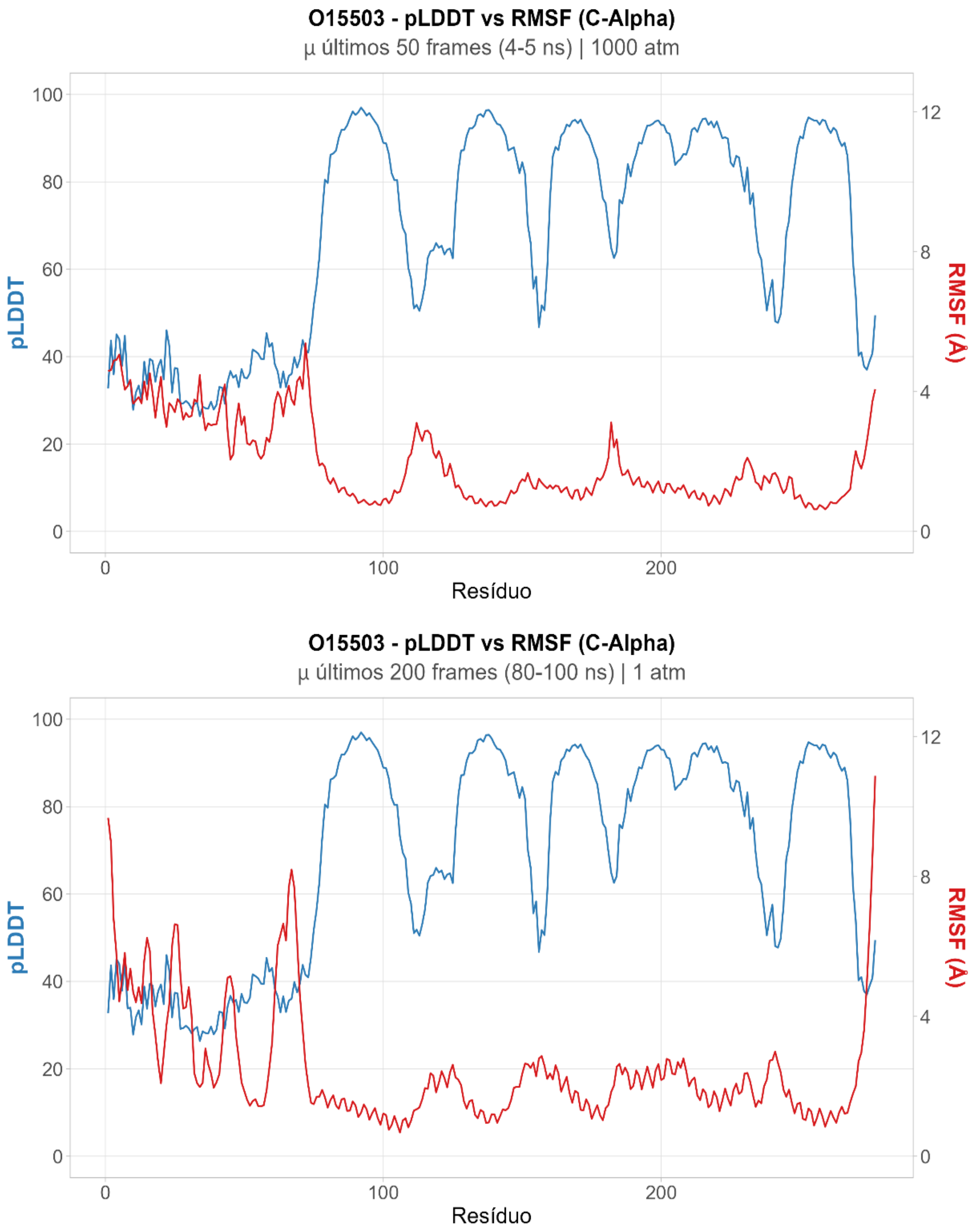

**Figura 19: RMSF x pLDDT médio - 015503. Comparações entre de 5 e 200 ns**.
Fonte: Os autores, 2025.

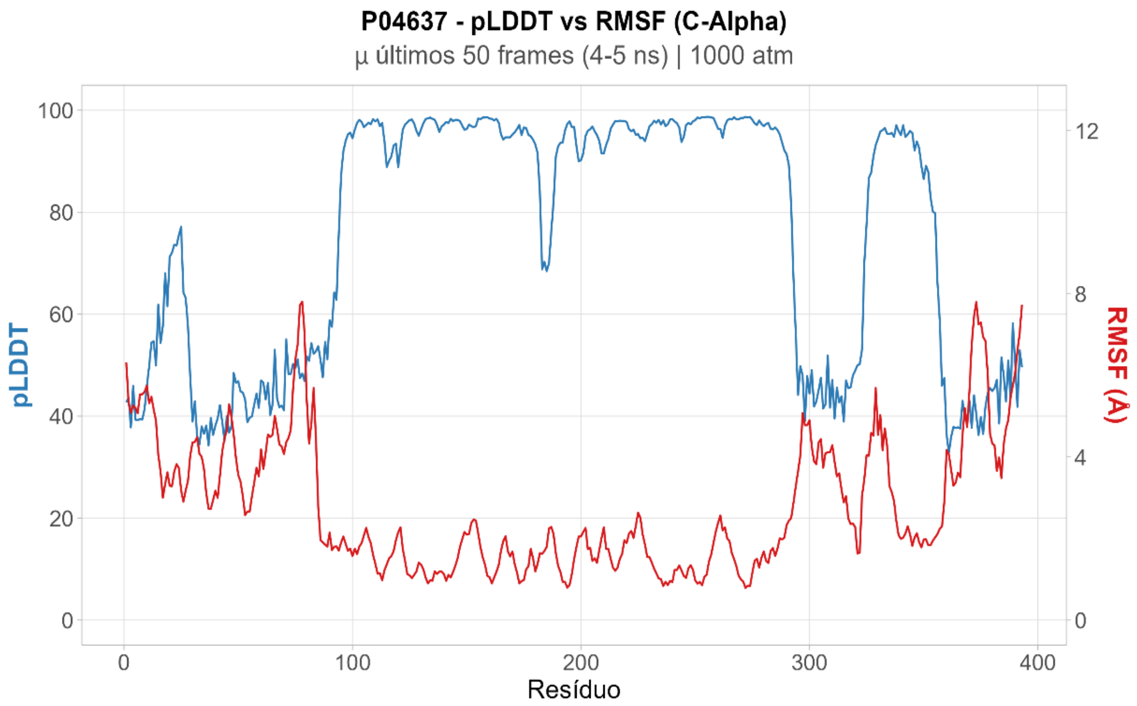

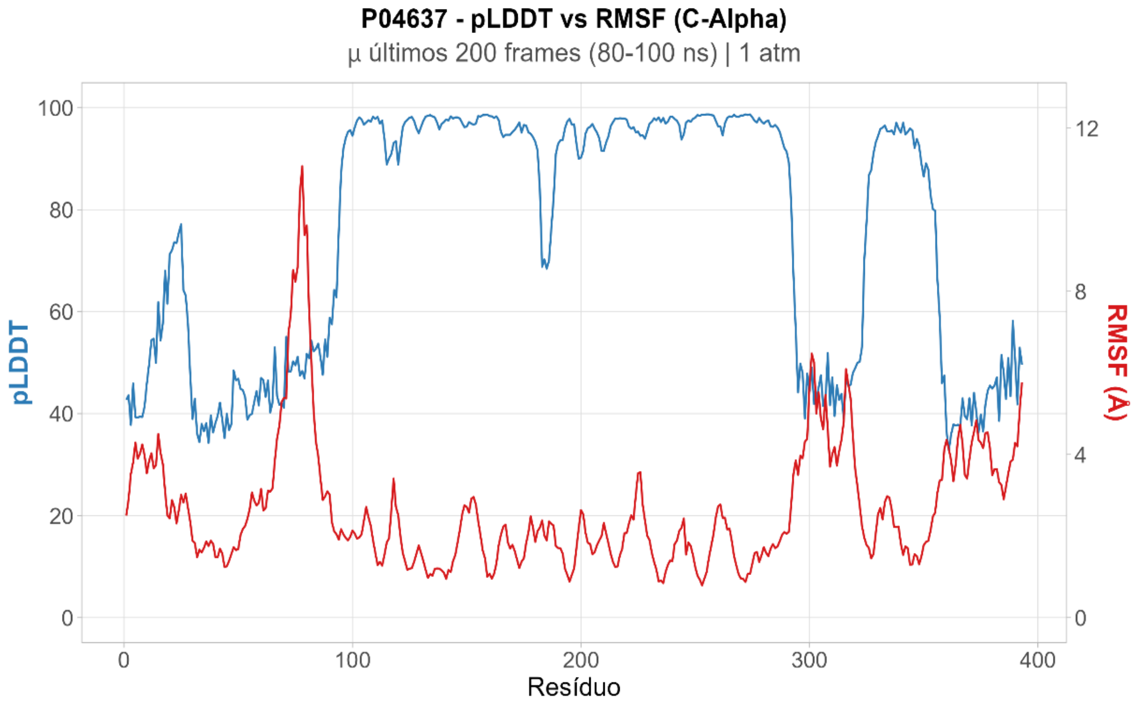

**Figura 20: RMSF x pLDDT médio – P04637. Comparações entre de 5 e 200 ns.**
Fonte: Os autores, 2025.

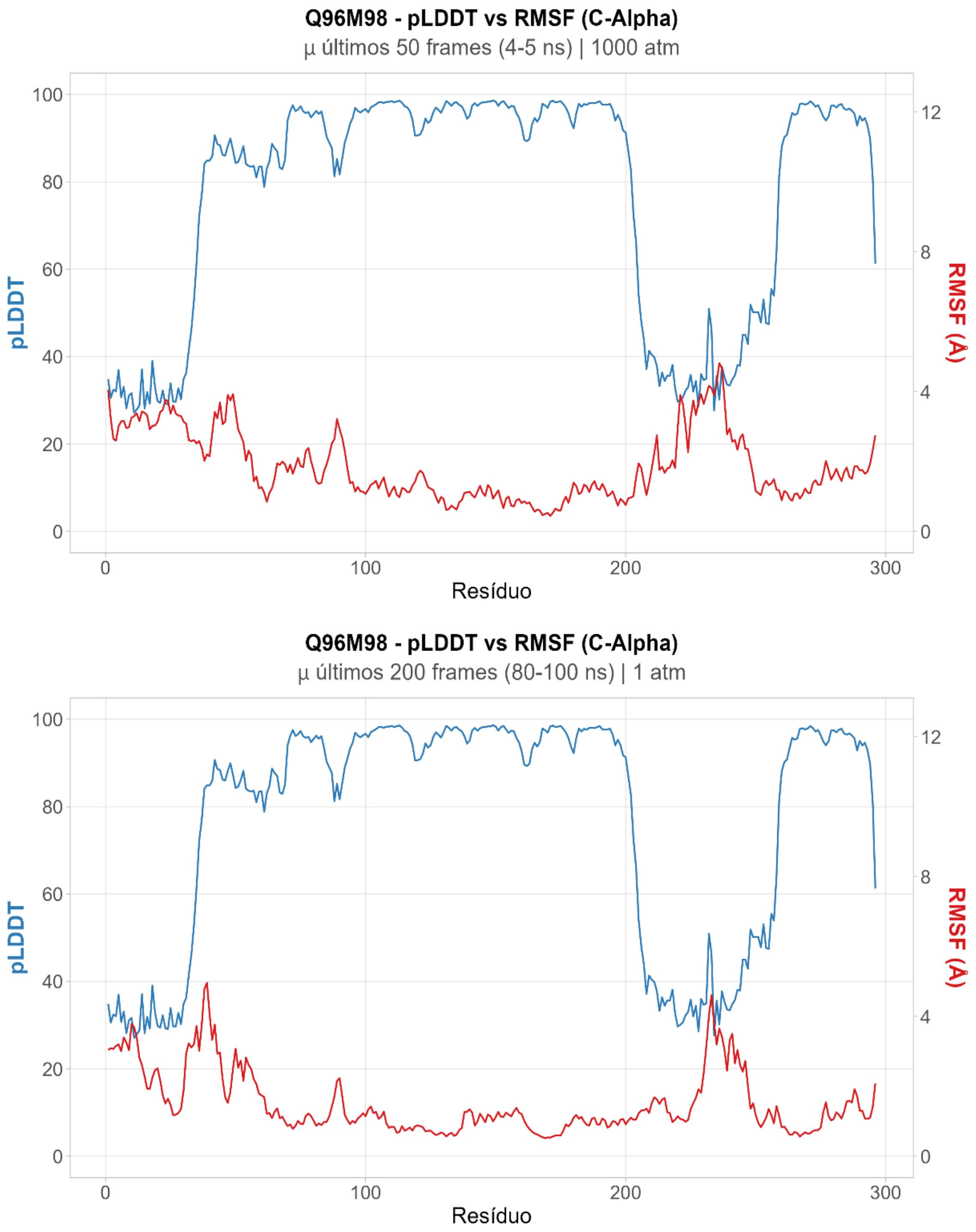

**Figura 21: RMSF x pLDDT médio – Q96M98. Comparações entre de 5 e 200 ns.**
Fonte: Os autores, 2025.

Os resultados mostram variações consistentes entre as proteínas, com valores de RMSD acompanhando, em geral, o nível de confiança preditiva. Esses resultados validam a premissa central da nossa metodologia. Se o AlphaFold fornece a "hipótese" estrutural através do pLDDT, a Dinâmica Molecular atua como a prova de estresse físico. A relação inversa entre as curvas confirma que a confiança local do modelo é um preditor excelente da estabilidade dinâmica real.

##### 5.3.2 ESTUDO DE CASO COM A MIOGLOBINA E SUAS MUTANTES

Em trabalho em colaboração em nosso grupo, objetiva-se enriquecer proteínas com substituição de resíduos por determinados aminoácidos. A experiência do grupo com substituições que por dose-efeito desestabilizavam a proteína nos motivou a questionar se regiões de baixa predição tinham dificuldade em se manter estruturadas. Logo, dispúnhamos do conhecimento de que trocas crescentes de resíduos naturais da proteína em um dado momento criariam instabilidade e utilizamos este modelo para estudo de caso da aplicação AlphaUnFold. A mioglobina 5XL0 é ma proteína modelo globular e é composta por 151 resíduos de aminoácidos (Tabela 07). Suas substituições progressivas por resíduos por arginina (R) em mioglobinas mutantes manteve valores baixos de RMSD (Figura 22) mesmo com até aproximadamente 40% de substituições por arginina, indicando preservação da estrutura global apesar das perturbações introduzidas até o tempo de simulação testado. Devido à carga positiva e alta polaridade, a arginina tende a alterar o empacotamento interno e as interações eletrostáticas da proteína, funcionando como um agente de perturbação estrutural. As trocas eram feitas em dose-efeito respeitando-se a inferência do efeito em trocas individuais onde era mendido o ΔΔG predito por cada troca por uma aplicação externa mCSM (Pires *et al.*, 2013). Assim eram detectadas trocas até estabilizadoras, passando por neutras e posteriormente desestabilizadoras. Faz sentido portanto, trocar decis ou quantidades crescentes priorizando as trocas menos desestabilizantes. Trocas nas “piores posições” (valores menores de ΔΔG) foram usadas como controle negativo de estabilização. Níveis mais elevados de mutação (quando são selecionadas posições energeticamente desfavoráveis) utilizadas como controle negativo, observa-se um aumento acentuado do RMSD (Figura 22), evidenciando perda de estabilidade conformacional.

Esses resultados indicam que a proteína apresenta uma resiliência estrutural inicial, seguida por um limite a partir do qual pequenas alterações adicionais resultam em grandes desvios estruturais. Todavia, apontam a possibilidade de trocas até uma quantidade significativa. A avaliação automatizada e rápida dessas possibilidades só foi possível com a aplicação AlphaUnFold, atuando na predição local e pronta com Alphafold3, em substituição a uma demorada modelagem com I-Tasser feita por servidor que retornava a modelagem por e-mail. Além disso, a preparação e execução da simulação com NAMD4 é feita automaticamente pela ferramenta, com preparação da molécula, etc. E com a utilização da dinâmica em alta pressão, validade pelos estudos acima, o resultado é obtido em menor tempo e com facilidade.

**Tabela 07: Composição da proteína mioglobina, PDB 5XL0. Obtida via ferramenta ProtParam (ExPASy).**

| Resíduo | n° de aminoácidos | % de aminoácidos |
|---|---|---|
| A | 17 | 11,3 |
| R | 4 | 2,6 |
| N | 1 | 0,7 |
| D | 7 | 4,6 |
| C | 0 | 0 |
| Q | 4 | 2,6 |
| E | 14 | 9,3 |
| G | 10 | 6,6 |
| H | 12 | 7,9 |
| I | 9 | 6 |
| L | 18 | 11,9 |
| K | 19 | 12,6 |
| M | 2 | 1,3 |
| F | 6 | 4 |
| P | 4 | 2,6 |
| S | 6 | 4 |
| T | 5 | 3,3 |
| W | 2 | 1,3 |
| Y | 3 | 2 |
| V | 8 | 5,3 |

Fonte: Os autores.

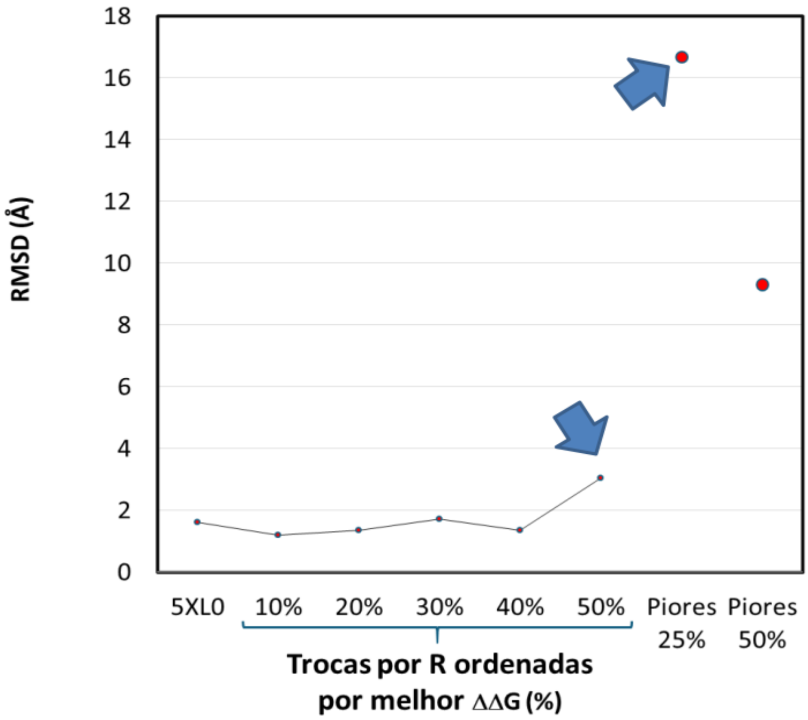

**Figura 22: Variação de RMSD da mioglobina (5XLO) em função da proporção de substituições por resíduos de arginina (R) via mCSM.**
Fonte: Os autores.

A ferramente além de RMSD extrai também os parâmetros raio de giro e área de exposição ao solvente (SASA), importantes para este projeto. Os resultados de raio de giro (RoG) e SASA (Figuras 23 e 24) reforçam o padrão observado anteriormente. Em níveis iniciais de substituição por arginina, o raio de giro varia pouco, indicando manutenção da compactação global da proteína. A partir de cerca de 40% de mutações, observa-se um aumento mais evidente, sugerindo início de expansão estrutural. Em paralelo, o SASA cresce de forma progressiva já em níveis intermediários, entre 20% e 30%, indicando aumento da exposição ao solvente antes de grandes alterações estruturais globais. Nos cenários mais desfavoráveis, utilizados como controle negativo, ambos apresentam elevações acentuadas, caracterizando descompactação e perda de organização estrutural.

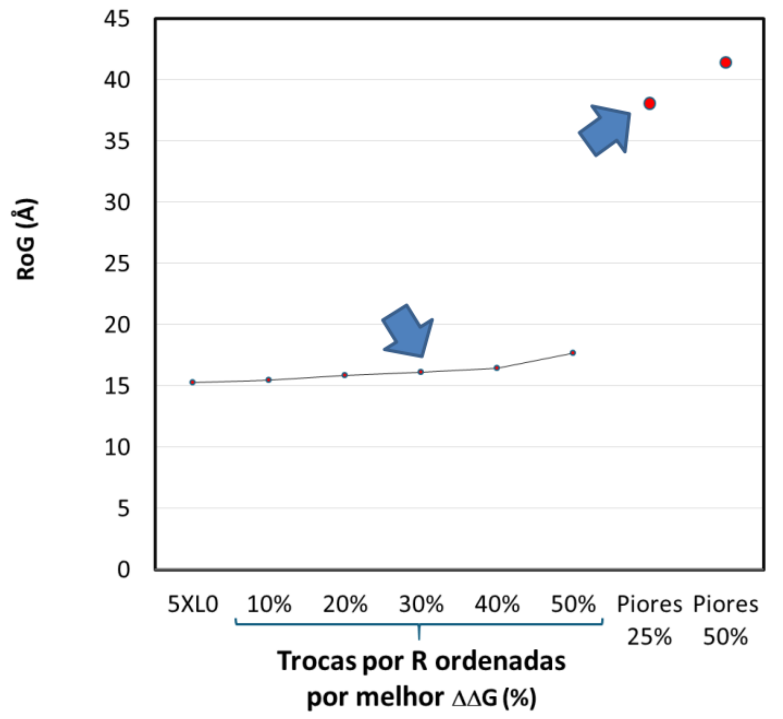

**Figura 23: Variação do raio de giro da mioglobina (5XLO) em função da proporção de substituições por arginina.**
Fonte: Os autores, 2025.

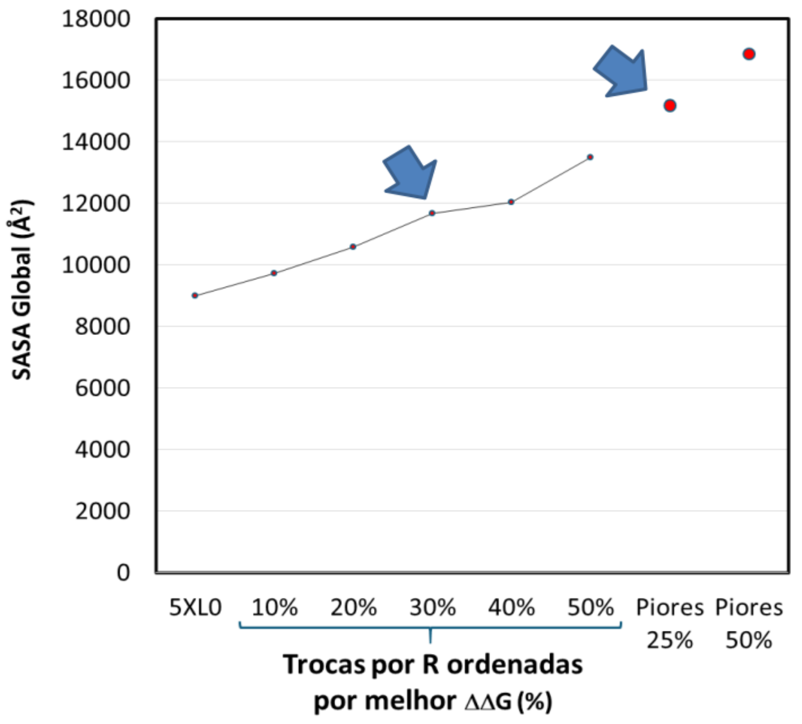

**Figura 24: Valores de SASA da mioglobina (5XLO) em função da proporção de substituições por arginina.**
Fonte: Os autores, 2025.

A análise do RMSF por resíduo ao longo dos decis de substituição por arginina mostra que a mioglobina mantém um perfil de flutuação próximo ao estado nativo até cerca de 30–40% de mutações (Figura 25). Nesse intervalo, as curvas permanecem semelhantes à estrutura original, indicando que a dinâmica local é pouco afetada mesmo com alterações distribuídas na sequência. Nos decis mais altos, em 40% e 50%, surgem aumentos pontuais de oscilação, sugerindo maior sensibilidade de regiões específicas às perturbações.

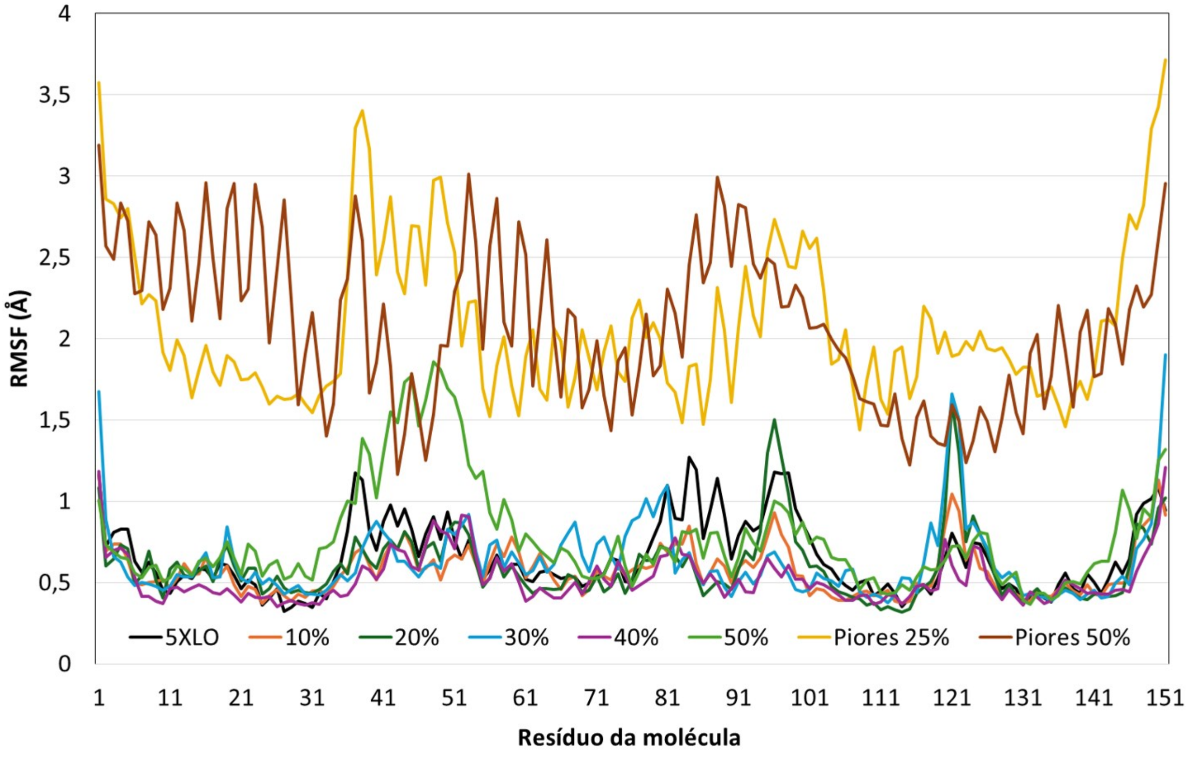

**Figura 25: Perfil de RMSF por resíduo da mioglobina (5XLO) ao longo dos decis de substituição por arginina.**
Fonte: Os autores, 2025.

Nos decis mais desfavoráveis, utilizados como controle negativo, o comportamento se altera de forma evidente. Observa-se aumento generalizado do RMSF e maior irregularidade ao longo da cadeia, indicando perda de coesão dinâmica e maior mobilidade estrutural. Esse contraste mostra que a estabilidade não depende apenas da quantidade de substituições, mas também de sua localização. Assim, a mioglobina tolera mutações nos decis mais favoráveis, enquanto apresenta desestabilização progressiva nos decis mais altos e, de forma mais intensa, quando as trocas ocorrem em posições críticas.

##### 5.3.3 ESTUDO DE CASO COM A FASEOLINA E SUAS MUTANTES

Um outro estudo de caso demonstrando a participação da aplicação AlphaUnFold em projeto de modelagem ou predição estrutural seguido de avaliação de estabilidade ou instabilidade estrutural está mostrado a seguir.

A Faseolina PDB 2PV7 é uma proteína de reserva do feijão, a qual pode ser enriquecida por substituição com aminoácidos essenciais ou semi-essenciais mas há uma demanda para execução de predição estrutural e teste de estabilidade em grupo, que pode ser suprida pela ferramenta AlphaUnFold. Ela é composta por 298 resíduos de aminoácidos e o teor dos mesmos na proteína estão mostrados na Tabela 08. A ferramenta AlphaUnFold é capaz de realizar trocas por valores de escore da tabela Blosum62, trocando primeiramente resíduos conservados em homólogos (valores positivos), valor zero onde a chance de encontrar a troca em um homólogo é igual ao esperado ao acaso, ou até valores negativos como -1, onde a chance de encontrar a troca em homólogo é um pouco inferior ao esperado aleatoriamente.

Em trabalho colaborativo no laboratório trocas com esse critério laboriosas foram feitas com a mioglobina como proteína modelo, mas agora com o AlphaUnFold poderíamos inferir rapidamente se as trocas testadas com mioglobina poderiam ser feitas com a faseolina do feijão. Gráfico de RMSD (Figura 26) mostra que a maioria das mutantes mantém valores de RMSD em torno de 4,2 angstrons (Å). A proteína selvagem (WT) tem o menor valor, 3,04 Å. A mutante que mais se aproxima do seu valor é a W-1L (3,08), seguida da Y-1 (3,37), M0at (3,76), I-1 (3,94) e K+n (3,97 Å). As próximas mutantes obtiveram valores maiores que 1 Å do valor da WT, H0ks (4,10), T0 (4,14), F-1 (4,23), C0tv (4,31) e V-1 (4,97 Å). As mutantes R-1 (5,91) um dos controles negativos e L-1 (6,32) apresentam os maiores valores, acompanhado do segundo controle negativo, R-2 (14,17 Å).

Os controles negativos utilizados são a troca por arginina até valores -1 e -2 apresentados na Blosum62, o que já sabíamos por resultados anteriores que eram trocas abusivas para manter a estabilidade. Os resultados prontamente suportaram a possibilidade de usar os modelos de troca planejados com a mioglobina, para a faseolina, exceto talvez para Leucina, onde trocar resíduos com escore -1 parece excessico, tendo que ser este enriquecimento refinado e talvez trocar para até escore zero apenas, enriquecendo um pouco menos a faseolina para leucina. Mesma coisa para Valina. Ressaltamos que todo o experimento foi feito por um único disparo sbatch utilizando Slurm.

**Tabela 08: Composição da proteína Faseolina, PDB 2PV7. Obtida via ferramenta ProtParam (ExPASy).**

| Resíduo | n° de aminoácidos | % de aminoácidos |
|---|---|---|
| A | 27 | 9,1 |
| R | 15 | 5,0 |
| N | 15 | 5,0 |
| D | 17 | 5,7 |
| C | 1 | 0,3 |
| Q | 15 | 5,0 |
| E | 21 | 7,0 |
| G | 18 | 6,0 |
| H | 8 | 2,7 |
| I | 22 | 7,4 |
| L | 36 | 12,1 |
| K | 14 | 4,7 |
| M | 8 | 2,7 |
| F | 14 | 4,7 |
| P | 8 | 2,7 |
| S | 15 | 5,0 |
| T | 13 | 4,4 |
| W | 4 | 1,3 |
| Y | 12 | 4,0 |
| V | 15 | 5,0 |

Fonte: Os autores.

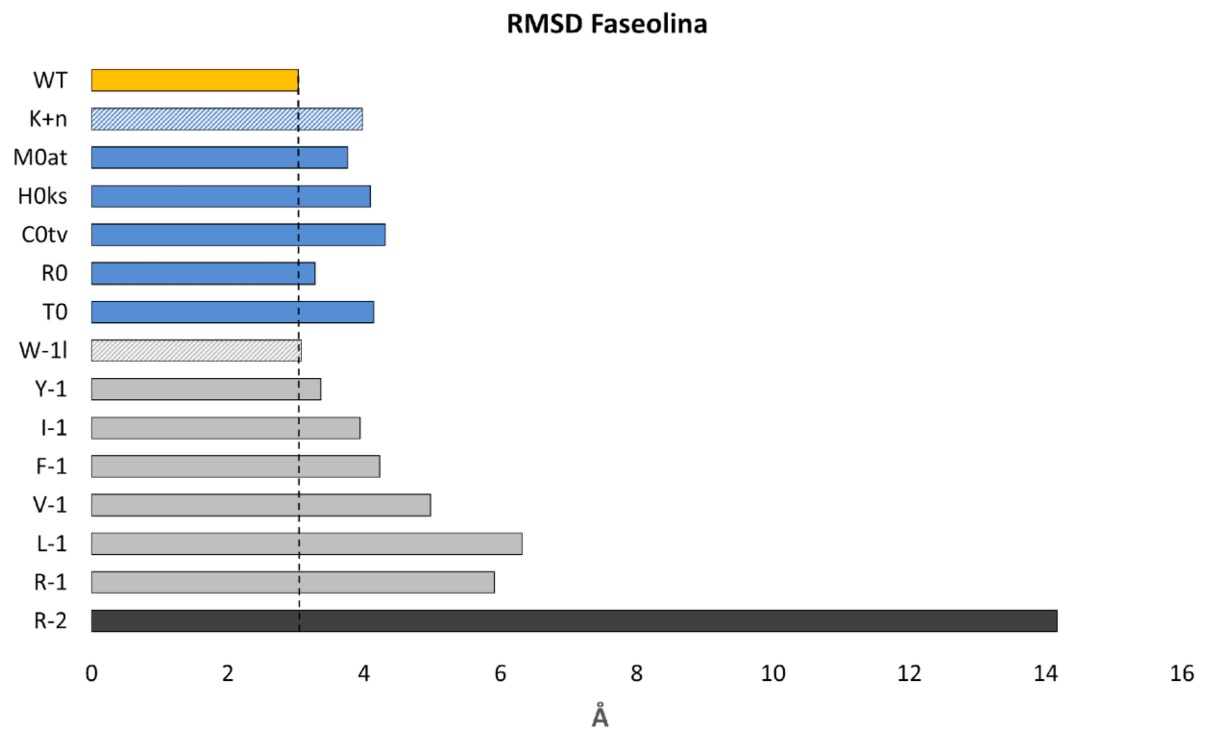

**Figura 26: Valores de RMSD da Faseolina (WT) e suas mutantes. A linha tracejada indica o valor médio de RMSD da WT como referência.**
Fonte: Os autores, 2026.

Portanto, pelos resultados prontamente obtidos, os resultados mostrados na Figura 26 (RMSD Faseolina), a mutação L-1 (substituições por leucina) merece atenção, pois apresentou RMSD elevado e ficou bem acima da proteína selvagem. Isso sugere que o aumento de leucina pode ter alterado o empacotamento interno da estrutura ou desestabilizado interações locais durante a simulação. Como se trata de um resíduo hidrofóbico e volumoso, trocas em posições inadequadas podem favorecer rearranjos conformacionais. Por isso, esse caso merece análise mais detalhada em estudos futuros, assim como algumas substituições por Valina, pela análise de RMSD. Como veremos abaixo, outros parâmetros são menos sensíveis às trocas excessivas por Leucina e Valina.

A acessibilidade ao solvente (SASA) (Figura 27) para a maioria das variantes mantém valores próximos ao da proteína selvagem (23281 Å). Os maiores valores foram para R0 (28020), R-1 (35271) e R-2 (51982 Å). O raio de giro apresentou valores muito semelhantes (Figura 28) variando de 24 (V-1) a 28 Å (R-1).

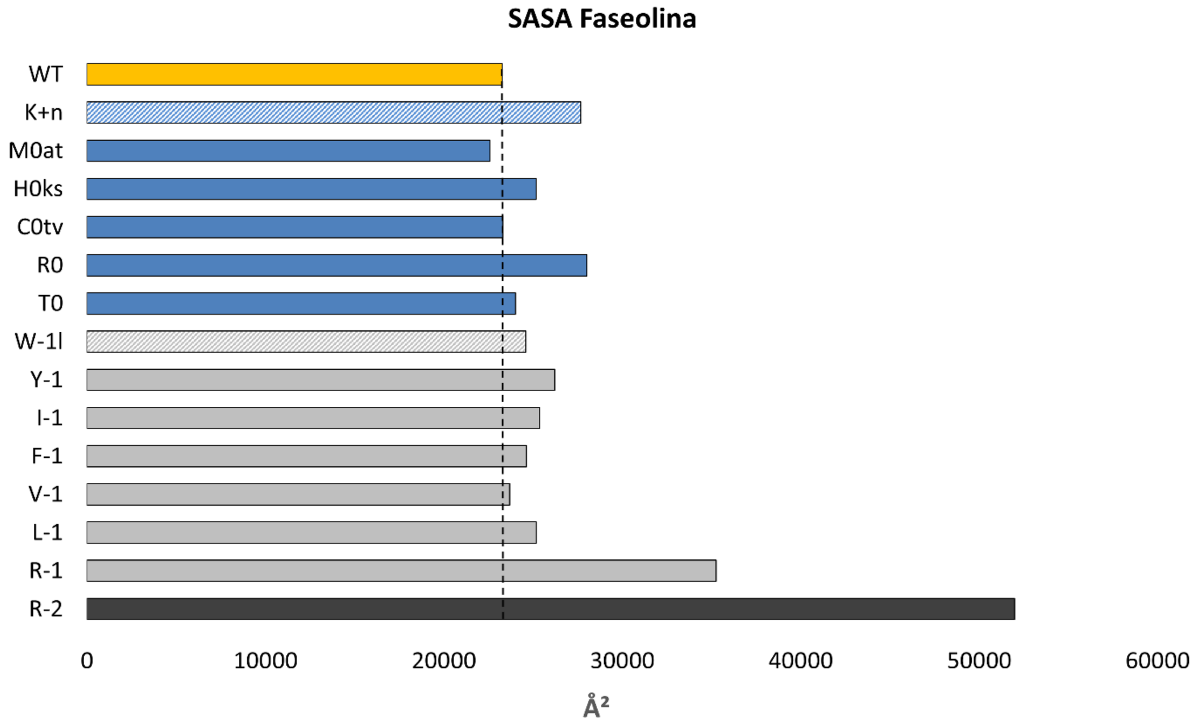

**Figura 27: Valores de SASA da Faseolina (WT) e suas mutantes. A linha tracejada indica o valor médio de RMSD da WT como referência.**
Fonte: Os autores, 2025.

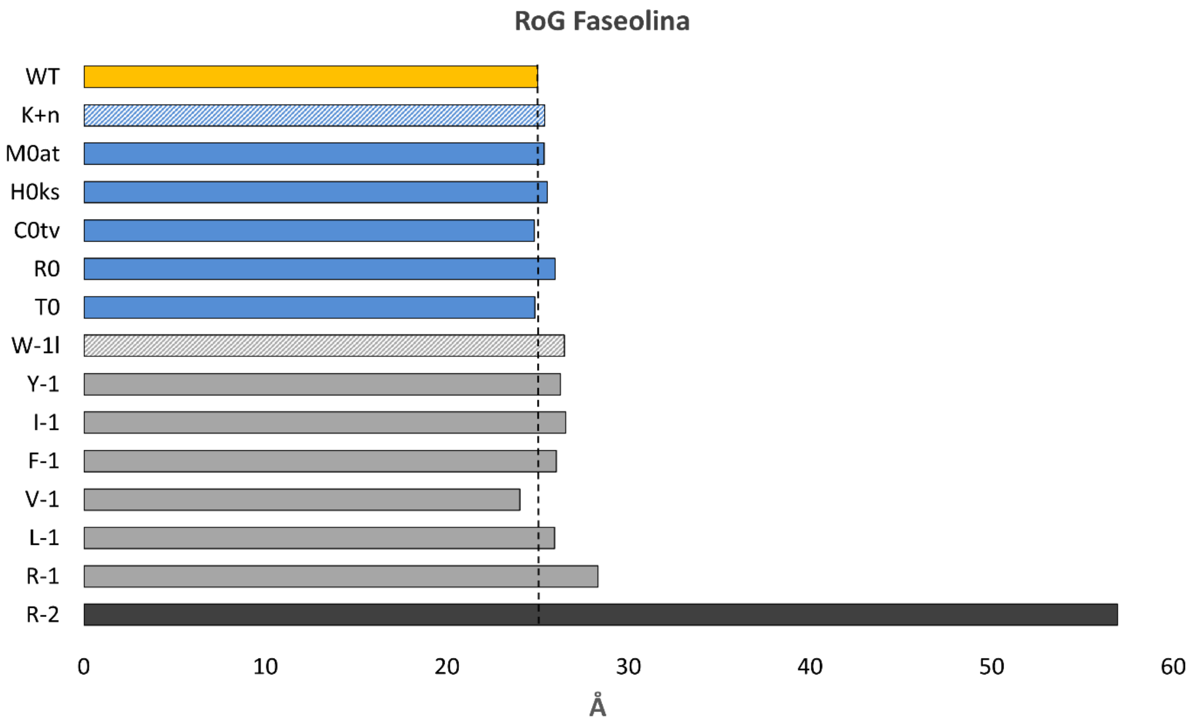

**Figura 28: Valores de raio de giro da Faseolina (WT) e suas mutantes. A linha tracejada indica o valor médio de RMSD da WT como referência.**
Fonte: Os autores, 2025.

Esse conjunto de resultados pode indicar preservação da estrutura global quando são aplicadas substituições mais conservadoras (valores de escore BLOSUM62 até -1 ou valores inferiores quando necessário) para a grande maioria das mutantes. Já as substituições por resíduos carregados (R e K) levam a aumentos um pouco mais expressivos. Esse comportamento indica que substituições mais permissivas introduzem perturbações estruturais relevantes, podendo levar a mudanças conformacionais significativas. O teste automatizado de protocolos de substituição por resíduos de aminoácidos essenciais ou semi-essenciais é apenas um dos possíveis estudos de caso onde há interesse em predição de estrutura tridimensional a partir da estrutura primária e teste da estabilidade estrutural em simulações de dinâmica molecular.

#### 6. CONSIDERAÇÕES FINAIS

O trabalho teve início com um escopo mais restrito, centrado no desenvolvimento do sistema A-DOBRA como ferramenta de modelagem estrutural. Nesse momento, o foco estava na obtenção dos modelos a partir do AlphaFold, inicialmente AlphaFold2 que demandava manipulação de bases de download lento, e posteriormente acompanhando a disponibilidade, migrou-se para AlphaFold3, sem uma preocupação aprofundada com etapas posteriores de validação. Ao longo do desenvolvimento, uma barreira relevante foi o lançamento do AlphaFold Server pela Google, que passou a oferecer uma solução acessível e eficiente para predição estrutural. Isso exigiu uma mudança de direcionamento do trabalho, que deixou de focar exclusivamente na modelagem para buscar um diferencial na análise e validação das estruturas.

Assim, o valor do projeto passou a estar na integração entre predição e dinâmica molecular, permitindo explorar aspectos que não são capturados apenas pelas métricas internas do AlphaFold. Essa ampliação permitiu uma avaliação mais completa das proteínas, considerando não apenas a estrutura prevista, mas também seu comportamento ao longo do tempo.

Nesse cenário, o pipeline AlphaUnFold se apresenta como uma solução flexível e reprodutível, que pode ser utilizada por outros pesquisadores em diferentes ambientes computacionais. Usuários podem configurar seus próprios clusters, por exemplo com Slurm, e executar todo o fluxo de forma automatizada. Além disso, há potencial para expansão do pipeline, permitindo que usuários customizem parâmetros das simulações de dinâmica molecular, definam condições específicas de execução e gerem análises e gráficos personalizados de acordo com seus objetivos de estudo.

Por fim, o sistema A-DOBRA permanece como uma interface acessível para modelagem estrutural, com potencial de evolução para integrar todo o pipeline AlphaUnFold. Essa integração pode transformar a plataforma em um ambiente mais completo, no qual o usuário não apenas gera modelos, mas também executa simulações e analisa resultados de forma centralizada. Como perspectiva futura, vislumbra-se a criação de uma interface mais interativa, com suporte à configuração de experimentos, visualização dinâmica dos resultados e geração automatizada de relatórios, ampliando ainda mais o alcance e a aplicabilidade do trabalho.

#### REFERÊNCIAS

ABRAHAM, Mark J.; MURRAY, David. **GROMACS: High performance molecular simulations through multi-level parallelism**. *SoftwareX*, v. 1–2, p. 19–25, 2020.

ABRAMSON, Josh et al. **Accurate structure prediction of biomolecular interactions with AlphaFold 3**. *Nature*, 2024. DOI: 10.1038/s41586-024-07487-w.

Agosta, F., & Cozzini, P. **A combined molecular dynamics and hydropathic interaction (HINT) approach to investigate protein flexibility: the PPARγ case study.** Molecules, 29(10), 2234. https://doi.org/10.3390/molecules29102234, 2024.

AGOSTA, Federica; COZZINI, Pietro. **A combined molecular dynamics and hydropathic interaction (HINT) approach to investigate protein flexibility: the PPARγ case study Molecules**, Basel, v. 29, n. 10, p. 2234, 2024.

Aldeghi, M., Biggin, P. C., Chackalamannil, S., Rotella, D., & Ward, S. E. **Advances in molecular simulation.** Annual Reports in Medicinal Chemistry, 51, 1-30, 2018.

Andrio, P., Hospital, A., Conejero, J., Jordá, L., Del Pino, M., Codo, L., ... & Orozco, M. **BioExcel building blocks: a software library for interoperable biomolecular simulation workflows.** Journal of Chemical Theory and Computation, 15(6), 3358–3367, 2019.

ANFINSEN, C. B. **Principles that govern the folding of protein chains**. Science, v. 181, n. 4096, p. 223-230, 1973. DOI: 10.1126/science.181.4096.223

Arsiccio, A., Metcalfe, C., Pisano, R., Raut, S., & Coxon, C. **A proximity-based in silico approach to identify redox-labile disulfide bonds: the example of FVIII.** International Journal of Molecular Sciences, 21(6), 2141, 2020.

Belghit, H., Spivak, M., Dauchez, M., Baaden, M., & Jonquet-Prevoteau, J. **From complex data to clear insights: visualizing molecular dynamics trajectories.** Computer Graphics Forum, 38(3), 291–302, 2019.

BERMAN, H. M.; WESTBROOK, J.; FENG, Z.; GILLILAND, G.; BHAT, T. N.; WEISSIG, H.; SHINDYALOV, I. N.; BOURNE, P. E. **The Protein Data Bank. Nucleic Acids Research**, v. 28, n. 1, p. 235-242, 2000.

Bernetti, M., Bertazzo, M., & Masetti, M. **Data-driven molecular dynamics: a multifaceted challenge.** Frontiers in Molecular Biosciences, 7, 1-10, 2020.

Blair, J. M. A., Webber, M. A., Bavro, V. N., Pietilä, M. K., & Piddock, L. J. V. (2024). **Molecular mechanisms of antibiotic resistance revisited**. *Nature Reviews Microbiology*, 22(4), 255. https://doi.org/10.1038/s41579-024-01014-4
BLAIR, J. M. A.; WEBBER, M. A.; BAVRO, V. N.; PIDDOCK, L. J. V. **Molecular mechanisms of antibiotic resistance revisited.** *Nature Reviews Microbiology*, v. 13, n. 1, p. 42-51, 2015.

BRÄNDÉN, Carl; TOOZE, John. **Introduction to Protein Structure**. 2. ed. New York: Garland Publishing, 1999.

Cheatham III, T. E., & Roe, D. R. **The impact of heterogeneous computing on workflows for biomolecular simulation and analysis.** Computing in Science & Engineering, 17(2), 30–39, 2015.

CHOI, J. M.; HAN, S. S.; KIM, H. S. Industrial applications of enzyme biocatalysis: Current status and future aspects. **Biotechnology Advances**, v. 33, n. 7, p. 1443–1454, 2015.

DAURA, X.; VAN GUNSTEREN, W. F.; MARK, A. E. Molecular dynamics simulations of biomolecules: from structure to function. Chichester: Wiley, 2019.

DEAN, Jeffrey; GHEMAWAT, Sanjay. **MapReduce: Simplified data processing on large clusters**. *Communications of the ACM*, v. 51, n. 1, p. 107–113, 2008.

Desai, V., Joshi, S., Soni, M., Sharma, P., & Soni, M. (2024). **Review of AlphaFold 3: Transformative Advances in Drug Design and Therapeutics.** *Cureus*, 16(7). https://doi.org/10.7759/cureus.64458

**DESAI, V.; JOSHI, S.; SONI, M.; SHARMA, P.; SONI, M.** *Review of AlphaFold 3: Transformative Advances in Drug Design and Therapeutics.* **Cureus**, v. 16, n. 7, 2024. DOI: 10.7759/cureus.64458

Di Natale, F., Bhatia, H., Carpenter, T. S., Neale, C., Kokkila-Schumacher, S., Oppelstrup, T., ... & Ingólfsson, H. I. **A massively parallel infrastructure for adaptive**

**multiscale simulations: modeling RAS initiation pathway for cancer.** Proceedings of the International Conference for High Performance Computing, Networking, Storage and Analysis (SC), 2019.

DILL, K. A.; MACCALLUM, J. L. **The protein-folding problem, 50 years on.** *Science*, v. 338, n. 6110, p. 1042-1046, 2012.

Douglas E V Pires, David B Ascher, Tom L Blundell. **mCSM: predicting the effects of mutations in proteins using graph-based signatures**, 2014 Feb 1;30(3):335-42. doi: 10.1093/bioinformatics/btt691. Epub Nov 26, 2013.

Doyle, M. T., Jimah, J. R., Dowdy, T., Ohlemacher, S. I., Larion, M., Hinshaw, J. E., & Bernstein, H. D. **Cryo-EM structures reveal multiple stages of bacterial outer membrane protein folding.** Cell, 185(7), 1143–1156, 2022.

EDDY, Sean R. **Accelerated Profile HMM Searches**. *PLoS Computational Biology*, v. 7, n. 10, e1002195, 2011.

ELSEVIER. **Protein Modeling**. *ScienceDirect Topics*. Disponível em: https://www.sciencedirect.com/topics/biochemistry-genetics-and-molecular-biology/protein-modeling. Acesso em: 8 mar. 2026.

FOWLER, Martin. *Arquiteturas de Interface Gráfica do Usuário*. 2006. Disponível em: https://martinfowler.com/pt-br/eaaDev/uiArchs.html. Acesso em: 24 Ago. 2025

GRIGORAKIS, Konstantinos; FEROUSI, Christina; TOPAKAS, Evangelos. **Protein Engineering for Industrial Biocatalysis: Principles, Approaches, and Lessons from Engineered PETases**. *Catalysts*, v. 15, n. 2, art. 147, 2025. DOI: 10.3390/catal15020147. Disponível em: https://doi.org/10.3390/catal15020147. Acesso em: 07 mar. 2026.

Hadi-Alijanvand, H., Di Paola, L., Hu, G., Leitner, D. M., Verkhivker, G. M., Sun, P., ... & Giuliani, A. **Biophysical insight into the SARS-CoV-2 spike–ACE2 interaction and its modulation by hepcidin through a multifaceted computational approach.** Journal of Physical Chemistry B, 125(6), 17024–17042, 2021.

HAUSER, A. S.; ATTWOOD, M. M.; RASK-ANDERSEN, M.; SCHIÖTH, H. B.; GLORIAM, D. E. **Trends in GPCR drug discovery: new agents, targets and technologies.** *Nature Reviews Drug Discovery*, v. 16, n. 12, p. 829-842, 2017.

HOLLINGSWORTH, S. A.; DROR, R. O. **Molecular Dynamics Simulation for All. Neuron,** 2018.

HUMBLE, Jez; FARLEY, David. **ontinuous Delivery: Reliable Software Releases through Build, Test, and Deployment Automation**. Boston: Addison-Wesley, 2011.

Jumper, J. et al. **Highly accurate protein structure prediction with AlphaFold**. Nature 596, 583–589 (2021).

Klyshko, E., Kim, J. S. H., & Rauscher, S. **LAWS: local alignment for water sites—tracking ordered water in simulations.** Journal of Chemical Theory and Computation, 14(5), 2871–2883, 2018.

Klyshko, E., Kim, J. S. H., McGough, L., Valeeva, V., Lee, E., Ranganathan, R., & Rauscher, S. **Functional protein dynamics in a crystal.** Proceedings of the National Academy of Sciences, 117(6), 3046–3054, 2020.

Krüger, A., Zimbres, F. M., Kronenberger, T., & Wrenger, C. **Molecular modeling applied to nucleic acid-based molecule development.** Molecules, 25(4), 1025, 2020.

KRYSHTAFOVYCH, A.; SCHWEDE, T.; TOPF, M.; FIDELIS, K.; MOULT, J. **Critical assessment of methods of protein structure prediction (CASP)—Round XIV.** *Proteins: Structure, Function, and Bioinformatics*, v. 89, n. 12, p. 1607-1617, 2021.

Kunszt, P., Malmström, L., Fantini, N., Sudholt, W., Lautenschlager, M., Reifler, R., & Ruckstuhl, S. **Accelerating 3D protein modeling using cloud computing: using Rosetta as a service on the IBM SmartCloud.** IEEE International Conference on Cloud Computing, 2012.

KURTZER, Gregory M.; SOCHAT, Vanessa; BAUER, Michael W. **Singularity: Scientific containers for mobility of computers**. *PLoS ONE*, v. 12, n. 5, e0177459, 2017

LEACH, A. R. **Molecular Modelling: Principles and Applications.** 2nd ed. Harlow: Pearson, 2001.

Lee, H., Turilli, M., Jha, S., Bhowmik, D., Ma, H., & Ramanathan, A. **DeepDriveMD: deep-learning driven adaptive molecular simulations for protein folding.** IEEE/ACM Workshop on Machine Learning in HPC Environments, 2019.

Liao, G., Luo, L., & Che, N. **Hybrid parallel bidirectional sieve based on SMP cluster.** Journal of Parallel and Distributed Computing, 74(1), 1920–1930, 2014.

**MAMBA DEVELOPMENT TEAM**. *Micromamba documentation*. Disponível em: https://mamba.readthedocs.io/en/latest/user_guide/micromamba.html. Acesso em: 01 dez. 2025.

MERKEL, Dirk. **Docker: Lightweight Linux containers for consistent development and deployment**. *Linux Journal*, n. 239, 2014.

Michaleas, A., & Ricke, D. O. **Scalable and portable pipelines for predicting 3D protein structures on standalone and HPC systems.** Future Generation Computer Systems, 107, 498–509, 2020.

Neamtu, A., Serban, D. N., Barritt, G. J., Isac, D. L., Vasiliu, T., Laaksonen, A., & Serban, I. L. **Molecular dynamics simulations reveal the hidden EF-hand of EF-SAM as a possible key thermal sensor for STIM1 activation by temperature.** Scientific Reports, 10, 1–12, 2020.

NELSON, D. L.; COX, M. M. Princípios de Bioquímica de Lehninger. 8. ed. Porto Alegre: Artmed, 2021.

NEWMAN, Sam. **Building Microservices: Designing Fine-Grained Systems**. 2. ed. Sebastopol: O'Reilly Media, 2021.

NGOC, L. T. N.; MOON, J.-Y.; LEE, Y.-C**. Insights into Bioactive Peptides in Cosmetics**. Cosmetics, v. 10, n. 4, p. 111, 2023. Disponível em: https://www.mdpi.com/2079-9284/10/4/111. Acesso em: 03 Set 2025. DOI: https://doi.org/10.3390/cosmetics10040111.

**NICKOLLS, J.; DALLY, W. J.** The GPU computing era. **IEEE Micro**, v. 30, n. 2, p. 56-69, 2010. DOI: 10.1109/MM.2010.41.

Nithin, C., Fornari, R. P., Pilla, S. P., Wroblewski, K., Zalewski, M., Madaj, R., ... & Kmiecik, S. **Exploring protein functions from structural flexibility using CABS-flex modeling.** Bioinformatics, 35(4), 694–695, 2019.

Ossyra, J. R., Sedova, A., Baker, M. B., & Smith, J. C. **Highly interactive, steered scientific workflows on HPC systems: optimizing design solutions.** Concurrency and Computation: Practice and Experience, 32(5), 2020.

Schlick, T., Portillo-Ledesma, S., Myers, C. G., Beljak, L., Chen, J., Dakhel, S., ... & Xue, E. **Biomolecular modeling and simulation: a prospering multidisciplinary field.** Annual Review of Biophysics, 50, 267–301, 2021.

Senior, A. W., Evans, R., Jumper, J., Kirkpatrick, J., Sifre, L., Green, T., ... & Hassabis, D. (2020). **Improved protein structure prediction using potentials from deep learning.** *Nature*, 577(7792), 706-710.

SILVA, Gustavo Dias da. **Utilização do algoritmo Apriori para traçar o perfil sociodemográfico do homem brasileiro com câncer de próstata**. 2022. Dissertação (Mestrado em Ciência e Tecnologia em Saúde) – Universidade Estadual da Paraíba, Campus de Campina Grande, Pró-Reitoria de Pós-Graduação e Pesquisa, Programa de Pós-Graduação em Ciência e Tecnologia em Saúde, Campina Grande, 2022.

**ŚLEDŹ; CAFLISCH (2018)** — *Protein structure-based drug design: from docking to molecular dynamics* (Current Opinion in Structural Biology).

STEINEGGER, Martin; SÖDING, Johannes. **MMseqs2 enables sensitive protein sequence searching for the analysis of massive data sets**. *Nature Biotechnology*, v. 35, p. 1026–1028, 2017.

Teixeira, J. M. C., Liu, Z. H., Namini, A., Li, J., Vernon, R. M., Krzeminski, M., ... & Forman-Kay, J. D. **IDPConformerGenerator: a flexible software suite for sampling the conformational space of disordered protein states.** Journal of Chemical Theory and Computation, 17(9), 5985–6003, 2021.

Tunstall, T., Portelli, S., Phelan, J., Clark, T. G., Ascher, D. B., & Furnham, N. **Combining structure and genomics to understand antimicrobial resistance.** Nature Reviews Genetics, 23, 451–465, 2022.

VAN DEN BEDEM, Henry; FRASER, James S. **Integrative, dynamic structural biology at atomic resolution: it's about time**. *Nature Methods*, v. 12, p. 307–318, 2015.

Wehlin, A., Cornaciu, I., Marquez, J. A., Perrakis, A., & von Castelmur, E. **Crystal structure of the phospholipase A and acyltransferase 4 (PLAAT4) catalytic domain.** Journal of Biological Chemistry, 295(6), 1613–1623, 2020.

WILSON, Daniel N. **Ribosome-targeting antibiotics and mechanisms of bacterial resistance**. *Nature Reviews Microbiology*, v. 12, n. 1, p. 35–48, 2014.

WORLD WIDE WEB CONSORTIUM (W3C). *Arquitetura de Web Services*. Recomendação W3C, 2004. Tradução livre do original *Web Services Architecture*. Disponível em: https://www.w3.org/TR/ws-arch/. Acesso em: [25-10-2025].

**YOO, A. B.; JETTE, M. A.; GRONDONA, M.** SLURM: Simple Linux Utility for Resource Management. In: **Workshop on Job Scheduling Strategies for Parallel Processing**. Springer, Berlin, Heidelberg, 2003. p. 44-60. DOI: 10.1007/10968987_3.

Ziemianowicz, D. S., Saltzberg, D., Pells, T., Crowder, D. A., Schrader, C., Hepburn, M., ... & Schriemer, D. C. **IMProv: a resource for cross-link-driven structure modeling that accommodates protein dynamics.** Nucleic Acids Research, 49(W1), W350–W356, 2021.

#### APÊNDICES

##### APÊNDICE I - REVISÃO SISTEMÁTICA

##### DESCRIÇÃO DO PROBLEMA

Com os avanços nas tecnologias muitos problemas da comunidade científica e da própria sociedade foram sendo melhor investigados, entre eles está a modelagem de proteínas na área de bioinformática. Essa tornou-se uma prática cada vez mais comum nos grandes laboratórios ao redor do mundo, bem como em pesquisas científicas. Todavia, é uma atividade que exige, além dos conhecimentos técnicos, alto poder de processamento computacional, o que a torna muitas vezes uma tarefa inacessível para muitos usuários e cientistas.

A bioinformática estrutural, em particular a modelagem e a dinâmica de proteínas são procedimentos relevantes para a comunidade científica, uma vez que servem para auxiliar em nichos importantes da sociedade, como em descobertas de novos medicamentos e vacinas. Existem softwares que realizam esses trabalhos, mas como supracitado existem desafios para poder utilizá-los. É neste contexto que vamos focar em nosso trabalho, prover um HPC - *High Processing Computing* integrado a uma interface web que possa prover facilidade, agilidade e rentabilidade à comunidade de bioinformatas e cientistas no geral para realizar seus experimentos *in silico*.

#### OBJETIVOS

##### Objetivo Geral

Levantar o conteúdo bibliográfico para compreensão do tema e das lacunas sobre o desenvolvimento de sistema web integrado a um HPC provendo um pipeline otimizado para aplicações de Bioinformática Estrutural.

##### Objetivos Específicos

- Analisar publicações científicas para delimitar as principais ferramentas que se propõem a realizar trabalhos similares;
- Contribuir com o trabalho de bioinformatas na execução de atividades e experimentos *in silico* que demandam altos custos financeiros e computacionais, com a o auxílio de materiais bibliográficos;

#### QUESTÕES DE PESQUISA

Ao cumprirmos os objetivos descritos na seção anterior teremos embasamento teórico para responder os seguintes questionamentos:

1. Quais as formas de aplicação de HPC (*High Processing Computing)* na modelagem de proteína no estado da arte?
2. Quais as aplicações inovadoras que versem sobre modelagem ou dinâmica de proteínas?
3. Quais as aplicações de software livre na modelagem de proteína?
4. Quais os critérios de validação de solução de modelagem e dinâmica molecular de proteínas estão sendo aplicados no estado da arte?

As perguntas apresentam o valor fundamental no desenvolvimento do entendimento da aplicação de solução HPC na modelagem e dinâmica de proteínas. Isso, sendo alcançado através primeiramente da resposta da primeira pergunta, onde objetivamos entender a aplicação HPC na modelagem de proteína, em segundo ponto na segunda questão buscamos elencar as aplicações modernas de modelagem e dinâmica de proteínas, na terceira buscaremos solução de software livre de modelagem de proteína e por último a quarta questão buscaremos os critérios que são utilizados pela academia para validar uma solução de modelagem de proteína.

**Estratégias de busca**

Nessa seção abordaremos todos os meios utilizados para realização das buscas. A definição das fontes de pesquisa para o levantamento bibliográfico, onde teremos a escolha das fontes de publicações, as definições das palavras-chave, os termos técnicos principais na área, a definição da string de busca que será aplicada nas buscas nas bases.

**Fontes de pesquisa**

Foram selecionadas as bibliotecas digitais IEEE XPLORE, ACM Library, ScienceDirect, PubMed e Scopus para a realização das consultadas em suas bases.

**Palavras-Chave**

Através das questões de pesquisa definidas na subseção 2.2, podemos retirar as seguintes palavras-chave: High Processing Computing (Computação de alto processamento), protein modeling (modelagem de proteína), protein dynamics (dinâmica de proteína), Workflow Description Language ( linguagem de descrição de fluxo), validation criteria (critérios de validação).

**String de busca**

Utilizando-se das palavras-chave anteriores e com a finalidade de remover os dados supérfluos e incluir os que deem embasamento para responder as questões de buscas a *string* de busca ficou de acordo com o Quadro 1, a mesma foi construída utilizando os sinônimos dos termos chaves para garantir que nos resultados encontrássemos trabalhos de acordo com o tema mesmo sendo escrito com termos semelhantes.

**Quadro 1. String de busca em inglês**

| **String de busca** |
| --- |
| Sugestão: "HPC" AND ("Protein Folding" OR "Protein dynamics") AND ("Workflow" OR "Pipeline" OR Workloader) |

Fonte: Os Autores, 2024.

Porém, para a consulta na base scielo, que é uma base que adota a língua portuguesa esta string de busca ficou escrita de acordo com o Quadro 2.

**Língua**

A língua adotada para escolha na busca é o inglês por se tratar da mais adotada como língua vernácula.

**Critérios de seleção e exclusão**

Para realizar o processo de seleção das publicações que serão analisadas na revisão, serão considerados os critérios de inclusão e de exclusão dispostos a seguir.

Os critérios de inclusão são:

- O resultado deve estar no idioma inglês;
- O resultado deve estar disponível integralmente na web;
- O resultado deve conter no título e no resumo alguma relação com o tema deste trabalho.

Os critérios de exclusão são:

- O resultado ter publicação superior a cinco anos;
- O resultado ainda se encontra na fase de desenvolvimento;
- O resultado obter o nível inferior ao nível low na avaliação de qualidade.

**Procedimento da seleção**

Para realização do processo de seleção dos resultados obtidos utilizamos a ferramenta Start (State of the Art through Systematic Review), software desenvolvido por um laboratório (LaPES) da Universidade Federal de São Carlos do Estado de São Paulo no Brasil, para gerenciar todas as referências e citações encontradas na busca.

No início realizamos a busca em cada fonte de pesquisa, depois o seu resultado foi exportado para um arquivo BibTex[1] que utilizado na ferramenta para o gerenciamento das citações. Depois de importado os arquivos das pesquisas o Start lista todos os resultados, e primeiramente questiona se desejamos classificar os trabalhos duplicados automaticamente ou de forma manual.

Diante dos resultados das fontes, o processo de seleção ocorrerá em duas fases, onde a primeira será referente ao levantamento dos trabalhos que obedecem aos critérios de inclusão, caso o resultado não obedeça pelo menos um item o mesmo será excluído. Na segunda fase haverá a consulta aos títulos e resumos para constatar se algum trabalho ainda atende aos critérios de exclusão, caso satisfaça pelo menos um o trabalho será rejeitado.

Logo após a realização dos passos de seleção restaram apenas os trabalhos que serão utilizados para extração dos dados relevantes para o trabalho em questão, que terão os seus passos melhor descritos nas próximas seções

**Análise de Qualidade**

Com a finalidade de considerar trabalhos com um teor de qualidade considerável na extração dos dados, o último critério de exclusão menciona uma avaliação de qualidade, que conterá fatores que darão esse teor de qualidade.

A avaliação de qualidade tem o objetivo de enquadrar o trabalho em alguns níveis de prioridade para a leitura do trabalho, que variarão de acordo com a sua elaboração. Os níveis de avaliação serão quatro, oferecidos pela ferramenta Start, sendo eles: *very low*, *low*, *high* e *very high*, que para cada um consideramos a sua forma com que trata o tema proposto nesse trabalho. Para esta etapa, os níveis da avaliação se encontram de acordo com o Quadro 3.

[1] BibTex: é um arquivo que normalmente é utilizado em conjunto com o sistema tipográfico LaTex, porém se tornou bastante utilizado por outras ferramentas no controle da bibliografia.

**Quadro 3. Avaliação de qualidade**

| **Níveis** | **Critérios** |
|---|---|
| Very low | O trabalho faz menção a apenas modelagem de proteína |
| Low | O trabalho faz menção à modelagem e a dinâmica de proteína |
| High | O trabalho faz menção à modelagem e a dinâmica de proteína |
| Very High | O trabalho faz menção à modelagem e a dinâmica de proteína com o uso de Workloader/Workflow e aplicação de HPC. |

Fonte: Os autores, 2024.

O processo de qualificação ocorrerá com a leitura das partes de introdução e conclusão dos trabalhos, para assim ter o nível de tratamento do estudo com relação ao desenvolvimento de sistema web integrado a um HPC provendo um pipeline otimizado para a modelagem e a dinâmica de proteínas. Ao fim dessa etapa os trabalhos, que restarem, serão repassados para a etapa de extração de dados.

**Extração dos Resultados**

Para a realização da extração dos dados realizou-se a sua catalogação dos trabalhos na ferramenta como pode ser visto na ficha presente na Figura a seguir, onde dentre todos os dados considerados para fins de extração temos: autores, título, palavras-chave, ano de publicação, base de indexação, técnicas de modelagem e dinâmica de proteína, ferramentas de disponibilização de resultados da modelagem, objetivo do estudo. Estes itens serão catalogados na guia *Data Extraction Form*, como pode ser visto na Figura abaixo.

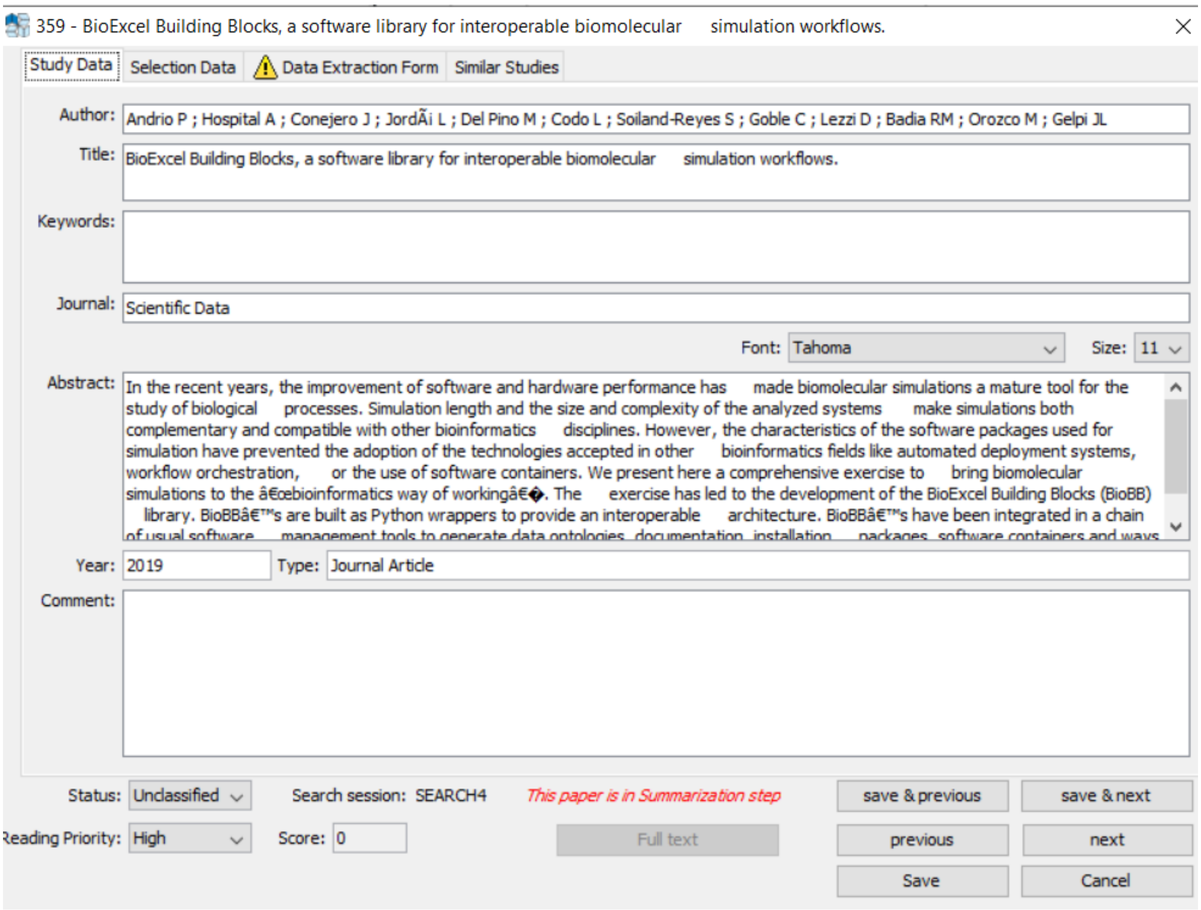

**Figura 29 - Catálogo dos campos para extração do software Start.**
Fonte: Adaptado de SILVA, 2022.

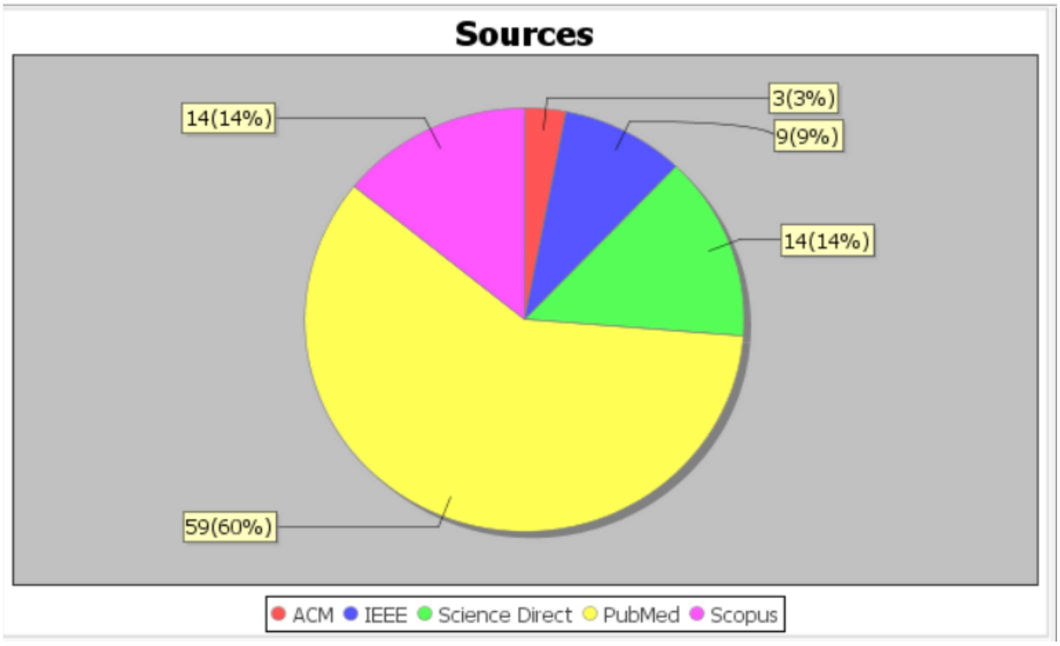

**Figura 30: Distribuição dos trabalhos retornados na busca por cada portal.**
Fonte: Start v.2.3.2

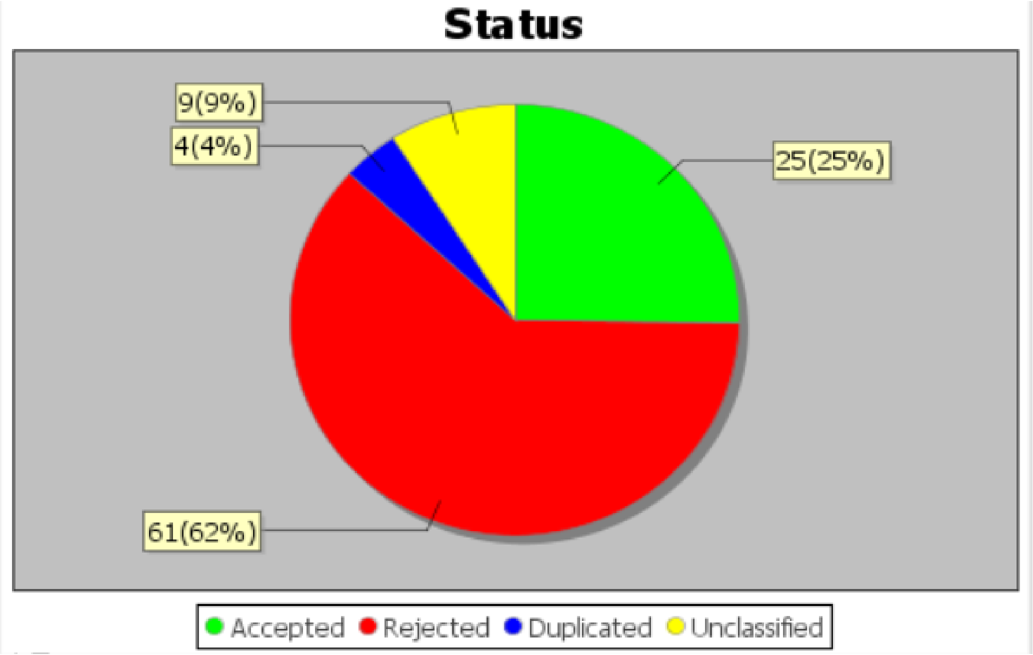

**Figura 31: Resultado da etapa de inclusão dos trabalhos.**
Fonte: Start v.2.3.2

### APÊNDICE II: TELAS INICIAIS DO SISTEMA WEB

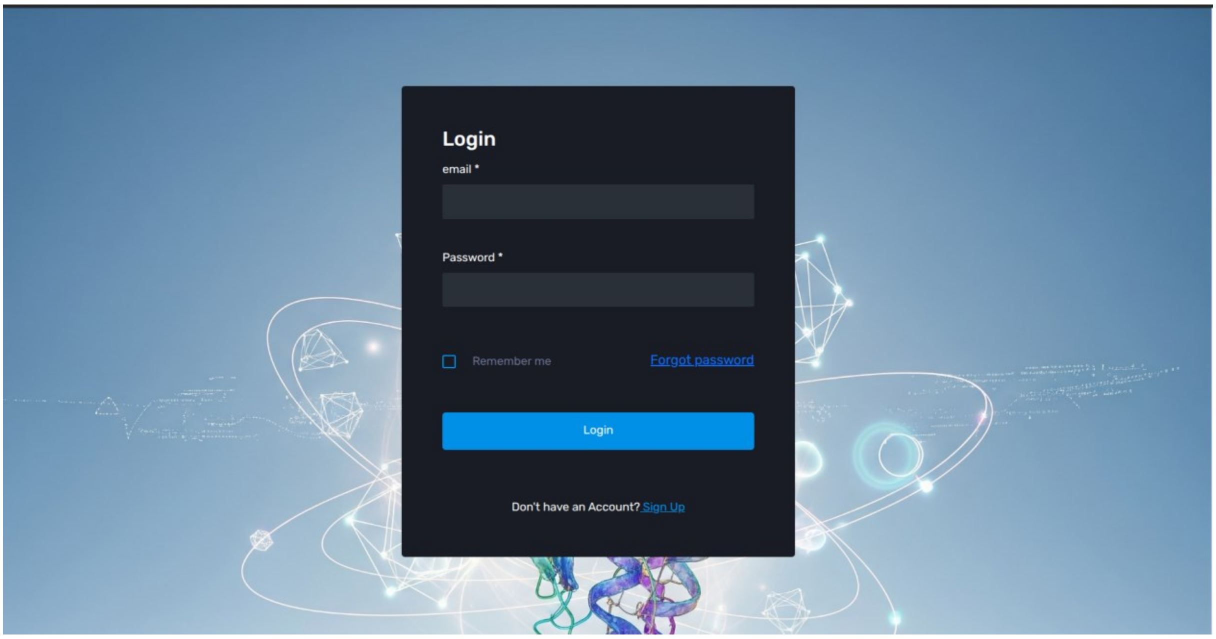

**Figura 32: Tela Inicial do Sistema A-DOBRA**
Fonte: Os Autores, 2025.

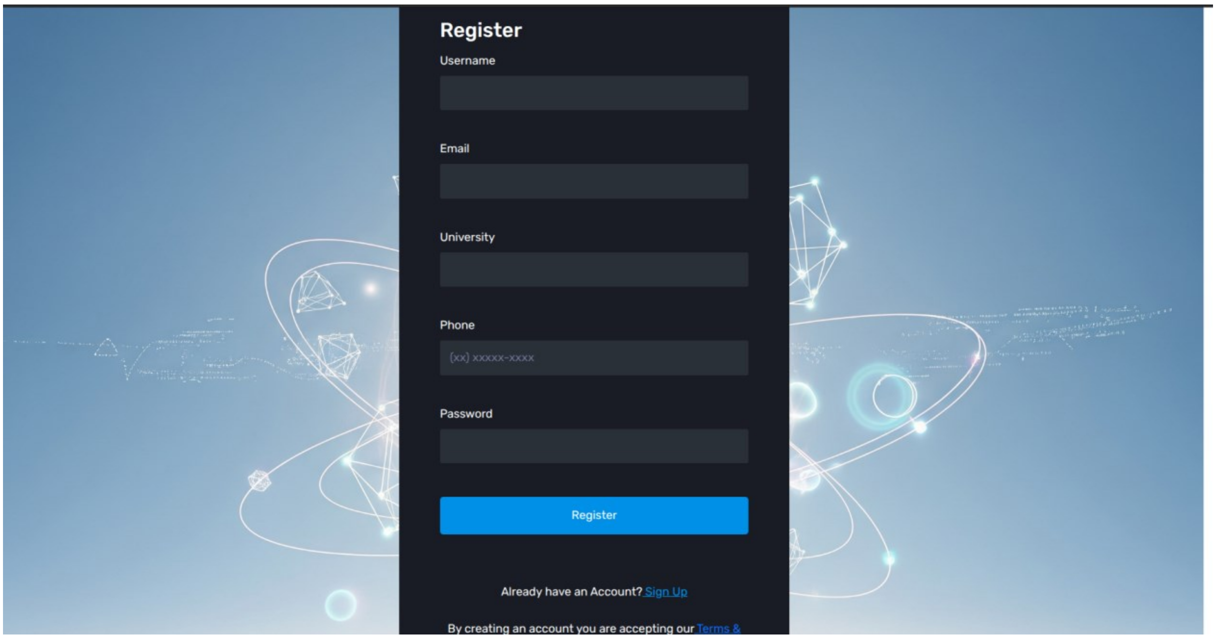

**Figura 33: Tela de Registro do Sistema A-DOBRA**
Fonte: Os Autores, 2025.

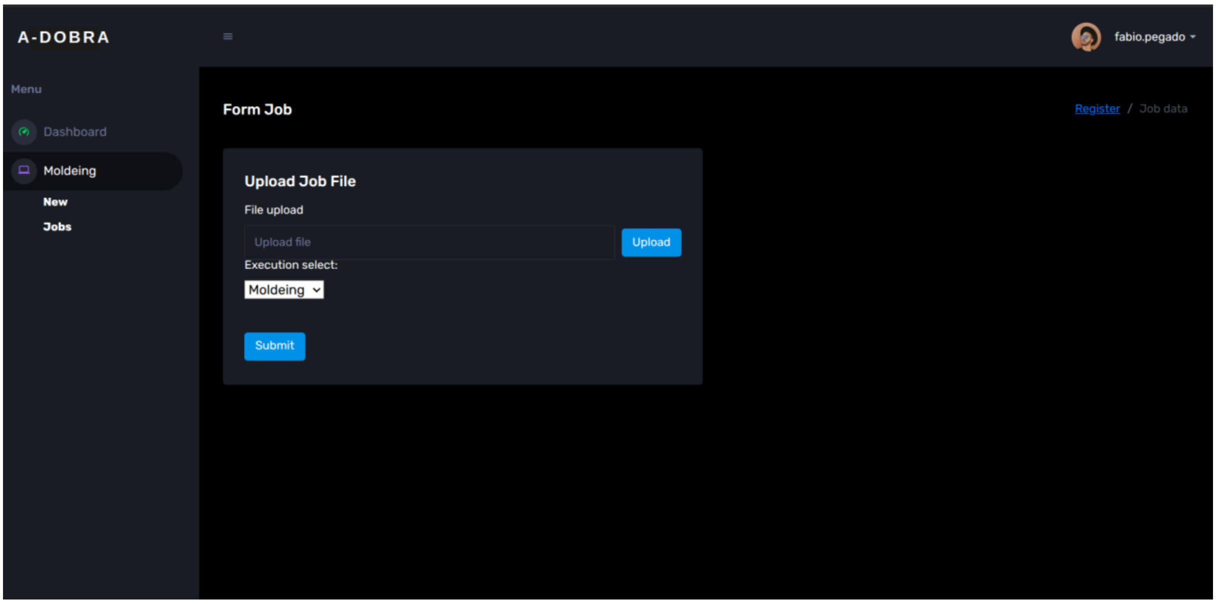

**Figura 34: Tela de Envios de *Jobs* do Sistema A-DOBRA**
Fonte: Os Autores, 2025.

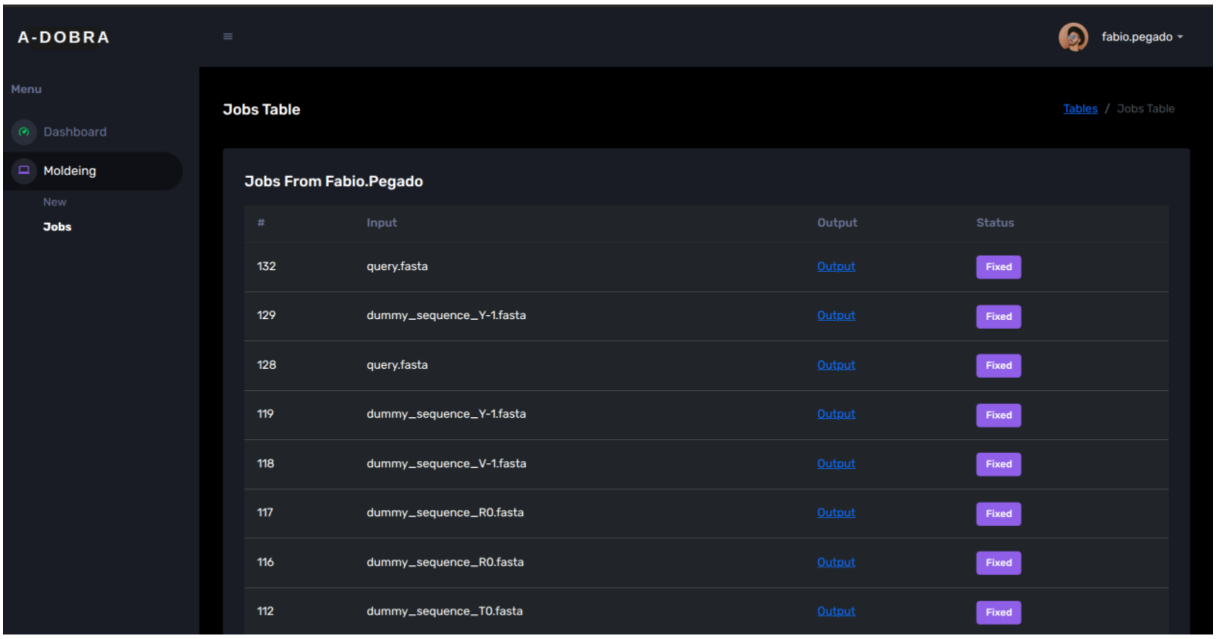

**Figura 35: Tela de Lista de *Jobs* do Sistema A-DOBRA**
Fonte: Os Autores, 2025.
